# Protein evolvability under rewired genetic codes

**DOI:** 10.1101/2023.06.20.545706

**Authors:** Hana Rozhoňová, Carlos Martí-Gómez, David M. McCandlish, Joshua L. Payne

**Affiliations:** Institute of Integrative Biology, ETH Zürich, Zürich, Switzerland; Swiss Institute of Bioinformatics, Lausanne, Switzerland; Simons Center for Quantitative Biology, Cold Spring Harbor Laboratory, Cold Spring Harbor, NY, USA

## Abstract

The standard genetic code defines the rules of translation for nearly every life form on Earth. It also determines the amino acid changes accessible via single-nucleotide mutations, thus influencing protein evolvability — the ability of mutation to bring forth adaptive variation in protein function. One of the most striking features of the standard genetic code is its robustness to mutation, yet it remains an open question whether this robustness facilitates or frustrates protein evolvability. To answer this question, we use data from massively-parallel sequence-to-function assays to construct and analyze empirical adaptive landscapes under hundreds of thousands of rewired genetic codes, including those of codon compression schemes relevant to protein engineering and synthetic biology. We find that robust genetic codes tend to enhance protein evolvability by rendering smooth adaptive landscapes with few peaks, which are readily accessible from throughout sequence space. By constructing low-dimensional visualizations of these landscapes, which each comprise more than 16 million mRNA sequences, we demonstrate that alternative genetic codes can radically alter the topological features of the network of high-fitness genotypes. Whereas the genetic codes that optimize evolvability depend to some extent on the detailed relationship between amino acid sequence and protein function, we also uncover general design principles for engineering non-standard genetic codes for enhanced and diminished evolvability, which may facilitate directed protein evolution experiments and the biocontainment of synthetic organisms, respectively. Our findings demonstrate that the standard genetic code, a critical and near-universal cellular information processing system, not only mitigates replication and translation errors as compared to most alternative genetic codes, but also facilitates predictable and directional adaptive evolution by enabling evolving populations to readily find mutational paths to adaptation.

## 1 Introduction

Proteins are the workhorses of the cell. They are the building blocks of cellular infrastructure, they transport molecules, regulate gene expression, and catalyze essential biochemical reactions. How do such protein functions evolve? The classic metaphor of the adaptive landscape is helpful to conceptualize this process (Wright, 1932). An adaptive landscape is a mapping from genotype space onto fitness or some related quantitative phenotype, which defines the “elevation” of each coordinate in this space. For proteins, genotype space comprises the set of all possible amino acid sequences of a given length (Maynard Smith, 1970) and the quantitative phenotypes of these sequences include catalytic activity, folding stability, and binding affinity. The evolution of protein function can then be viewed as a hill-climbing process in such a landscape, in which mutation and natural selection tend to drive evolving populations toward adaptive peaks of improved functionality (Romero and Arnold, 2009).

Central to this process is evolvability — the ability of mutation to bring forth adaptive phenotypic variation (Payne and Wagner, 2019; Pigliucci, 2008). For short-term, one-step adaptation, evolvability depends on the immediate mutational neighborhood of a protein sequence (Fig. 1A). That is, it depends on the amount of adaptive phenotypic variation accessible via point mutation. For longer-term, multi-step adaptation, evolvability depends on the topography of the adaptive landscape. A smooth single-peaked landscape facilitates evolvability, because mutation can easily bring forth adaptive phenotypic variation from anywhere in the landscape, except atop the global peak; in contrast, a rugged landscape diminishes evolvability, because its adaptive valleys often preclude the generation of adaptive phenotypic variation (Kauffman and Levin, 1987; de Visser and Krug, 2014; Payne and Wagner, 2019) (Fig. 1B). Landscape ruggedness also influences the predictability of evolution: while in a smooth single-peaked landscape, an evolving population will converge on the global peak regardless of its starting point, in a rugged, multi-peaked landscape, the population may become trapped on any one of the landscape’s local peaks, depending on starting conditions and the order in which adaptive mutations go to fixation (de Visser and Krug, 2014; Starr et al., 2017; Papkou et al., 2023).

**Figure 1:**
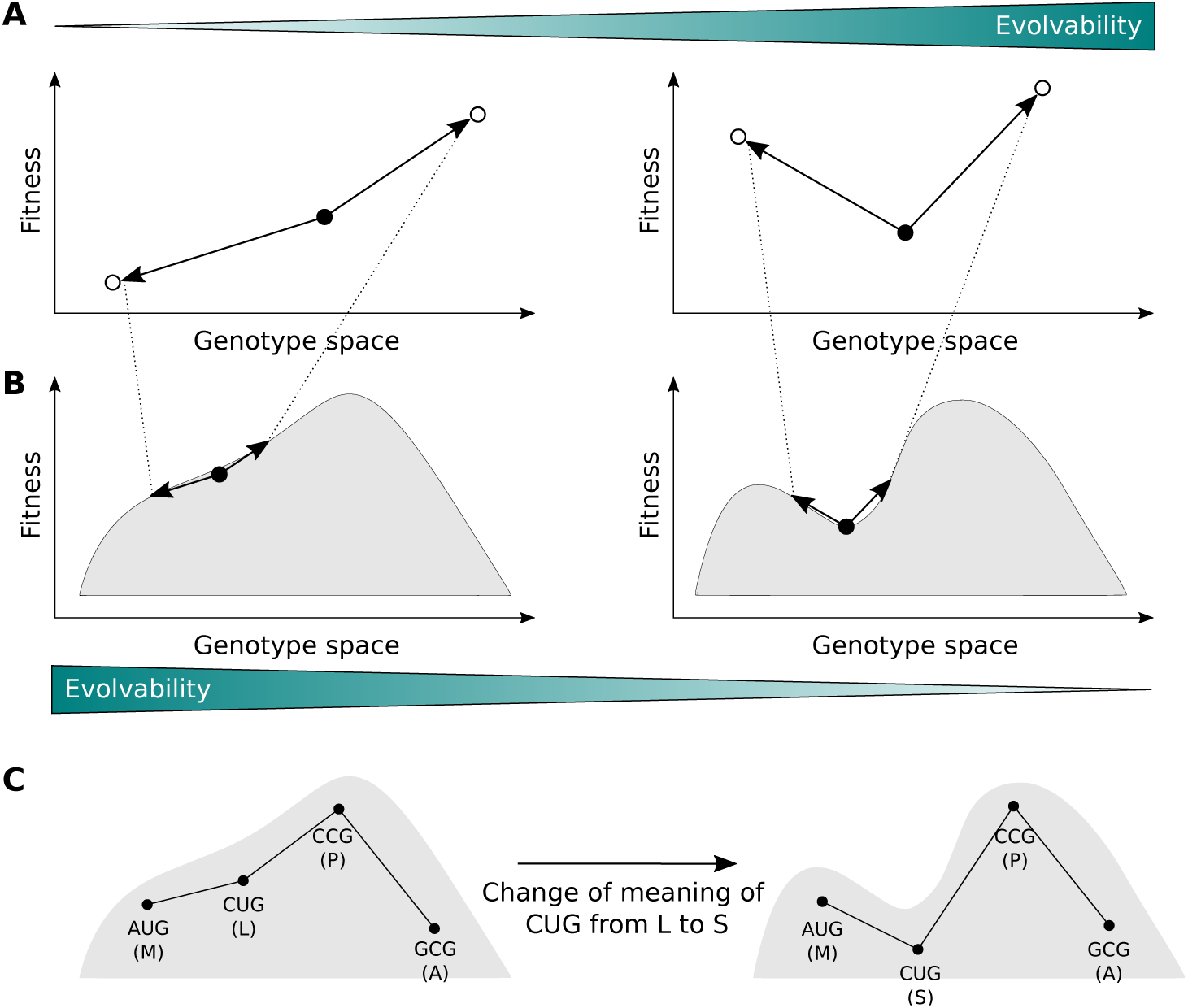
Evolvability and adaptive landscapes. (A) In one-step adaptation, evolvability depends on the amount of adaptive phenotypic variation accessible via point mutation. Therefore, the genotype shown with a filled circle in the right panel is more evolvable than the one shown in the left panel. (B) Zooming out and considering multi-step adaptation, landscape topography becomes important. Smoother landscapes promote evolvability (left panel), whereas rugged landscapes hinder evolvability (right panel), because an evolving population is more likely to be trapped on a local optimum. (C) Landscape topography is influenced by the genetic code. As a toy model, a sequence consisting of a single codon is shown. Under the standard genetic code, there is a single peak, which is also a global optimum (left panel). If the meaning of the CUG codon is changed from leucine to serine (as is the case in some yeast species (Krassowski et al., 2018)), an adaptive valley is formed (right panel). The population now cannot leave the local optimum consisting of the AUG codon without crossing a maladaptive valley.

What determines whether a protein’s adaptive landscape is smooth or rugged? One primary factor is the standard genetic code, which defines the rules of protein synthesis for nearly every life form on Earth (Knight et al., 2001). The importance of the standard genetic code arises because it determines which amino acid changes are accessible via alteration of a single nucleotide. For example, point mutations to the CUG codon can change the amino acid leucine to methionine (AUG), valine (GUG), proline (CCG), glutamine (CAG), and arginine (CGG), but not to any other of the remaining 14 amino acids. The standard genetic code thus defines the wiring diagram of protein space (Maynard Smith, 1970), determining which mutational paths to adaptation are closed or open (Fig. 1C).

The structure, history, and evolutionary implications of the standard genetic code have fascinated scientists for decades (Woese, 1965; Crick, 1968; Knight et al., 1999; Koonin and Novozhilov, 2009, 2017). Given the nearly infinite space of alternatives, why did life converge on the standard genetic code? What makes it so special? Answers to this question are typically based on comparisons of the properties of the standard genetic code to those of hypothetical, alternative codes (Haig and Hurst, 1991; Freeland and Hurst, 1998), of which there are many (Freeland et al., 2003). Even if one maintains the degeneracy of the standard code, but simply randomizes which amino acids are assigned to which codon blocks, there are 20! *≈* 10^18^ possible rewirings. By sampling a large number of such rewired codes, one can ask whether a given quantitative property of the standard genetic code has a value higher or lower than expected by chance. For example, using a measure of so-called “error tolerance” based on how well point mutations preserve polar requirement (a measure of hydrophilicity), and taking into consideration mutation bias toward transitions relative to transversions, Freeland and Hurst (1998) showed that only one in a million rewired codes preserves the hydrophilicity of amino acids to a greater extent than the standard genetic code. The standard genetic code is thus highly robust to error, in that point mutations and translation errors tend to cause minor changes to the physicochemical properties of amino acids.

What are the implications of code robustness for protein evolvability? By definition, a robust genetic code limits the amount of phenotypic variation that point mutations can cause (e.g., in terms of amino acid hydrophilicity). However, opinions differ on whether this hinders or facilitates evolvability. Inspired by Fisher’s Geometric model (Fisher, 1930), early theoretical work argues that code robustness may facilitate protein evolvability exactly because it minimizes the effects of mutations, thus increasing the probability that mutations will be adaptive (Freeland, 2002). Indeed, by analyzing the fitness effects of point mutations to the antibiotic resistance gene TEM-1 *β*-lactamase and two influenza hemagglutinin inhibitor genes, it has been shown that missense mutations are enriched for adaptive amino acid changes, relative to amino acid changes that require multiple point mutations (Firnberg and Ostermeier, 2013; Firnberg et al., 2014). In contrast, more recent theoretical work (Pines et al., 2017), motivated by advances in synthetic biology (Ostrov et al., 2016; de la Torre and Chin, 2021; Zürcher et al., 2022; Fredens et al., 2019; Chin, 2014; Liu and Schultz, 2010), argues that protein evolvability can be enhanced by reducing code robustness, because by doing so one can increase the number and diversity of amino acids accessible via point mutation to any codon.

Whether code robustness hinders or facilitates protein evolvability therefore remains an open question. In answering this question, we are faced with both conceptual and scientific challenges. Conceptually, the challenge pertains to the timescale of adaptation. If the timescale is short, for example where adaptation proceeds via a single mutation, then reducing code robustness likely enhances protein evolvability, because it increases the number and diversity of amino acids in the mutational neighborhood of any codon (Pines et al., 2017). Over longer evolutionary timescales, where adaptation proceeds via a sequence of mutations, protein evolvability depends on adaptive landscape topography, leading us to the scientific challenge of constructing realistic adaptive landscapes under the standard genetic code, as well as under a large number of rewired codes.

Some steps in this direction have been taken (Firnberg and Ostermeier, 2013; Firnberg et al., 2014; Tripathi and Deem, 2018; Zhu and Freeland, 2006; Aita et al., 2000), but these studies suffer from at least one of two key limitations. The first is a focus on how missense mutations change the physicochemical properties of amino acids (Zhu and Freeland, 2006; Haig and Hurst, 1991; Freeland and Hurst, 1998), rather than how missense mutations change the phenotype of a protein (e.g., its stability or catalytic activity) or the corresponding fitness of an organism. This is a major limitation because it is not currently possible to predict the phenotypic effects of missense mutations based on the amino acid changes they cause (Yampolsky and Stoltzfus, 2005). The second limitation is a lack of suitable data, with studies relying on a purely theoretical model of landscape topography (Zhu and Freeland, 2006), a categorical, rather than quantitative, protein phenotype (Tripathi and Deem, 2018), an incomplete fitness landscape (Firnberg and Ostermeier, 2013), or assumptions of additivity regarding the combined effects of mutations (Aita et al., 2000). These are major limitations because categorical phenotypes (e.g., the protein binds a ligand or not) do not provide quantitative information about phenotypic variation and are therefore not amenable to landscape-based analyses, and because key assumptions of theoretical models of landscape topography, particularly additivity, are commonly violated (Wu et al., 2016; de Visser and Krug, 2014; Weinreich et al., 2006). We therefore do not know how the structure of a genetic code, standard or otherwise, influences the evolvability of proteins beyond one-step adaptation. This is an important knowledge gap, because protein evolution often proceeds via a sequence of adaptive mutations that improve protein function, as evidenced by comparisons of orthologous sequences (Karageorgi et al., 2019; Natarajan et al., 2018) and directed protein evolution experiments (Fasan et al., 2008; Goldsmith and Tawfik, 2017). Moreover, given the increasing interest in engineering non-standard genetic codes (Ostrov et al., 2016; de la Torre and Chin, 2021; Zürcher et al., 2022; Fredens et al., 2019; Chin, 2014; Liu and Schultz, 2010), it is desirable to deduce design principles for engineering genetic codes with reduced or enhanced evolvability, as these might be used to form a ‘genetic firewall’ (Calles et al., 2019) or accelerate directed evolution (Pines et al., 2017), respectively.

Here, we overcome the limitations of prior studies using experimental data from massively-parallel sequence-to-function assays (Kinney and McCandlish, 2019). In particular, we use combinatorially complete data, which provide a quantitative characterization of protein phenotype for all possible combinations of 20*^L^* amino acid sequence variants at a small number *L* of protein sites (Wu et al., 2016; Lite et al., 2020; Hartman et al., 2019). These data facilitate the construction of complete adaptive landscapes without assumptions regarding the combined effects of individual mutations (e.g., additvity). Importantly, the combinatorially complete nature of these data allow us to construct such landscapes under arbitrary genetic codes. The reason is that, no matter which code we use, we are guaranteed that each of the 4^3*L*^ possible mRNA sequences can be computationally translated into an amino acid sequence with an experimentally assayed phenotype. We stress that this is not the case for other sources of data, such as from experiments that assay all possible 2*^L^* combinations of wildtype and mutant alleles at *L* sites (Weinreich et al., 2006), all possible combinations of fewer than 20 amino acids at *L* sites (Jacquier et al., 2013; Pokusaeva et al., 2019; Bank et al., 2016), or deep mutational scanning experiments that assay all 19*L* possible single-amino acid changes to a wild-type sequence of length *L* (Kinney and McCandlish, 2019; de Visser and Krug, 2014).

Leveraging the availability of these combinatorially complete data sets, here we characterize the topographies of three empirical adaptive landscapes under the standard genetic code, as well as under hundreds of thousands of rewired codes, and perform population-genetic simulations on these landscapes. We show that robust genetic codes, i.e. genetic codes that tend to preserve physicochemical properties of amino acids, tend to produce smooth adaptive landscapes with few peaks and, consequently, allow evolving populations to reach on average higher fitness. We also show that under robust genetic codes, the set of high-fitness sequences is more densely connected than under less robust codes. Thus, the robustness of a genetic code not only helps to mitigate the potentially deleterious effects of replication and translation errors, but it also transforms the problem of molecular evolution from one that depends on the vicissitudes of individual mutations into one where evolving populations can readily find mutational paths toward adaptation.

## 2 Results

### 2.1 Data

We construct empirical adaptive landscapes using three combinatorially complete data sets for two proteins. The first protein is GB1, a Streptococcal protein that binds immunoglobulin (Sjöbring et al., 1991; Sauer-Eriksson et al., 1995). Wu et al. (2016) experimentally assayed the binding affinity of GB1 to immunoglobulin for all 20^4^ = 160, 000 amino acid sequences at four protein sites (V39, D40, G41, and V54; Supp. Fig. S1), which are known to interact epistatically and influence binding affinity (Olson et al., 2014). In particular, they measured the relative frequencies of sequence variants before and after selection for binding immunoglobulin. Binding affinities are then defined as log enrichment ratios (Methods). The second protein is ParD3, a bacterial antitoxin that is part of the ParD-ParE family of toxin-antitoxin systems, which are commonly found on bacterial plasmids and chromosomes (Fraikin et al., 2020). Such systems comprise a toxin that inhibits cell growth unless bound and inhibited by the cognate antitoxin. Lite et al. (2020) experimentally assayed bacterial cell growth for all 20^3^ = 8, 000 amino acid sequence variants at 3 sites in ParD3 (D61, K64, E80; Supp. Fig. S1), in the presence of its cognate toxin ParE3, as well as a related, but non-cognate toxin ParE2. This resulted in two datasets, one per toxin, in which cell growth was used as a quantitative readout of the degree to which individual ParD3 variants antagonize a given toxin.

Following the protein evolution literature (Wu et al., 2016; Tokuriki and Tawfik, 2009; Romero and Arnold, 2009), we assume that fitness is directly proportional to the binding affinity (GB1) or growth rate (ParD3), and will use the term ‘fitness’ generically for both landscapes from now on. Using the raw measurements described above (binding affinities and cell growth), we inferred the fitness values, as well as imputed the missing sequence variants (6.6% of the GB1 data set) using empirical variance component regression (Zhou et al., 2022) (Methods and Supp. Fig. S2).

For each of the three data sets, we constructed adaptive landscapes using the standard genetic code, as well as hundreds of thousands of rewired codes. Specifically, we represented each mRNA sequence of length 12 (GB1) or 9 (ParD-ParE2 and ParD-ParE3), respectively, as a vertex in a mutational network and connected vertices with an edge if their corresponding sequences differed by a single point mutation (Wagner, 2009) (Methods). We labeled each vertex with the fitness of its corresponding translation under a given genetic code, thus defining the “elevation” of each coordinate in genotype space.

At the amino acid level (i.e., assuming no genetic code), the resulting landscapes differ significantly in their ruggedness: Whereas the GB1 landscape is fairly rugged (an additive model explains only 52.6% of the variance in the fitness values; Methods), the ParE2 and ParE3 landscapes are nearly additive (an additive model explains 84.5% and 86.0% of the variance, respectively; Methods).

### 2.2 More robust codes cause smoother adaptive landscapes

How does the robustness of a genetic code influence adaptive landscape topography? To answer this question, we generated 100,000 rewired genetic codes by amino acid permutation, a rewiring scheme that preserves the synonymous codon block structure of the standard genetic code, but randomly permutes the 20 amino acids amongst these blocks (Haig and Hurst, 1991; Freeland and Hurst, 1998). We quantified the robustness of each code as the proportion of point mutations that do not change the physicochemical properties of amino acids, using the properties defined in Pines et al. (2017) (Supp. Fig. S3; Methods). According to this measure, the robustness of the standard genetic code is 0.385, meaning that 38.5% of point mutations do not change the physicochemical properties of amino acids. In comparison, the range of code robustness for the 100,000 rewired codes is between 0.257 and 0.462, with a median of 0.336, such that 5.48% of these codes exhibit robustness greater than or equal to the standard code. Therefore, when defining robustness in terms of multiple amino acid properties, the standard genetic code is highly robust, but not surprisingly so, an observation that has been made previously (Haig and Hurst, 1991) and to which we return later.

To study the relationship between code robustness and adaptive landscape topography, we constructed an adaptive landscape using each of the 100,000 rewired genetic codes, for each of the three combinatorially complete data sets, and characterized the topographies of these landscapes using three measures of landscape ruggedness (Aguilar-Rodríguez et al., 2017): the number of adaptive peaks, the prevalence of various types of epistasis, and the proportion of accessible mutational paths to the global peak (Methods). Below, we focus our analyses on the GB1 data, and report analogous results for the ParD data in Supp. Tab. S1.

#### 2.2.1 Adaptive peaks

The number of adaptive peaks is a straightforward measure of landscape ruggedness, and thus of evolvability. The more local peaks a landscape has, the more likely an evolving population is to become trapped on one of these peaks, thus precluding the generation of further adaptive phenotypic variation. Under the standard genetic code, the GB1 landscape comprises 115 adaptive peaks, whereas under the 100,000 rewired codes, the number of adaptive peaks ranges from 97 to 478, with a median of 231. We note that Wu et al. (2016) reported the GB1 landscape to contain only 30 peaks in their analyses that did not consider the genetic code.

Fig. 2A shows the number of adaptive peaks in relation to code robustness, revealing that more robust genetic codes tend to produce adaptive landscapes with fewer peaks than less robust codes (Pearson’s correlation *R* = *−*0.144, *p <* 2.2 *·* 10*^−^*^16^, GB1; *R* = *−*0.119, *p <* 2.2 *·* 10*^−^*^16^, ParD-ParE2; *R* = *−*0.035, *p <* 2.2 *·* 10*^−^*^16^, ParD-ParE3). However, these trends are relatively weak, such that for any level of code robustness, there is considerable variation in the number of peaks. For example, for the 5.48% of codes with robustness greater than or equal to the standard code, the number of peaks ranges from 115 to 403. Strikingly, among all 100,000 codes, only 0.037% of the corresponding landscapes have less than or equal the number of peaks in the landscape produced by the standard code. The GB1 landscape is therefore exceptionally smooth under the standard genetic code. This, however, is not true for the ParD-ParE2 and ParD-ParE3 landscapes, where the number of peaks in the landscape under the standard genetic code lies in the 0.326 and 0.570 quantile, respectively (Supp. Tab. S2). This highlights that the influence of a genetic code on protein evolvability can be protein-specific.

**Figure 2:**
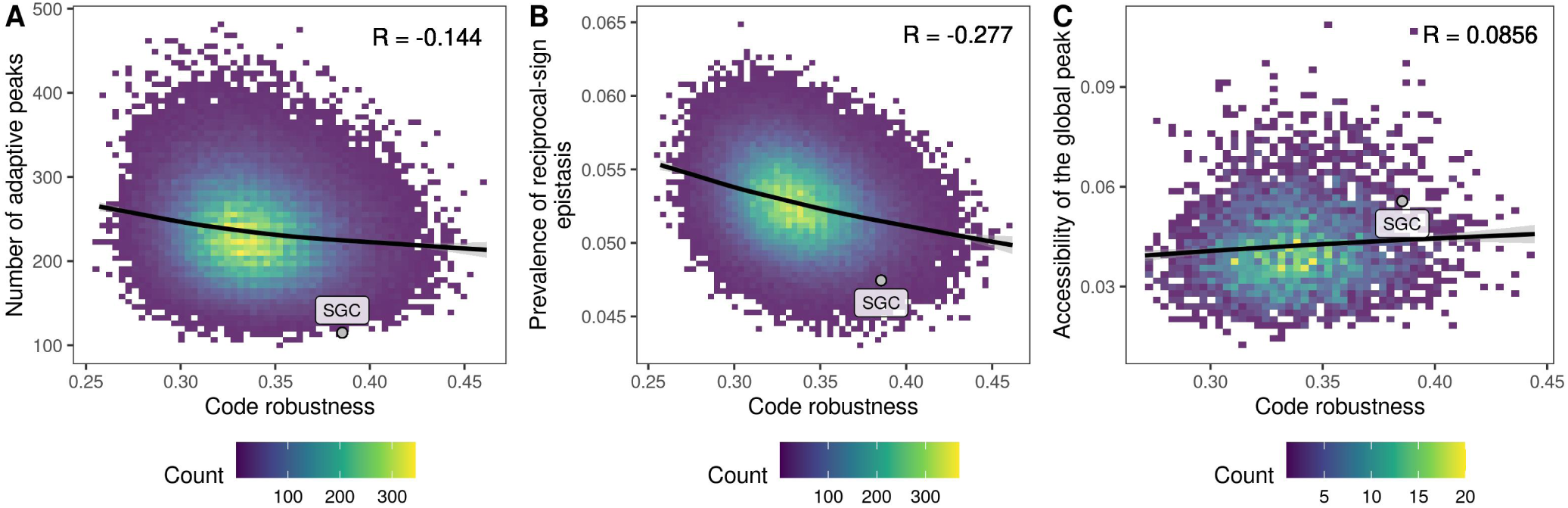
More robust codes result in smoother adaptive landscapes. Three measures of landscape ruggedness are shown in relation to code robustness, defined as the proportion of point mutations that do not change the physicochemical properties of amino acids. (A) The number of adaptive peaks, (B) the prevalence of reciprocal sign epistasis, and (C) the proportion of mutational paths to the global peak that are accessible. Panel (C) shows only genetic codes that preserve the size of the global peak relative to the standard genetic code and in which none of the amino acids contained in the global peak (WWLA) are encoded by the split codon block (*n* = 3, 769). In each panel, the labeled point denotes the standard genetic code. All results pertain to the GB1 landscape. Analogous results for the ParD landscapes can be found in Supp. Tab. S1.

#### 2.2.2 Epistasis

Epistasis, where a mutation’s effect depends on the genetic background in which it occurs, is a cause of landscape ruggedness (Weinreich et al., 2005; Poelwijk et al., 2007). It can be classified into three types — magnitude, simple sign, and reciprocal sign (Weinreich et al., 2005) (Methods). Reciprocal sign epistasis occurs when two mutations each have a positive (negative) effect on fitness, but each mutation has negative (positive) effect when introduced in the background of the other mutation. That is, the sign of each mutation’s effect flips when introduced in the presence of the other mutation. Reciprocal sign epistasis forms local valleys in an adaptive landscape, which preclude the generation of at least some adaptive phenotypic variation, thus decreasing evolvability. To measure the prevalence of these three types of pairwise epistasis, we randomly sample a large number of squares in each adaptive landscape’s underlying mutational network, each of which contains an mRNA sequence variant, two of its single-mutant neighbors, and a double mutant that can be constructed from the single mutants. Based on the fitness values of these four sequences, we classify the square as exhibiting magnitude, simple sign, or reciprocal sign epistasis (Methods).

Because more robust codes tend to produce adaptive landscapes with fewer adaptive peaks (Fig. 2A), we expect landscapes produced under more robust codes to exhibit less reciprocal sign epistasis than landscapes produced under less robust codes. Fig. 2B confirms this expectation, showing a negative correlation between reciprocal sign epistasis and code robustness (*R* = *−*0.277, *p <* 2.2 *·* 10*^−^*^16^, GB1; *R* = *−*0.262, *p <* 2.2 *·* 10*^−^*^16^, ParD-ParE2; *R* = *−*0.191, *p <* 2.2 *·* 10*^−^*^16^, ParD-ParE3). Similarly, simple sign epistasis, which contributes to landscape ruggedness to a lesser extent than reciprocal sign epistasis, because it involves only a single sign flip, also exhibits a negative correlation with code robustness (Supp. Tab. S1). However, these two forms of epistasis characterize only a minority of squares in the GB1 landscape, ranging in prevalence from 18.8% to 20.9% for the 1% least robust codes, to 18.0% to 20.5% for the 1% most robust codes. The remaining majority of squares exhibit either no epistasis or magnitude epistasis (Methods; Supp. Tab. S1). Thus, sign epistatic interactions are relatively rare in the GB1 landscape, and their prevalence is further reduced by increasing code robustness. Robust genetic codes therefore diminish the kinds of epistatic interactions that cause landscape ruggedness.

#### 2.2.3 Global peak accessibility

One consequence of landscape ruggedness is that the global adaptive peak may be less accessible to an evolving population, which may instead follow mutational paths to local adaptive peaks. We therefore expect that the global adaptive peaks of landscapes produced under more robust codes will be more accessible than those of landscapes produced under less robust codes. To test this, we quantified the mutational accessibility of the global peak of each landscape by calculating the probability that a randomly chosen, direct mutational path that starts at a randomly chosen mRNA sequence and ends at the global peak is accessible, meaning that fitness increases monotonically along the path (Aguilar-Rodríguez et al., 2017; Weinreich et al., 2006; Franke et al., 2011). In contrast to expectation, we observe that the global peaks of landscapes produced under more robust codes are only marginally more accessible than those of landscapes produced under less robust codes for the GB1 landscape (*R* = 0.0238, *p* = 5.45 *·* 10*^−^*^14^), not significantly more accessible for the ParD-ParE2 landscape (*R* = 0.0052, *p* = 0.099), and significantly less accessible for the ParD-ParE3 landscape (*R* = *−*0.151, *p <* 2.2 *·* 10*^−^*^16^).

We reasoned that the accessibility of the global peak might be confounded by its size: As the number of codons encoding an amino acid ranges from 1 to 6 in the amino acid permutation codes, the number of distinct mRNAs encoding the protein sequence with the highest fitness value ranges from 2 (= 1 *·* 1 *·* 2; there are only two codon blocks of size 1 and in all three landscapes the global peak sequence – WWLA for GB1, ELK for ParD-ParE2, and DWE for ParD-ParE3 – consists of three different amino acids) to 6^4^ = 1296 for GB1 or 6^3^ = 216 for ParD-ParE2 and ParD-ParE3 (there are three codon blocks of size 6, hence all three amino acids contained in the global peak sequence may be encoded by a 6-codon block). Indeed, we observe that the mutational accessibility of the global peak is strongly correlated with its size (*R* = 0.801, *p <* 2.2 *·* 10*^−^*^16^, GB1; *R* = 0.710, *p <* 2.2 *·* 10*^−^*^16^, ParD-ParE2; *R* = 0.810, *p <* 2.2 *·* 10*^−^*^16^, ParD-ParE3; Supp. Fig. S4). Moreover, due to the fact that one of the synonymous codon blocks is split (UCN and AGY, with N denoting any nucleotide and Y denoting U or C; encoding serine in the standard genetic code), there might be several disconnected regions of the landscape encoding the protein sequence with the highest fitness value. When this is the case, the mutational accessibility of the global peak is significantly higher compared to codes where the global peak comprises a single connected region in genotype space (mutational accessibility 0.086 vs. 0.055, *p <* 2.2 *·* 10*^−^*^16^, GB1; 0.264 vs. 0.200, *p <* 2.2 *·* 10*^−^*^16^, ParD-ParE2; 0.180 vs. 0.131, *p <* 2.2 *·* 10*^−^*^16^, ParD-ParE3; Supp. Fig. S5). In order to make the landscapes more comparable, we restricted our analysis to only those landscapes in which the size of the global peak, in terms of number of mRNAs encoding the corresponding protein sequence, is the same as in the standard genetic code and, moreover, none of the amino acids contained in the global peak sequence are encoded by the split codon block. There were 3,769 such landscapes for GB1, 12,059 for ParD-ParE2, and 6,781 for ParD-ParE3. In this subset of landscapes, we observe the expected positive correlation between code robustness and accessibility of the global peak for the GB1 and ParD-ParE2 landscapes (*R* = 0.086, *p* = 1.43 *·* 10*^−^*^7^, GB1; *R* = 0.0920, *p <* 2.2 *·* 10*^−^*^16^, ParD-ParE2) (Fig. 2C) and a weak negative relationship between the two quantities for the ParE3 landscape (*R* = *−*0.085, *p* = 3.01 *·* 10*^−^*^12^).

While statistically significant, the strength of the correlation between code robustness and global peak acces-sibility is weak and the magnitude of the effect is not large (mean global peak accessibility 0.055 vs. 0.058 for the 1% least and most robust codes, respectively; Fig. 2C). Given the low prevalence of sign epistatic interactions in the landscapes generated under even the least robust genetic codes, we reasoned that the range of landscape ruggedness observed in our data may simply be too small to observe a strong positive correlation between global peak accessibility and code robustness. We therefore artificially inflated the ruggedness of the GB1 landscape under the standard genetic code by separately increasing the number of local peaks and the prevalence of reciprocal sign epistasis (Methods), producing landscapes that ranged in their number of local peaks from 115 to 3,356 and in their prevalence of reciprocal sign epistasis from 0.047 to 0.130. With these landscapes, we observed a strong correlation between global peak accessibility and the two measures of landscape ruggedness (*R* = *−*0.893, *p <* 2.2 *·* 10*^−^*^16^, number of peaks vs. mutational accessibility of the global peak; *R* = *−*0.987, *p <* 2.2*·*10*^−^*^16^, prevalence of reciprocal-sign epistasis vs. mutational accessibility of the global peak; Supp. Fig. S6). Moreover, we observe that the moderate effect size is consistent with the range of reciprocal-sign epistasis prevalence in the amino acid permutation codes (Supp. Fig. S6B) and that the expected effect size based on the range in the number of peaks would be even lower (Supp. Fig. S6A). In sum, the mutational accessibility of the global peak is strongly influenced by the number of its constituent mRNA sequences and whether they occupy disjoint regions of genotype space, and only weakly influenced by code robustness, due to the limited range of landscape ruggedness produced by the 100,000 amino acid permutation codes.

#### 2.2.4 Random codon assignment codes

So far, we have computationally rewired the genetic code using amino acid permutation, which is only one of many possible rewiring schemes (Caporaso et al., 2005; Wichmann and Ardern, 2019; Rozhoňová and Payne, 2021). To test the sensitivity of our results to choice of rewiring scheme, we repeated the analyses above for 100,000 genetic codes generated by randomly assigning an amino acid meaning to each of the 61 sense codons, ensuring that each of the 20 amino acids is assigned at least one codon (Caporaso et al., 2005; Rozhoňová and Payne, 2021; Tripathi and Deem, 2018). We refer to these as ‘random codon assignment’ codes. These differ from the codes generated using amino acid permutation by lacking the synonymous codon block structure of the standard genetic code. Consequently, their average robustness is much lower than that of the amino acid permutation codes (*p <* 2.2 *·* 10*^−^*^16^, Welch two sample t-test; Supp. Fig. S7). Consistent with our previous observations (Fig. 2), we find that increasing code robustness decreases landscape ruggedness under this alternative rewiring scheme. Specifically, more robust codes yield landscapes with fewer local peaks and a reduced prevalence of sign epistasis, as well as a marginal increase in the accessibility of the global adaptive peak (Supp. Tab. S3). Our results are thus qualitatively insensitive to this choice of rewiring scheme.

### 2.3 Relevant amino acid properties are protein-specific

Our measure of code robustness assigns amino acids to discrete groups based on seven key physicochemical properties, such as whether the amino acids are acidic or basic (Supp. Fig. S3; Methods). However, there are hundreds of physicochemical properties that can be used to characterize amino acids. For example, the AAindex database includes 566 such properties (Kawashima et al., 1999; Kawashima and Kanehisa, 2000), which belong to four higher-level categories: “alpha and turn propensity”, “beta propensity”, “hydrophobicity’, and “other” (Tomii and Kanehisa, 1996; Bartonek et al., 2020). To better understand which particular amino acid properties drive the correlation between code robustness and landscape ruggedness in our three datasets, we recomputed the robustness of each of the 100,000 amino acid permutation codes in terms of each of the 553 properties from the AAindex database that do not contain any missing values, separately (Methods), and calculated the correlation with our various measures of landscape ruggedness. We determined the statistical significance of the correlations by comparison with a null distribution calculated from 1,000,000 amino acid “properties” with randomly chosen values, and corrected for testing multiple hypotheses (see Methods).

We observe many amino acid properties that are consistent with our previous observation that more robust codes cause smoother adaptive landscapes, and only very few that support the opposite statement (i.e., less robust codes implying smoother landscapes; Supp. Data S1-S3). For example, for the GB1 data, 169 out of the 553 properties (30.6% of the tested properties) exhibit a significant negative correlation between code robustness and the number of peaks, whereas there are only 2 properties (i.e., less than 0.4% of the database) for which the opposite is true. For GB1, the statistically significant properties are enriched in beta-sheet propensity indices and, less consistently across the different landscape ruggedness measures, in alpha-helix propensity and hydrophobicity indices (Supp. Tab. S4). The importance of the preservation of hydrophobicity is consistent with V39 and V54 having buried sidechains; however, the consistent significance of beta-sheet propensity indices is somewhat surprising, as only one of the four residues is located in a beta sheet (Supp. Fig. S1). The properties that are significant for the two ParD3 landscapes, in contrast, are consistently enriched in alpha-helix propensity indices (Supp. Tab. S4), which is consistent with all three screened residues being found in alpha-helices (Supp. Fig. S1). In sum, the amino acid properties most relevant to code robustness are protein-specific, and depend on the structural and functional properties of the assayed residues in each protein. Increasing code robustness relative to these properties generally results in smoother adaptive landscapes.

### 2.4 Evolutionary simulations reveal that robust genetic codes promote evolvability

Our analyses suggest that code robustness promotes evolvability by producing smooth adaptive landscapes with few peaks and little sign epistasis. As a consequence, we anticipate evolving populations to obtain higher fitness, on average, when translating proteins using more robust codes than when using less robust codes. To determine if this is the case, we turn to evolutionary simulations. We studied two different models of adaptive walks: greedy adaptive walks (de Visser and Krug, 2014) and weak mutation adaptive walks (Gillespie, 1984). The greedy adaptive walks model adaptive evolution of a large population with pervasive clonal interference, such that all possible point mutations to a sequence are simultaneously present in the population, and the fittest of these variants goes to fixation. For each of the 100,000 amino acid permutation landscapes and each of the three datasets, we initialized the walks in each of the 61*^L^* possible nucleotide sequences that did not contain a stop codon (61^4^ = 13, 845, 841 sequences for GB1 dataset, 61^3^ = 226, 981 sequences for the two ParD3 datasets). We terminated a walk when it reached a local or global adaptive peak, and recorded the fitness of that peak sequence (Methods). The weak mutation adaptive walks represent adaptive evolution under the regime where mutations occur so infrequently that any mutation will either go to extinction or to fixation prior to the arrival of a subsequent mutation. The probability of fixation depends on both the improvement in fitness and the population size, which controls the strength of genetic drift. For each genetic code, each landscape and each choice of one of four different population sizes, we simulated 100,000 random walks, initialized in randomly chosen sequences. In each step of the walk, a neighboring sequence was proposed and accepted with probability determined by the Moran process (Moran, 1958) (Methods). We recorded the fitness values reached after 500 proposed mutations. In the following, we focus on the greedy adaptive walks, because they are easier to analyze due to their deterministic nature and because the large population assumption is a more suitable description of directed protein evolution experiments. The weak mutation results are mentioned briefly at the end of this section and provided in detail in the Supplement.

Fig. 3A shows the average fitness reached by the greedy adaptive walks in relation to code robustness. As expected from our landscape-based analyses, evolving populations reached higher fitness, on average, when translating proteins using more robust genetic codes for the GB1 and ParD-ParE2 landscapes (*R* = 0.107, *p <* 2.2 *·* 10*^−^*^16^, GB1; *R* = 0.121, *p <* 2.2 *·* 10*^−^*^16^, ParD-ParE2). The results for the ParD-ParE3 landscape were not statistically significant (*R* = *−*0.004, *p* = 0.182). Similar to the analysis of accessible paths above, we reasoned that the lack of correlation in the ParD-ParE3 data set might be caused by variation in the size of the global peak, such that larger global peaks are easier to “find” than smaller global peaks, simply because they contain more mRNA sequences. Indeed, we observe a positive correlation between the size of the global peak and mean fitness reached by the greedy adaptive walks in all three data sets (*R* = 0.285, *p <* 2.2 *·* 10*^−^*^16^, GB1; *R* = 0.377, *p <* 2.2 *·* 10*^−^*^16^, ParD-ParE2; *R* = 0.352, *p <* 2.2 *·* 10*^−^*^16^, ParD-ParE3). We thus again restricted our analysis to those genetic codes for which the size of the global peak is the same as under the standard genetic code and occupies a single connected region in genotype space. In this subset of codes, we consistently observe a positive correlation between code robustness and mean fitness reached by the greedy adaptive walks (*R* = 0.130, *p* = 1.199 *·* 10*^−^*^15^, GB1; *R* = 0.180, *p <* 2.2 *·* 10*^−^*^16^, ParD-ParE2; *R* = 0.092, *p* = 1.026 *·* 10*^−^*^12^, ParD-ParE3).

**Figure 3:**
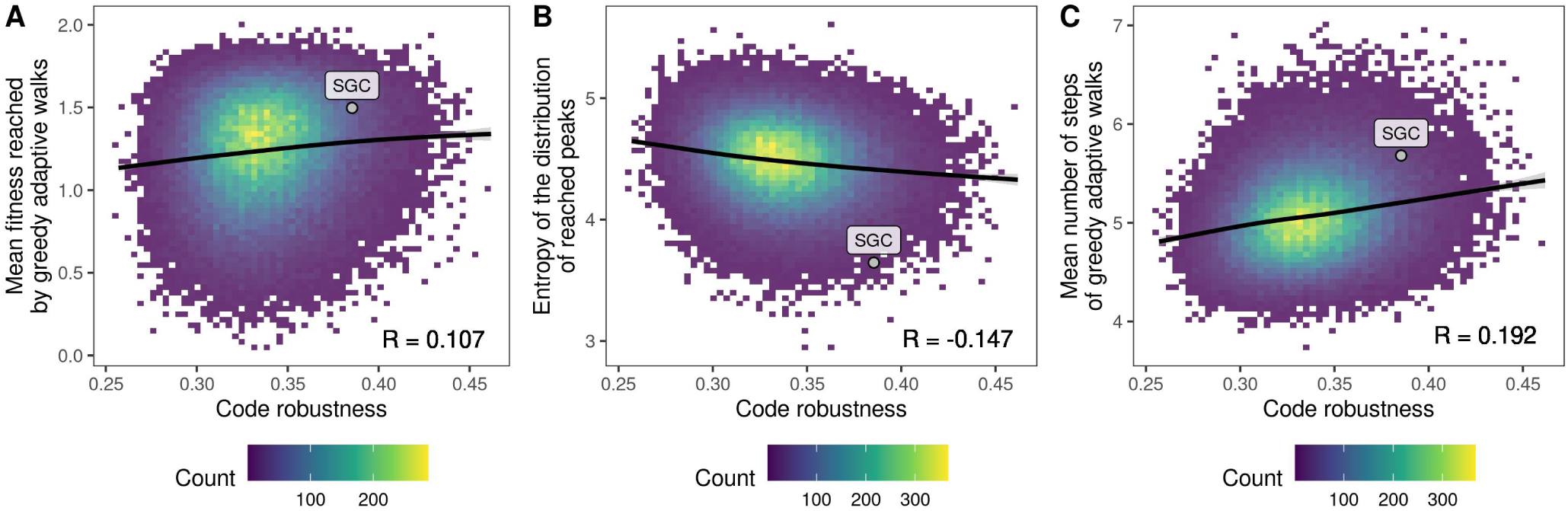
Robust genetic codes promote evolvability. Relationship between code robustness and results of greedy adaptive walks. The labeled point denotes the results obtained using the standard genetic code. Data pertain to GB1.

What is the cause of this correlation? To answer this question we further investigated the subsets of landscapes that preserve the size of the global peak and in which the global peak consists of a single connected region in genotype space. A potential explanation is that under more robust codes the probability of reaching the global peak increases, because increasing code robustness increases global peak accessibility. This is indeed the case for the ParD-ParE2 (*R* = 0.109, *p <* 2.2 *·* 10*^−^*^16^) and ParD-ParE3 (*R* = 0.033, *p* = 0.00666) landscapes. However, the strength of the correlation is much weaker than the correlation between code robustness and mean fitness, and for the GB1 landscape the global peak is reached less often under robust codes (*R* = *−*0.025, *p* = 4.846 *·* 10*^−^*^13^). Indeed, even when considering only those adaptive walks that terminated on a local peak, we still observe a positive correlation between code robustness and mean fitness (*R* = 0.131, *p* = 7.158 *·* 10*^−^*^16^, GB1; *R* = 0.114, *p <* 2.2 *·* 10*^−^*^16^, ParD-ParE2; *R* = 0.081, *p* = 2.076 *·* 10*^−^*^11^, ParD-ParE3). This trend is not caused by the local peaks being on average higher under robust codes: in fact, in all three data sets, there is a negative correlation between code robustness and the mean height of peaks (*R* = *−*0.094, *p <* 2.2 *·* 10*^−^*^16^, GB1; *R* = *−*0.136, *p <* 2.2 *·* 10*^−^*^16^, ParD-ParE2; *R* = *−*0.106, *p <* 2.2 *·* 10*^−^*^16^, ParD-ParE3). The only possible explanation of the correlation between code robustness and mean fitness reached in adaptive walks is that under robust codes the basins of attraction of the high-fitness peaks are relatively larger compared to those of less robust codes; in other words, under robust codes the adaptive walks tend to converge on a smaller number of high-fitness peaks. Indeed, we observe that with increasing code robustness the Shannon entropy of the distribution of peaks reached by the greedy walks decreases (Fig. 3B; Methods).

To model the relationship between code robustness, peak height, and basin of attraction more explicitly, we fitted a linear model that, for each of the genetic codes, predicts the logarithm of the size of the basin of attraction of a peak as a linear function of its height, log(*size of basin*) = *β*_0_ + *β*_1_(*peak height*). The *β*_1_ coefficient controls how fast the size of the basin changes with peak height; for example, using the standard genetic code and the GB1 landscape, the coefficient is 0.591, meaning that if the peak height increases by 1, the size of the basin is expected to increase exp (0.591) *≈* 1.8-times. The bigger the *β*_1_ coefficient, the faster the basin of attraction grows with peak height and the more concentrated the ends of the adaptive walks are on the high peaks. Having computed the *β*_1_ coefficients for all genetic codes in the subset of genetic codes that preserve the size of the global peak, we then correlated them with the corresponding robustness. As expected, we observe a positive correlation in all three data sets (*R* = 0.141, *p <* 2.2 *·* 10*^−^*^16^, GB1; *R* = 0.250, *p <* 2.2 *·* 10*^−^*^16^, ParD-ParE2; *R* = 0.228, *p <* 2.2 *·* 10*^−^*^16^, ParD-ParE3). These analyses thus show that under robust genetic codes, evolutionary trajectories to adaptation become more predictable, in that they converge on a smaller number of adaptive peaks, and moreover, they preferentially converge on high-fitness peaks.

We also observe that the average length of the walks tended to be longer under robust codes (Fig. 3C; 4.89 vs. 5.30 steps, on average, for the 1% least and most robust codes, respectively), revealing that the benefit of increased fitness afforded by code robustness comes at the cost of longer evolutionary trajectories to adaptation. This is in line with our observations concerning landscape ruggedness. In landscapes with many local peaks, a greedy walk is more likely to be initialized near one of these peaks, which it will likely ascend in only a small number of mutational steps. In contrast, in landscapes with few local peaks, a greedy walk is more likely to be initialized farther away from one of these peaks, thus increasing the length of the mutational path to adaptation, be it to a local or global peak.

We observe qualitatively the same results for the weak mutation adaptive walks (Supp. Tab. S6) and using codes constructed by random codon assignment (Supp. Tab. S7 and S8). In sum, robust genetic codes promote evolvability by producing smooth adaptive landscapes in which high-fitness peaks have large basins of attraction, thus increasing the mean fitness reached by simulated populations of evolving proteins, the time to convergence, and the predictability of the evolutionary process.

### 2.5 The genetic code governs the genetic architecture of long-term molecular evolution

In the previous section, we studied a short-term adaptive process, in which high-fitness protein variants evolve from low-fitness variants via mutation and selection. However, once an evolving population reaches high fitness, it behaves like a random walk amongst the mutationally-interconnected set of high-fitness variants, which we refer to here as a genotype network. Like the topographical structure of a fitness landscape, the topological structure of a genotype network has a strong influence on evolvability (Wagner, 2008; Schuster et al., 1994; Lipman et al., 1991). For example, evolvability is diminished when high-fitness variants tend to be connected through long branch-like structures, because traversing these requires a large number of mutations that must occur in a specific order. In contrast, grid-like structures, in which different regions of a protein sequence may evolve independently, enhance evolvability, because such modularity limits the dependence on the order in which mutations occur.

To assess how different code rewirings influence genotype network topology, we apply a visualization technique that captures the dynamics of a finite population evolving on a genotype network at mutation-selection-drift balance (McCandlish, 2011). Intuitively, the resulting “diffusion axes” capture the main barriers to diffusion in sequence space, and distances between sequences in this low-dimensional representation reflect the expected time to evolve from one sequence to another (Methods). Moreover, the diffusion axes have a natural ordering, such that the most important barriers to diffusion are displayed along Diffusion Axis 1, the next most important along Diffusion Axis 2, etc.

In an earlier study, we used this technique to explore the structure of the GB1 landscape at the amino acid level (Zhou and McCandlish, 2020) and found that it consists of three main regions of high-fitness protein variants that differ primarily in the placements of a small non-polar and bulkier amino acids at positions 41 and 54. The first and largest of these regions is characterized by 41G, which is compatible with most amino acids at position 54 and contains the wild-type sequence VDGV; we will refer to this as Region 1. The second region involves Gly at 54, while tolerating Thr at 54 in some contexts, together with Leu or Phe at position 41, and we will refer to this as Region 2. The final region, Region 3, is characterized by 54A, which can be paired at position 41 with Cys, Ser or Ala, and to a lesser extent Leu and Phe. At the amino acid level, we found that these different regions are mutationally interconnected through smaller sets of high fitness sequences that act as transition complexes in protein space. Specifically, 41G-54G connects Regions 1 and 2, 41G-54A connects Regions 1 and 3, and 41C-54G connects Regions 2 and 3 (Zhou and McCandlish, 2020). Here, we consider how the genetic code, standard or otherwise, reshapes the structure of these regions and restricts their mutational interconnections, focusing on the standard genetic code, as well as the two most and two least robust of the amino acid permutation codes (see Supp. Fig. S8 for the corresponding codon tables).

#### 2.5.1 Standard genetic code

Fig. 4A shows our visualization of the GB1 landscape under the standard genetic code. In the visualization, each vertex is an mRNA sequence and edges connect sequences that differ by a single nucleotide substitution. Fig. 4A top shows all possible mRNA sequences, while Fig. 4A bottom shows the structure of the genotype network formed by the fittest 1% of sequences, and both the top and bottom panels show an embedding based on the first three diffusion axes, which in this case are sufficient to display the major qualitative features of the genotype network. Looking down Diffusion Axis 1 (Fig. 4A, vertical axis), we see that Region 1 (characterized by 41G) is at the top of this axis whereas Region 2 (characterized by 41F or L and 54G or T) is at the bottom, indicating that under long-term purifying selection for GB1 functionality it would take an extremely long time for a population to evolve from Region 1 (which contains the wild-type sequence) to Region 2. The reason is that under the standard genetic code, neither 41F nor 41L is accessible from 41G, and so high-fitness paths from Region 1 to Region 2 instead pass through Region 3, which remains accessible from both Regions 1 and 2.

**Figure 4:**
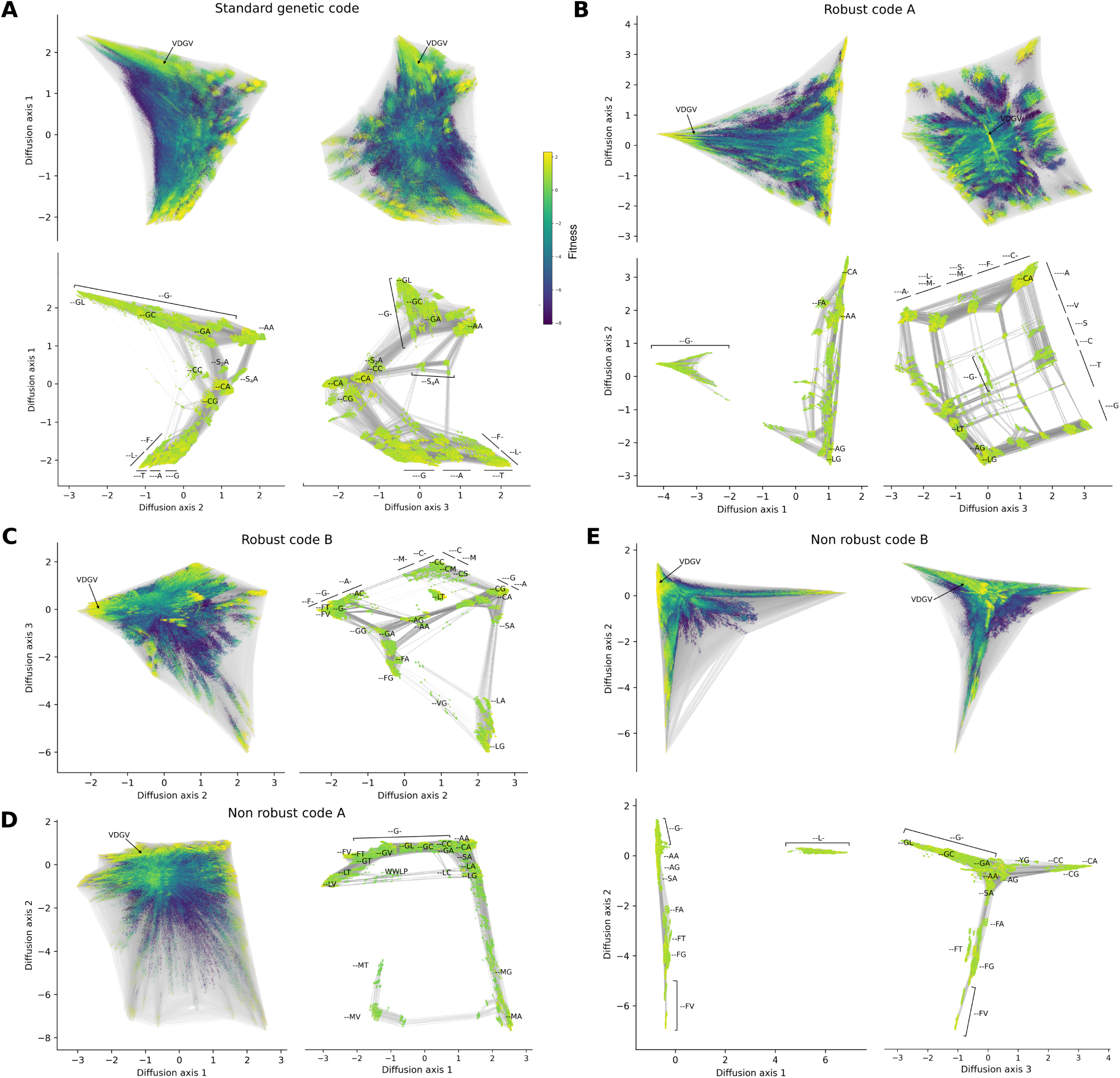
The genetic code governs genotype network topology and the genetic architecture of long-term molecular evolution. Fitness landscape for GB1 under the (A) standard genetic code, (B, C) the two most and (D, E) the two least robust codes in the amino acid permutation set. Vertices represent 12-nucleotide sequences and edges connect vertices if their corresponding sequences differ by a single point mutation. Vertex color represents protein fitness (color bar in (A) applies to all panels). Vertices are placed at the coordinates along the diffusion axes, which at a technical level are defined by the subdominant eigenvectors of the rate matrix describing the weak mutation dynamics (McCandlish, 2011) (see methods for details). For each pair of diffusion axes shown, there are two subpanels: one that shows all ≈16 million genotypes, with the location of the sequences encoding the wild-type protein sequence VDGV marked, and another that shows only the genotype network of high-fitness variants (top 1% of fitness distribution), which better shows the connectivity between high-fitness regions and which is annotated with the protein sequence features that characterize each cluster or subset of nucleotide sequences.

Besides reducing the connectivity between regions of high-fitness sequences that are accessible to each other in amino acid sequence space, different genetic codes can also restrict the connectivity within these high-fitness regions, or can even break such a region into several disconnected pieces. Under the standard genetic code, we see both of these phenomena, as Region 1 is spread along Diffusion Axis 2, with 41G-54L at one end and the connection to Region 3, 41G-54A, at the other, and Region 2 is in fact broken into two pieces (defined by 54G and 54T, which are not accessible to each other under the standard genetic code) and spread along Diffusion Axis 3. These two pieces of Region 2 are then connected by a portion of Region 3 consisting of 41L or 41F together with 45A, which is adjacent to both pieces of Region 2 (since under the standard genetic code Gly and Thr are both accessible from Ala). The end result is that while in amino acid sequence space any two highly fit sequences are typically connected by a high-fitness path with at most 4-5 substitutions, in nucleotide space a typical trajectory from Region 1 to Region 2 contains far more substitutions, many of which must be accumulated in a specific order, e.g., Supp. Fig. S9 highlights a path that requires 11 mutations even without including substitutions at positions 39 or 40, synonymous changes, or reversions.

Nonetheless, while any genetic code acts to reduce evolvability relative to amino acid sequence space, we see that the standard genetic code manages to retain evolvability in several different ways. First, we see the genotype network remains connected, such that high-fitness protein variants in distant reaches of sequence space are mutationally-accessible from one another via a series of intermediates that are also of high fitness. This connectedness is important for evolvability, because an evolving population can diffuse across the genotype network to produce new phenotypic variants, and populations with sufficiently high mutation rates will accumulate genetic diversity, which can be revealed as phenotypic variation upon environmental change (Zheng et al., 2019). Second, whereas traversing from one end of the network to another typically requires many mutations, this is not always the case. For example, in Fig. 4A bottom right, we can see that distant pieces of the genotype network are in fact accessible to each other via Ser_4_, the larger of the two disconnected sets of codons for Ser (named Ser_2_ and Ser_4_ for the number of codons in each set (Maeshiro and Kimura, 1998)). We call such a path a “wormhole”, as it allows a population to jump from one region of the genotype network to another. Finally, an important aspect of evolvability is modularity, which in this case refers to amino acids positions that can evolve relatively independently from each other and which produces extended regions of amino acid sequence space where mutations can be accumulated in any order. Under any given genetic code, such regions can either remain connected or be broken into separated pieces, with the maintenance of connectivity resulting in a grid-like region of the visualization. We saw such a region already at the bottom of Diffusion Axis 1, where Phe and Leu at position 41 can be combined with any of Gly, Ala and Thr at position 54.

#### 2.5.2 Robust genetic codes

We have shown that robust genetic codes tend to produce smooth fitness landscapes, but also that for any level of code robustness, there is considerable variation in landscape ruggedness (Fig. 2). To understand how code robustness influences landscape topography in more detail, we visualize the GB1 landscapes and genotype networks under the two most robust of the amino acid permutation codes, named Robust Code A and Robust Code B (Supp. Fig. S8A and B). Under Robust Code A, Region 1 is disconnected from the remaining high-fitness protein variants in Regions 2 and 3, as can be seen when the landscape is visualized along Diffusion Axis 1 (Fig. 4B). The reason is that this code does not allow substitutions from Gly to any of the key intermediate amino acids that connect Region 1 to Regions 2 or 3 (Cys, Leu, Phe, Ala; Supp. Fig. S8A). In contrast, Regions 2 and 3 are connected, and form a large 2-dimensional grid-like structure that spreads out along Diffusion Axes 2 and 3. The grid is formed by variants at position 54 along Diffusion Axis 2 and by variants at position 41 along Diffusion Axis 3. However, the grid is imperfect, in that it contains some “holes” that correspond to incompatible amino acid combinations at positions 41 and 54, such as 41C-54T or 41S-54V. These incompatibilities reduce the number of accessible paths between the high-fitness sequences at the corners of the grid. The grid also contains “bypasses” that connect pairs of protein variants via indirect paths along each axis of the grid. For example, Cys and Met are directly accessible under Robust Code A, however, it is also possible to pass via a Phe intermediate (see the upper right corner of the grid).

The high-level topology of the genotype network is similar under Robust Code B, except that now it is the 41L-54T portion of Region 2 that is disconnected from the rest of the genotype network. The resulting fitness valley is the largest barrier to diffusion and hence dominates Diffusion axis 1 (Supp. Fig. S10), but the rest of the genotype network is connected and its structure is well-captured by Diffusion Axes 2 and 3 (Fig. 4C). Together, these visualizations show that robust codes tend to yield connected genotype networks with grid-like structures and bypasses that enhance evolvability, but also how even exceptionally robust codes can interact with the idiosyncrasies of a particular protein to break crucial links between high-fitness variants, yielding disconnected genotype networks and holes within the grid-like structures, both of which diminish evolvability.

#### 2.5.3 Non-robust genetic codes

To understand whether non-robust codes induce qualitatively different properties in the structure of the fitness landscape, we next study the structure of the genotype network under the two least robust of the amino acid permutation codes, named Non-Robust Code A and Non-Robust Code B (Supp. Fig. S8C and D). Under Non-Robust Code A (Fig. 4D), the genotype network adopts a long linear structure stretching from 41L-54V to 41M-54T. This greatly diminishes evolvability, both because traversing among amino acid sequences that differ even in only one position (e.g., 41L-54V and 41L-54G) may require many nucleotide mutations, and because only very few mutational paths exist between any pair of sequences in the genotype network. This is especially true for 41M sequences. 41M is compatible with amino acids at position 54 typical for Region 2 and 3, i.e., Ala, Gly, Thr, and Val, and 41M sequences thus usually cluster with sequences in Regions 2 and 3 (see e.g. Fig. 4B). However, under Non-Robust Code A, Met is encoded by a single codon (UGG; Supp. Fig. S8C), which differs by more than one mutation from any of the codons for the other high-fitness amino acids at position 41. As a result, only very few mutational paths exist from the 41M sequences to other high-fitness protein variants. Despite the mostly linear structure of the genotype space, though, this genotype network, similar to the one under the standard genetic code, has a “wormhole”, in which WWLP sequences bridge otherwise distant regions of the genotype network, namely the 41L-54V and 41L-54G sequences. Under Non-Robust Code B (Fig. 4E), we also observe limited connectivity amongst the high-fitness protein variants, with the 41L-sequences disconnected from the rest along Diffusion Axis 1 and the remaining sequences laid out in a star-like geometry along Diffusion Axis 2 and 3. Taken together, these visualizations illustrate how non-robust codes yield genotype networks with long branch-like structures that diminish evolvability.

### 2.6 Codon compression schemes reveal additional code features influencing evolvability

The results above suggest that it is possible, in principle, to design genetic codes that diminish or enhance protein evolvability by manipulating code robustness. However, engineering amino acid permutation codes in a living organism would require an extensive recoding of the genome, including the engineering of many orthogonal tRNA-aminoacyl tRNA synthetase pairs. For example, in the most robust of the 100,000 codes we analyzed, only two amino acids occupy the same synonymous codon block as in the standard genetic code (Supp. Fig. S8B). In contrast, to date, the synthetic biology community has engineered rewired genetic codes that change the meaning of up to only a small handful of codons (Mukai et al., 2017; Zürcher et al., 2022; Chin et al., 2003; Nyerges et al., 2023). There is therefore a large disconnect between the space of theoretically- and practically-realizable rewired genetic codes.

This motivated us to study a subset of rewired genetic codes that require only a small number of codon reassignments, as it may be possible to engineer these codes in a living organism using currently available technology. In particular, we studied the 57-codon *Eschirichia coli* genome synthesized by Ostrov et al. (2016), in which all occurrences of 7 codons, from 4 synonymous codon blocks, together with the corresponding tRNAs were removed from the genome and are thus theoretically free for reassignment (Supp. Fig. S11). Assuming each of the 4 synonymous blocks is reassigned to one amino acid or a stop signal (as might be required by the tRNA wobble rules (Dong et al., 1996; Agris et al., 2018)), there are in total 21^4^ = 194, 481 possible rewirings based on this compression scheme, one of them being the standard genetic code. We computationally generated all of these ‘Ostrov’ codes and repeated the landscape-based analyses and evolutionary simulations described above.

Relative to the permutation codes, the Ostrov codes exhibited an even stronger negative correlation between our measures of landscape ruggedness and code robustness (Supp. Tab. S9), as well as a stronger positive correlation between the average fitness reached by adaptive walks (greedy or random) and code robustness (Supp. Tab. S10 and S11). Notably, the range of the landscape ruggedness measures, e.g., in the number of peaks, is roughly the same as for the amino acid permutation codes, even though the Ostrov codes exhibit a much smaller range of code robustness (from 0.330 to 0.406, as compared to 0.257 to 0.462 for the permutation codes). Because the Ostrov codes differ from the permutation codes in that they do not all have the same synonymous codon block structure or the same number of stop codons as the standard code, we reasoned that these two structural features may provide an explanation for these observations.

The Ostrov codes are not required to maintain the synonymous codon block structure of the standard code, so they can have more or fewer split codon blocks than the standard code. In the set of the 194,481 Ostrov codes, the number of split codon blocks ranges from zero to four (Supp. Fig. S12). Increasing the number of split codon blocks decreases code robustness (*R* = *−*0.345*, p <* 2.2 *·* 10*^−^*^16^; Supp. Fig. S13A), due to the increase in the number of non-synonymous mutations. This causes an increase in landscape ruggedness (Fig. 5A and Supp. Tab. S12), because maladaptive valleys can form in the mutational spaces between synonymous codons of split codon blocks. While the relationship between the number of split codon blocks and the average fitness reached by random adaptive walks depends on population size (Supp. Tab. S14), the average fitness reached by the greedy adaptive walks consistently decreases as the number of split codon blocks increases (*R* = *−*0.261, *p <* 2.2 *·* 10*^−^*^16^, GB1; *R* = *−*0.214, *p <* 2.2 *·* 10*^−^*^16^, ParD-ParE2; *R* = *−*0.109, *p <* 2.2 *·* 10*^−^*^16^, ParD-ParE3; Fig. 5B and Supp. Tab. S13). Consistent with the growing number of local peaks, we also observe that the adaptive walks get on average shorter and their endpoints less predictable as the number of split codon blocks increases (Supp. Tab. S13). Interestingly, this effect remains even when restricting our analyses to codes that have the same robustness but differ in the number of split codon blocks (Supp. Fig. S14 and S15), showing that code robustness, as defined here, does not capture the full spectrum of effects mediated by changes in the number of split codon blocks.

**Figure 5:**
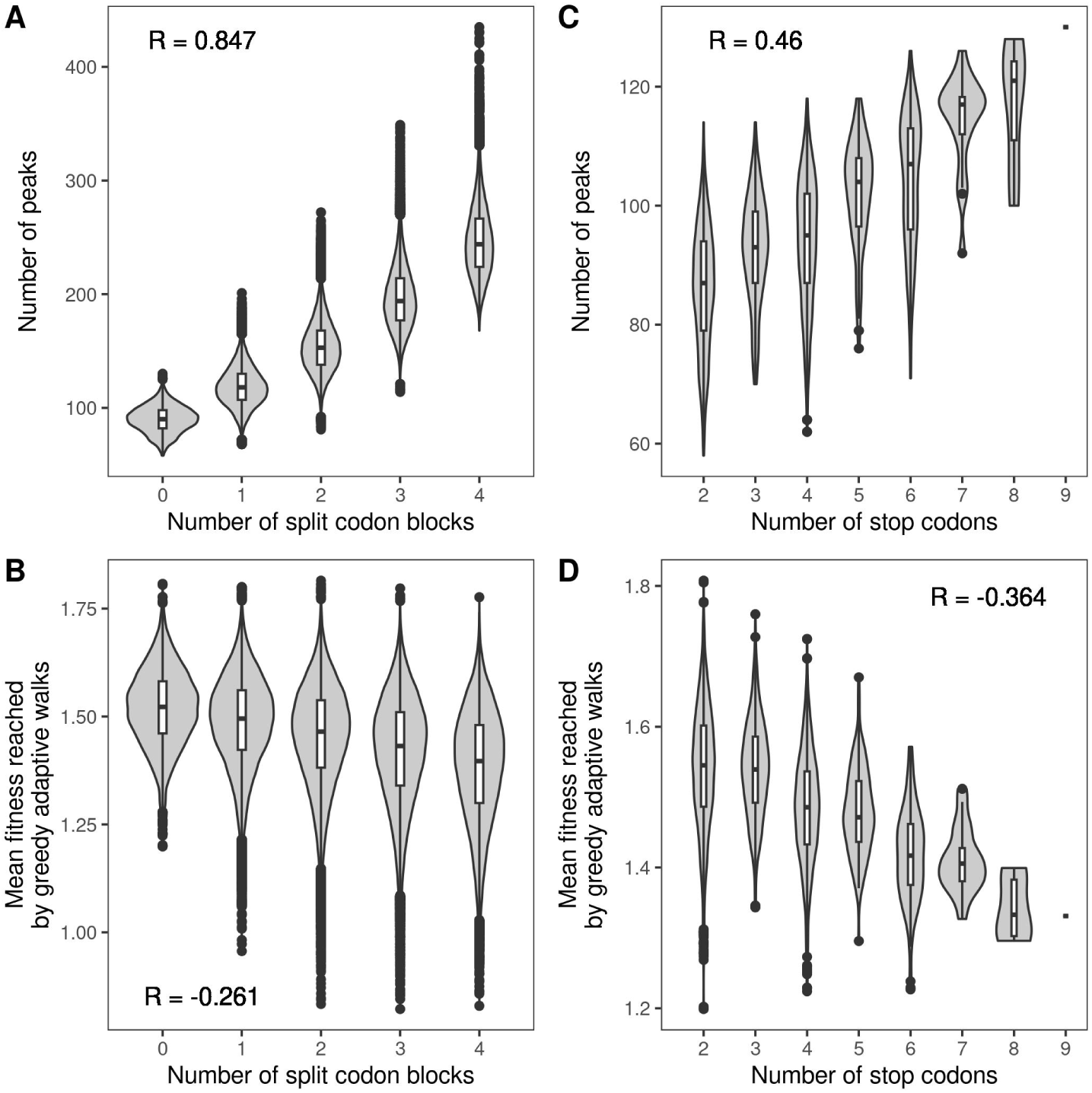
Additional code features influencing protein evolvability. Violin plots of the number of local peaks and the mean fitness reached by the greedy adaptive walks, shown in relation to (A, B) the number of split codon blocks in the 194,481 Ostrov codes and (C, D) the number of stop codons in the 3965 Ostrov codes with no split codon blocks. Data pertain to GB1. The violin plots show the distribution and the box-and-whisker plots the median, 25th and 75th percentile. The upper whisker extends from the top of the box to the largest value no further than 1.5-times the inter-quartile range, the lower whisker extends from the bottom of the box to the smallest value no further than 1.5-time the inter-quartile range. Data beyond the end of the whiskers are plotted individually.

The Ostrov codes can also have more or fewer stop codons that the standard code, which has three (UAG, UAA, and UGA). Because only the stop codon UAG has been freed for reassignment in the Ostrov codes, the minimum number of stop codons is two, whereas the maximum is nine, corresponding to the assignment of all freed codons to a termination signal (Supp. Fig. S16). Supp. Fig. S13B shows that increasing the number of stop codons tends to decrease code robustness (*R* = *−*0.160, *p <* 2.2 *·* 10*^−^*^16^), due to the increase in the number of nonsense mutations. Moreover, the number of stop codons is negatively correlated with the number of split codon blocks (*R* = *−*0.269, *p <* 2.2 *·* 10*^−^*^16^), because if a codon block is assigned to a stop signal, it cannot be part of a split codon block. Thus, when measuring the effect of the number of stop codons on landscape ruggedness or the outcomes of adaptive walks, one has to condition on a given number of split codon blocks. In the following, we report results for codes with 0 split codon blocks; results for other numbers of split codon blocks can be found in Supp. Tab. S15, S16, and S17. We observe that, among codes with a given number of split codon blocks, increasing the number of stop codons leads to an increase in the number of local peaks (Fig. 5C and Supp. Tab. S15), as well as decreased accessibility of the global peak (Supp. Tab. S15); the effect on epistasis is more complex (Supp. Tab. S15 and Supp. Note S1). Correspondingly, the average fitness reached by both the greedy and random adaptive walks decreases as the number of stop codons increases (greedy walks: *R* = *−*0.364, *p <* 2.2 *·* 10*^−^*^16^, GB1; *R* = *−*0.147, *p <* 2.2 *·* 10*^−^*^16^, ParD-ParE2; *R* = *−*0.183, *p <* 2.2 *·* 10*^−^*^16^, ParD-ParE3; Fig. 5D and Supp. Tab. S16 and S17). This is expected, as in our adaptive landscapes sequences containing stop codons are assigned a fitness value lower than any of the sequences without stop codons (Methods), reflecting the fact that the inclusion of a stop codon in an open reading frame causes the premature termination of translation and thus protein truncation, which is usually deleterious to protein function. We also observe that the greedy adaptive walks get shorter and less predictable as the number of stop codons increases (Supp. Tab. S16). Moreover, these effects of increasing the number of stop codons remain even among codes with the same robustness (Supp. Fig. S17 and S18). In sum, our analyses of all possible code rewirings under the codon compression scheme proposed by Ostrov et al. (2016) reveals additional code features influencing protein evolvability, namely the number of split codon blocks and the number of stop codons.

### 2.7 Design principles: Genetic codes enhancing and diminishing evolvability

We previously discussed how code robustness can be defined in terms of different amino acid properties, and showed that the amino acid properties most relevant to landscape topography are protein-dependent. Whereas for GB1, beta-sheet propensity and, to a lesser extent, hydrophobicity are key properties, alpha-helix propensity plays a more important role for ParD3. This suggests that a genetic code that promotes evolvability for one protein might not do so for another. Indeed, in our evolutionary simulations with the Ostrov codes, the mean fitness reached by the greedy walks is not strongly correlated across our three data sets (Supp. Tab. S18). Nonetheless, there is a small subset of codes that promote evolvability across all three data sets, and we reasoned that these may exhibit commonalities that could inform design principles for engineering genetic codes to promote evolvability across a diversity of proteins.

We therefore ranked each Ostrov code in descending order according to mean fitness reached in the evolutionary simulations, separately for each of the three data sets. Fig. 6A shows a Venn diagram of the top 20% of codes in each ranked list, revealing that 675 codes consistently rank in the top 20% of all three lists. We note that the standard genetic code is not a member of this set of consistently high-ranking codes, as it only ranks in the top 20% of codes for the ParD-ParE2 data set (Supp. Tab. S19). This shows that even relatively small changes to the standard genetic code can enhance protein evolvability.

**Figure 6:**
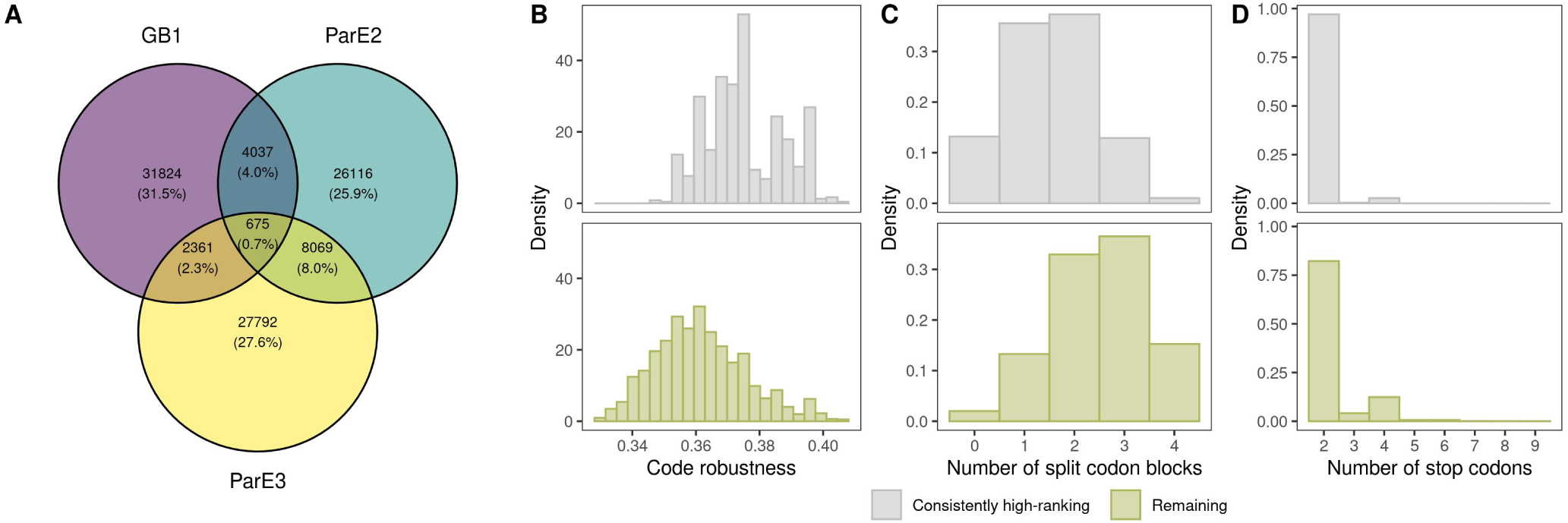
Design principles for enhancing evolvability. (A) Venn diagram of the top 20% of Ostrov codes, ranked according to mean fitness reached in the evolutionary simulations, for each of the three data sets. (B)-(D) Comparison of the properties of the 675 consistently high-ranking codes (three-way intersection in (A)) with the remaining 193,806 codes, in terms of (B) code robustness, (C) number of split codon blocks, and (D) number of stop codons.

We then compared these consistently high-ranking codes to the remaining 193,806 codes, in terms of robustness, number of split codon blocks, and number of stop codons. The consistently high-ranking codes have significantly higher robustness (*p <* 2.2 *·* 10*^−^*^16^, Welch two-sample t-test; Fig. 6B), fewer split codon blocks (*p <* 2.2 *·* 10*^−^*^16^, Welch two-sample t-test; Fig. 6C), and fewer stop codons (*p <* 2.2 *·* 10*^−^*^16^, Welch two-sample t-test; Fig. 6D). This suggests there are some basic design principles to engineering genetic codes that promote evolvability across a diversity of proteins. Specifically, minimize the number of split codon blocks and the number of stop codons, and assign amino acids to codon blocks such that point mutations cause only small changes to amino acid properties, using an aggregate measure of a diversity of amino acid properties (Pines et al., 2017).

To illustrate these design principles, Fig. 7A shows the genetic code with the highest robustness of the consistently high-ranking Ostrov codes (‘Evolvable Ostrov Code A’). It has a robustness of 0.41, the minimal number of zero split codon blocks, and the minimal number of two stop codons. Engineering this code in a living organism is in principle possible with existing technology, although it requires the reassignment of all seven freed codons, which is no small feat. In contrast, most experimental studies of rewired genetic codes only change the meaning of the UAG stop codon (Mukai et al., 2017). Bacterial strains containing no genomic TAG, as well as a variety of orthologous translation systems that decode UAG as a nonstandard amino acid are commercially available, so engineering a strain with reassigned UAG is relatively straightforward. There are two such codes among our consistently high-ranking codes, specifically those reassigning UAG to leucine (Fig. 7B, ‘Evolvable Ostrov Code B’) and alanine (Fig. 7C, ‘Evolvable Ostrov Code C’). These can be readily engineered with commercially-available recoded bacterial strains and plasmid-borne orthogonal translation systems. How do these codes compare to the standard genetic code in our evolutionary simulations? All three of them rank better in fitness on the GB1 and ParD-ParE3 landscapes, and Evolvable Ostrov Code B and C even rank better than the standard genetic code on the ParD-ParE2 landscape, even though on this landscape the standard genetic code outperforms 96.9% of the Ostrov codes (Supp. Tab. S19). We note that in the GB1 and ParD-ParE2 landscapes, the mean fitness reached under the standard genetic code, as well as Evolvable Ostrov Codes A, B, and C is even higher than when using genetic codes specifically designed for increased evolvability (Pines et al., 2017) (Supp. Tab. S19), even though they are much easier to engineer than those proposed by Pines et al. (2017). Evolvable Ostrov Codes B and C are thus promising candidates for easily engineerable genetic codes that are expected to provide a moderate increase in evolvability compared to the standard genetic code.

**Figure 7:**
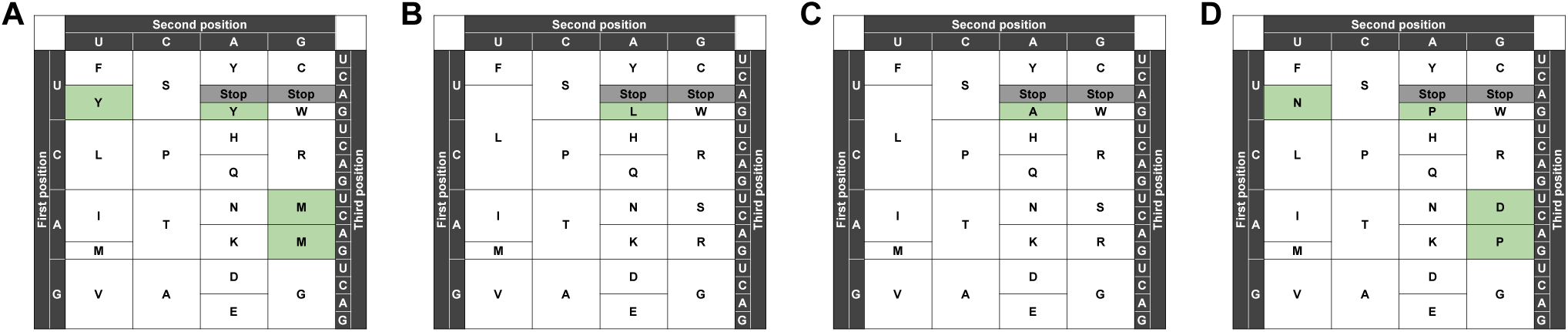
Examples of codes promoting (A-C) or diminishing (D) evolvability, identified based on their robustness, number of stop codons, and number of split codon blocks, as well as the results of the greedy adaptive walks. Changes compared to the standard genetic code are highlighted in green.

On the other side of the spectrum, there are 3,645 genetic codes that consistently rank among the bottom 20% of codes, according to mean fitness reached by the greedy adaptive walks; i.e., they consistently decrease evolvability. We note that this number is much higher than the number of consistently high-ranking codes (675), suggesting that decreasing evolvability is less data set-specific. In line with the previous results, we observe that the consistently low-ranking codes have lower robustness (*p <* 2.2 *·* 10*^−^*^16^, Welch two-sample t-test), more split codon blocks (*p <* 2.2 *·* 10*^−^*^16^, Welch two-sample t-test), and more stop codons (*p <* 2.2 *·* 10*^−^*^16^, Welch two-sample t-test) compared to the remaining codes (Supp. Fig. S19). However, unlike for the consistently high-ranking codes, it is not possible to optimize all of these design principles at the same time. For example, a genetic code where all free codon blocks are assigned to a stop signal will have the maximum possible number of stop codons (9), but it will also have the minimal number of split codon blocks (0). It is thus impossible to highlight one genetic code that would be expected to decrease evolvability the most based on these design principles. Instead, in Fig. 7D we show a genetic code that ranks among the bottom 2% of codes for all three data sets (mean fitness 1.032, ranking better than 0.35% of codes, GB1; mean fitness −0.193, ranking better than 0.51% of codes, ParD-ParE2; mean fitness −0.256, ranking better than 1.57% of codes, ParD-ParE3). This code, while having only two stop codons, has the maximum number of split codon blocks (4) and a robustness of 0.340, which is lower than 92.3% of the Ostrov codes. We again compared the level to which this code diminishes evolvability with codes specifically designed to slow down the rate of evolution (Calles et al., 2019) (Supp. Tab. S19). While reducing evolvability beyond the majority of the Ostrov codes, the mean fitness reached in adaptive walks using genetic code D is still much higher than when using the codes proposed by Calles et al. (2019) (mean fitness −1.78 and −0.87 for the ‘FS20’ and ‘RED20’ codes, respectively, proposed by Calles et al. (2019), vs. 1.03 for Code D, GB1 data set). However, we emphasize that, similar to the codes proposed by Pines et al. (2017), the codes proposed by Calles et al. (2019) require extensive genome recoding, such that the majority of codons are ‘null’, meaning they encode neither an amino acid nor a stop signal. We hope the design principles we have identified here will provide guidance for engineering genetic codes that significantly enhance or diminish evolvability, but remain within reach of current technology.

## 3 Discussion

The standard genetic code defines the rules of protein synthesis for nearly every life form on Earth (Knight et al., 2001). It imparts an extreme, “one in a million” level of error tolerance (Freeland and Hurst, 1998) that buffers the deleterious effects of infidelity in replication, transcription, and translation (Haig and Hurst, 1991; Freeland and Hurst, 1998), and provides a striking example of biological robustness at the heart of an essential cellular information processing system (Wagner, 2005). Despite decades of research on the origins and evolutionary implications of the standard genetic code (Crick, 1968; Woese, 1965; Knight et al., 1999; Koonin and Novozhilov, 2017), its influence on protein evolvability remained poorly understood (Freeland, 2002; Pines et al., 2017). Here, by computationally translating millions of mRNA sequences under hundreds of thousands of rewired genetic codes using experimental data for three proteins, we reveal that the robustness of the standard genetic code facilitates protein evolvability by rendering smooth adaptive landscapes upon which evolving populations readily find mutational paths to adaptation.

Prior theoretical work, limited by a lack of suitable data, has disagreed on whether the robustness of the standard genetic code hinders (Pines et al., 2017) or facilitates (Freeland, 2002) protein evolvability. These conflicting conclusions derive in part from a difference in the timescale of adaptation. Over short evolutionary timescales, for example where adaptation occurs via a single mutation, a robust genetic code may hinder evolvability by limiting the number and physicochemical diversity of amino acids accessible via point mutation (Pines et al., 2017). In contrast, over intermediate evolutionary timescales, where adaptation proceeds via a sequence of several mutations, a robust genetic code may facilitate evolvability by ensuring that missense mutations cause at most small changes to the physicochemical properties of amino acids (Freeland, 2002), which are less likely to be deleterious than large changes (Firnberg and Ostermeier, 2013). While single-step adaptation is undeniably important, as features emerging via a single mutation include e.g. antibiotic resistance (Jin and Gross, 1988; Manson et al., 2017; Woodford and Ellington, 2007), many other adaptations require a complex interplay of several mutations (Karageorgi et al., 2019; Meyer et al., 2012; Tenaillon et al., 2012). For example, in directed protein evolution experiments, on average five mutations are needed per order-of-magnitude improvement in catalytic efficiency (Goldsmith and Tawfik, 2017). Our results, based on experimental data, suggest that over such timescales, increasing code robustness indeed enhances protein evolvability. However, this enhanced evolvability comes at the cost of an increased time to convergence, with organisms using robust genetic codes reaching higher fitness peaks on average, but using more mutations. This trade-off should be kept in mind when designing directed protein evolution experiments that utilize synthetic organisms with non-standard genetic codes (Drienovska and Roelfes, 2020; Hammerling et al., 2014, 2016; Tack et al., 2018).

Over even longer timescales, evolvability is influenced by the topology of the set of high-fitness variants, which we refer to here as a genotype network (Lipman et al., 1991; Schuster et al., 1994; Wagner, 2008). We studied the topology of this network under the standard genetic code, as well as under rewired genetic codes with exceptionally low and high robustness. We did so by visualizing the approximate evolutionary distances between high-fitness variants at mutation-selection-drift balance (McCandlish, 2011). The influence of a genetic code on genotype network topology was immediately apparent from the diversity of shapes these networks adopt under different codes (Fig. 4). In terms of evolvability, both robust and non-robust genetic codes produced fitness peaks isolated from the bulk of high-fitness sequences, as well as long winding paths between high-fitness protein sequences that differ by only 1 or 2 amino acids, driven by the interaction of the genetic code with the geometry of the fitness landscape at the amino acid level. However, we observed that the robust codes, as well as the standard genetic code, tend to provide stronger connectivity between high-fitness sequences, often exhibiting grid-like structures that facilitate the independent evolution of sets of mutations. This reduces dependence on the order in which mutations accumulate, thus increasing the number of mutational paths between high-fitness sequences. In contrast, under non-robust codes, the high-fitness sequences tended to form linear or tree-like structures, which increase dependence on the order in which mutations accumulate and thus limit the number of mutational paths between high-fitness sequences.

Landscape ruggedness has long been viewed as an impediment to adaptation (Wright, 1932), with implications for a diversity of evolutionary phenomena, including the evolution of genetic diversity, sex, and reproductive isolation (Szendro et al., 2013). As such, ruggedness has been studied extensively in both theoretical (Kauffman and Levin, 1987; Kingman, 1977; Kauffman and Weinberger, 1989) and empirical (de Visser and Krug, 2014; Wu et al., 2016; Schenk et al., 2013; Gong et al., 2013; Jiménez et al., 2013; Sarkisyan et al., 2016; Aguilar-Rodríguez et al., 2017; Olson et al., 2014; Hartman et al., 2019) adaptive landscapes. It has also been used as a proxy for evolvability (Payne and Wagner, 2019). The intuition is that ruggedness frustrates evolvability by blocking “uphill” mutational paths to the global adaptive peak, thus limiting the ability of mutation to bring forth adaptive phenotypic variation. Our results confirm this intuition in the context of rewired genetic codes, in that adaptive walks tend to achieve higher fitness on smooth landscapes caused by more robust codes than on rugged landscapes caused by less robust codes. However, this is not solely attributable to an increase in the mutational accessibility of the global peak. Rather, in the landscapes we study, as ruggedness decreases, a positive correlation emerges between the height of a peak and its basin of attraction. This causes adaptive walks to preferentially converge on a small number of high-fitness peaks in less rugged landscapes, and more uniformly to all peaks in more rugged landscapes. As such, simplistic measures of landscape ruggedness based solely on the number of peaks may be an insufficient proxy for evolvability (Payne and Wagner, 2019) or for predicting evolutionary dynamics (Lässig et al., 2017).

An additional factor influencing the accessibility of the global adaptive peak is its size. By randomly permuting amino acids amongst synonymous codon blocks, we created landscapes that vary significantly in the number of mRNA sequences that translate to the highest-fitness protein sequence. In comparing the outcomes of evolutionary simulations on these landscapes, we observe that protein sequences encoded by a large number of mRNA sequences are easier to evolve than equally fit sequences encoded by fewer mRNA sequences. In other words, amino acid sequences encoded by a large number of mRNAs are more “findable,” because they occupy a larger fraction of genotype space (McCandlish, 2013; Dingle et al., 2021; Schaper and Louis, 2014). This observation implies that amino acids encoded by a large number of codons, such as serine or leucine, should be relatively more abundant in protein sequences than amino acids encoded by few codons, such as methionine or tryptophan. Indeed, across the tree of life, there is a positive correlation between the abundance of an amino acid and its number of constituent codons (King and Jukes, 1969; Gilis et al., 2001), and as early as 1973, Jack L. King attributed this correlation to differences in amino acid “findabilities” caused by the structure of the standard genetic code (King, 1973). Our results generalize this observation to non-standard genetic codes, and suggest that if life had converged on a different standard code, the amino acid composition of proteins would likely be very different from the one we know. Such variation in the proteomic abundance of amino acids may already be apparent in the proteomes of organelles and organisms that use non-standard genetic codes in nature (Knight et al., 2001; Ambrogelly et al., 2007; Shulgina and Eddy, 2021), and may emerge in directed laboratory evolution experiments that use synthetic organisms with non-standard genetic codes (Liu and Schultz, 2010; de la Torre and Chin, 2021; Zürcher et al., 2022). If so, these systems may provide empirical support for entropic arguments of adaptation (Schaper and Louis, 2014; Dingle et al., 2021).

There are several caveats to the results presented here. First, because there are so few combinatorially-complete data sets measuring a quantitative phenotype, our conclusions are based on only three empirical adaptive landscapes. However, the three data sets we use differ in several important aspects – the level of ruggedness of the corresponding landscape under the standard code, the location of the screened residues in the protein, as well as the assayed phenotype – which supports the generality of our findings. Second, all three landscapes pertain to just a small number of sites within a larger protein, so it is not possible to understand how sequence variants at these sites interact with other sites in the protein. Third, while the rate of cell growth, a common measure of bacterial fitness (Wiser and Lenski, 2015), was measured for the ParD-ParE2 and ParD-ParE3 data sets, in the case of GB1 the screened phenotype is the relative binding affinity of the protein to immunoglobulin. How this protein phenotype relates to organismal fitness is not immediately apparent. Even so, a large body of literature attests to the power of such quantitative phenotypes in teaching us about protein evolvability (Tokuriki and Tawfik, 2009). Fourth, the error tolerance of a genetic code, standard or non-standard, is influenced by mutation bias and codon usage (Freeland and Hurst, 1998; Radványi and Kun, 2021) as they make some mutations more likely than others. While mutation bias and codon usage may influence peak accessibility in adaptive landscapes (Cano and Payne, 2020), they do not affect landscape topography, which is why we have not considered these effects here. We hope that in the future it will become possible to overcome these caveats and confirm our results, both theoretically, as more combinatorially complete data sets become available, and experimentally, by comparing the dynamics and outcomes of laboratory evolution experiments with proteins and organisms that use different genetic codes.

Such experiments are becoming more broadly accessible, as a diversity of recoded organisms and plasmid-borne orthogonal translation systems are now commercially available. Moreover, these experiments are becoming increasingly scalable. For example, Zürcher et al. (2022) have recently engineered bacterial strains with as many as 16 different genetic codes. Understanding the relationship between code structure and evolvability is therefore highly topical, as the future in which synthetic organisms with non-standard genetic codes are utilized in science and in industry (Li and Liu, 2014; Jin et al., 2019), for example to accelerate directed evolution experiments (Romero and Arnold, 2009; Pines et al., 2017; Zürcher et al., 2022), to achieve bio-containment (Marliere, 2009; Kubyshkin and Budisa, 2017; Calles et al., 2019; Zürcher et al., 2022; Nyerges et al., 2023; Fujino et al., 2020), or to produce drugs (Romesberg, 2023; Sun et al., 2014; Ptacin et al., 2021) is now tangibly close. We have identified general design principles, as well as a few concrete candidate codes, that are expected to increase evolvability beyond that of the standard genetic code. These are compatible with the 57-codon *E. coli* genome reported by Ostrov et al. (2016), and could thus be engineered in the lab using existing technology. Our analyses with this codon compression scheme explored all 194,481 genetic codes that reassign one or more of the freed codon blocks, assuming that the whole synonymous codon block needs to be assigned to one amino acid. However, this assumption might not be needed, as shown by a recent refactoring of the genetic code, in which the UCG and UCA codons were reassigned independently, even though the naturally occurring 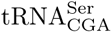 and 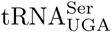 are not specific to their anticodon (Zürcher et al., 2022). In the context of the 57-codon *E. coli* genome, it might thus be possible to change the meaning of up to 7 codons independently, leading to a staggering 21^7^ *≈* 1.8 *·* 10^9^ possible code rewirings. Together with other codon compression schemes (Fredens et al., 2019; Lajoie et al., 2013; Lau et al., 2017; Wang et al., 2016), the space of possible code rewirings available today is practically infinite, and will continue to grow as larger-scale rewirings become feasible.

Building upon our results, there are several directions for future research. First, the advances in biotechnology discussed above now enable experimental tests of the relationship between genetic code robustness and evolvability, as recently proposed by Zürcher et al. (2022). Do robust genetic codes indeed lead to larger improvements of protein function in directed evolution experiments? And are organisms with robust genetic codes better able to adapt to changing environmental conditions? Second, in this paper we have worked only with rewired genetic codes, i.e., codes that change the mapping between codons and amino acids, but we have not considered expanded genetic codes, i.e., codes that include a 21st, non-standard amino acid. Examples of expanded genetic codes can be found both in nature, with the expansion of the standard genetic code to include selenocysteine in many different organisms across the tree of life (Gladyshev and Kryukov, 2001; Hatfield and Gladyshev, 2002) and the addition of pyrrolysine in methanogenic archea (Brugère et al., 2018), as well as in the lab, with recoded genomes that can in principle incorporate any non-standard amino acid (Chin, 2014; Xie and Schultz, 2006; de la Torre and Chin, 2021; Liu and Schultz, 2010; Lajoie et al., 2013; Romesberg, 2023; Dumas et al., 2015; Nödling et al., 2019). While a small number of evolutionary experiments using organisms with expanded genetic codes have been reported (Hammerling et al., 2014, 2016; Tack et al., 2018; Thyer et al., 2018), how the addition of a 21st non-standard amino acid influences protein and organismal evolvability is not yet fully understood. This question could be addressed experimentally by generating combinatorially-complete data for all 21*^L^* sequence variants, using a diversity of 21st non-standard amino acids. Indeed, because the ParD assay is based on bacterial cell growth, it should be directly compatible with off-the-shelf recoded organisms, such as the recoded *E. coli* genome engineered by Lajoie et al. (2013). Moreover, the GB1 assay may be amenable to cell-free translation systems that allow for the incorporation of additional amino acids (Shimizu et al., 2001; Hartman et al., 2007). How increasing the number of amino acids in a genetic code influences evolvability could also be addressed theoretically, for example by subsampling combinatorially-complete data to contain fewer than 20 amino acids. Such analyses could provide answers to fundamental questions like “Why are there only 20 proteinogenic amino acids even though the standard genetic code theoretically has the capacity to encode up to 64?” (Hayes, 1998). Finally, our results may shed light on the evolution of the naturally-occurring deviations of the standard genetic code (Knight et al., 2001; Ambrogelly et al., 2007; Shulgina and Eddy, 2021). It has been suggested that the changes observed in these alternative genetic codes might be adaptive because they increase code robustness (Błażej et al., 2019) and, based on our results, such an increase in code robustness may promote evolvability. It should be possible to test whether organisms using alternative genetic codes indeed show signs of increased evolvability, e.g., by comparing dN/dS ratios in organisms using different genetic codes.

In conclusion, our results suggest that the robustness of a genetic code not only buffers against replication and translation errors, but also facilitates the generation of adaptive phenotypic variation. Such robustness is therefore essential to both life’s survival and its advancement.

## 4 Methods

### Data processing

We estimated the fitness of each measured amino acid variant following Rubin et al. (2017). For GB1, the fitness of variant *v* is equal to

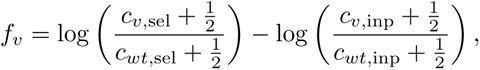

where *c_v,_*_sel_ is the count of variant *v* in the sample after selection for binding immunoglobulin, *c_wt,_*_sel_ is the count of the wild type (VDGV) in the sample after selection for binding immunoglobulin, *c_v,_*_inp_ is the count of variant *v* in the input sample, and *c_wt,_*_inp_ is the count of the wild type in the input sample. The variance of the estimate is equal to

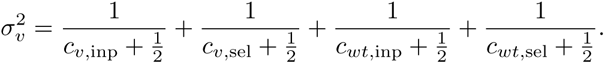

For ParD-ParE2 and ParD-ParE3 there are two replicates for each measurement. For each variant and each replicate we computed the fitness and variance as described above for GB1. The final fitness of variant *v* is then the weighted average of the two replicates, with weights given by the inverse of the corresponding variance:

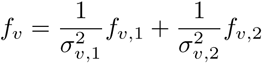

and the variance is computed as

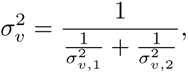

where by *f_v,i_*and 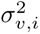 we denote the fitness and variance, respectively, of *i*-th replicate.

Based on these raw fitness estimates and variances for observed variants, we inferred the full adaptive landscape as the maximum *a posteriori* estimate under empirical variance component regression, an empirical Bayes modeling framework that naturally incorporates all orders of genetic interaction (Zhou et al., 2022).

### Additive models of the data

To assess the ruggedness of the three adaptive landscapes, we attempted to model each data set using an additive model. In particular, we assumed that the final fitness is a linear combination of contributions of individual sites. Formally,

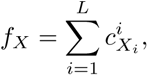

where *f_X_* is the fitness of variant *X*, 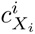 is the contribution of amino acid *X_i_* at position *i* of the protein towards fitness, and *L* is the length of the protein (4 for GB1, 3 for ParD). The coefficients 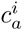 were found by fitting a linear regression model to the fitness values.

### Constructing adaptive landscapes

To construct an adaptive landscape, we consider the set of all possible mRNA sequences of length 12 (for GB1, so that they encode 4 amino acids; there is 4^12^ = 16, 777, 216 such sequences) or 9 (for ParD-ParE2 and ParD-ParE3, encoding 3 amino acids; 4^9^ = 262, 144 sequences), respectively. We represent each sequence as a vertex in a mutational network. Two vertices are connected with an edge if the Hamming distance of the corresponding mRNA sequences is 1, i.e., the two sequences differ by a single point mutation. This underlying network is the same for all genetic codes.

For each genetic code, we then assign an “elevation” to each vertex, equal to the fitness of the sequence, translated using a given genetic code. Sequences containing stop codons are assigned an arbitrary elevation lower than the fitness of any sequence not containing stop codons (we used a value of −100, but the precise value is not relevant for the analyses presented here).

### Code robustness

We define code robustness as the proportion of single-nucleotide substitutions that do not change the physicochemical properties of amino acids. We divided amino acids into 7 physicochemical groups following Pines et al. (2017) (Supp. Fig. S3): acidic (D, E); aliphatic (A, I, L, V); aromatic (F, W, Y); basic (H, K, R); glycine (G); polar (C, M, N, Q, S, T); and proline (P). We considered mutations from an amino acid to a stop codon or vice versa as a change in physicochemical properties, whereas we did not consider mutations among stop codons as a change in physicochemical properties.

### Number of adaptive peaks

Intuitively, a local peak is a sequence whose fitness is higher than the fitness of any of its neighbors. In our case, due to the degeneracy of the genetic code, several vertices often have the same elevation and, moreover, those vertices will usually be connected; peaks are thus usually plateaus rather than a single vertex. Formally, we define a local peak as a set of vertices that (1) are connected in the genotype space, (2) all have the same elevation, and (3) whose neighbors are either part of the set or have a lower elevation.

### Epistasis analysis

A square is a quadruplet of sequences that contains a ‘wild type’ sequence, two of its one-mutant neighbours, and the corresponding double mutant. In the following, we denote by *f*_00_ the fitness of the wild type, by *f*_01_ and *f*_10_ the fitness of the two single mutants, and by *f*_11_ the fitness of the double mutant. The ‘mutational effect’ of a given mutation is denoted by Δ*f*, e.g., Δ*f*_00_*_→_*_10_ = *f*_10_ *− f*_00_ is the change in fitness caused by mutating the wild type sequence to one of the single mutants. We say that there is no epistasis if

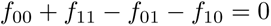

Because in our case the fitness values are real numbers, this only happens if at least two of the mutations are synonymous. The square is classified as having magnitude epistasis if

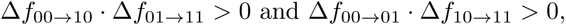

i.e., the effects of both mutations have the same sign (increase fitness or decrease fitness) regardless of the genetic background. Similarly, the square is classified to have reciprocal-sign epistasis if

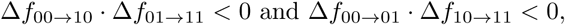

i.e., the effects of both mutations have opposite signs in different genetic backgrounds. The remaining cases, i.e.,

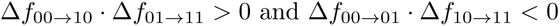

and

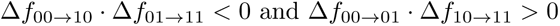

are classified as simple-sign epistasis: the sign of one of the mutations is the same in the different backgrounds, whereas the sign of the second mutation changes in the different backgrounds.

Due to the size of the genotype networks, listing all squares is computationally prohibitive. Thus, we randomly sampled 1,000,000 squares by first sampling a random sequence and then sampling two random mutations at two different positions in the sequence.

### Mutational accessibility of the global peak

We define the mutational accessibility of the global peak as the probability that, picking a random sequence not containing stop codons and a random direct path from the sequence to the global peak, the chosen path is accessible, i.e., the fitness increases monotonically along the path. In our case, the global peak is composed of several mRNA sequences; we define direct paths as those paths that reach any of the global peak sequences in the smallest possible number of steps. For example, considering the standard genetic code and the ParD-ParE3 data set, the global peak consists of 4 sequences: GAU UGG GAA, GAU UGG GAG, GAC UGG GAA, and GAC UGG GAG (all translate to DWE). Starting from sequence GAU UGG AUG (DWM), there are two direct paths to the global peak: GAU UGG AUG - GAU UGG **G**UG - GAU UGG G**A**G and GAU UGG AUG - GAU UGG A**A**G - GAU UGG **G**AG. Notice that in both cases the end point of the direct paths is only one of the 4 global peak sequences, since reaching the other 3 sequences in the global peak would require more than 2 mutations (and hence those paths are not considered direct). Technically, we computed the number of direct mutational paths from each vertex and their accessibility using breadth-first search.

### Artificial inflation of landscape ruggedness

We artificially inflated the ruggedness of the GB1 landscape by (1) increasing the number of local peaks, or (2) increasing the prevalence of reciprocal-sign epistasis. We then assessed the mutational accessibility of the global peak under the assumption of the standard genetic code. For each level of ruggedness inflation we created 100 adaptive landscapes, as described below.

To increase the number of local peaks, we chose a given number (ranging from 1 to 10,000) of protein sequences at random and set their fitness to such a value that it ensured that the corresponding region of the genotype network would form a local peak. In particular, we used a value of 2.5, which is halfway between the fitness value of the global peak (WWLA, fitness 2.52) and the second-best binding sequence (FYAA, fitness 2.48). Even when changing the fitness value of the same number of protein sequences, the number of local peaks in the resulting landscapes varies slightly, due to three reasons: (1) the original landscape contains 115 local peaks, some of which might cease to be local peaks if a neighboring sequence is artificially elevated; on the contrary, if a local peak is chosen and its elevation increased, the number of local peaks in the landscape does not change; (2) protein sequences containing the amino acid serine, which is encoded by the split codon block, are encoded by two disconnected regions in the genotype network, and artificially increasing their fitness thus creates two local peaks instead of one; (3) if two chosen sequences are neighbors in the genotype space, the corresponding mRNA sequences form one large plateau, and thus only one local peak is created instead of two.

To artificially increase the prevalence of reciprocal-sign epistasis, we randomly sampled an mRNA sequence of length 12 (i.e., encoding 4 amino acids), making sure it did not translate to the global peak sequence (WWLA) and the translation did not contain any stop codons. We then sampled two mutations such that they happened in different positions of the sequence, they were non-synonymous, both alone and in combination, and none of the single mutants or the double mutant contained any stop codons. We further required that none of the single mutants translated to the global peak sequence. We then permuted the fitness values of the corresponding protein sequences so that the double mutant had the highest value, the wild type the second highest, and the two single mutants the two lowest values. Since permuting the fitness values of a quadruplet of protein sequences changes the shape of many squares, it again holds that even among landscapes in which the same number of squares was changed the proportion of different types of epistasis is variable. The number of squares that were artificially forced to show reciprocal-sign epistasis ranged between 1 and 100,000.

### Analysis of the physicochemical properties of amino acids

We downloaded the set of 566 different descriptors of amino acids from the AAindex database, version 9.2 (Kawashima et al., 1999; Kawashima and Kanehisa, 2000). Of those, 13 contained at least one missing value; we discarded those, so the final set contained 553 different amino acid properties. These properties were previously divided into 4 categories: “alpha and turn propensity”, “beta propensity”, “hydrophobicity”, and “other” (Bartonek et al., 2020).

For each property *p* and each of our 100,000 amino acid permutation codes we computed the mean absolute change in the property under genetic code *c*, MAC(*p, c*), as

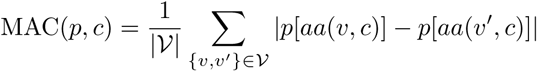

where *V* is the set of all codon pairs that are one substitution away from each other (excluding pairs that contain at least one stop codon), *aa*(*v, c*) is the amino acid encoded by codon *v* in genetic code *c*, and *p*[*a*] is the value of property *p* for amino acid *a*. MAC(*p, c*) quantifies the sensitivity of code *c* with respect to amino acid property *p*: if MAC(*p, c*) is large, it means that single-nucleotide substitutions tend to cause large changes in property *p*; if MAC(*p, c*) is low, single-nucleotide substitutions tend to preserve property *p*.

For each property *p* we then computed Pearson’s correlation between the sensitivities of our amino acid permutation codes with respect to property *p* and the different measures of landscape ruggedness:

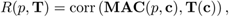

where **MAC**(*p,* **c**) is the vector of the values of MAC(*p, c*) for the set of 100,000 amino acid permutation codes and **T**(**c**) is the vector of values of certain landscape ruggedness measure (e.g., number of peaks) for these codes. For the mutational accessibility of the global peak, we only considered codes that preserve the size of the global peak, relative to the standard genetic code, and under which the global peak consists of a single connected region in the genotype space.

We determined the significance of each *R*(*p,* **T**) by comparison with a null distribution calculated from 1,000,000 randomly generated amino acid ‘properties’: We generated 1,000,000 null amino acid ‘properties’ by uniformly sampling 20 random numbers between 0 and 1. For each such null property *p*_null_, we computed **MAC**(*p*_null_, **c**) and *R*(*p*_null_, **T**) as described above. The significance of the true correlation coefficient is then the proportion of these 1,000,000 null correlation coefficients that are more extreme than the true value. We then corrected the significance values for each landscape ruggedness measure for multiple testing using Benjamini-Hochberg correction (Benjamini and Hochberg, 1995).

The enrichment of a certain category among the statistically significant properties (i.e., properties for which the p-value of the correlation coefficient is lower than 0.05 after correction for multiple testing) was tested using a one-tailed binomial test.

### Greedy adaptive walks

We simulated greedy adaptive walks on the landscape in which the most fit of the 1-mutant neighbors is fixed in every step, until a global or local peak is reached. However, the degeneracy of the genetic code means that the fitness values in the landscape are not unique, as all mRNA sequences encoding the same protein share the same fitness. The ‘most fit’ neighbor thus does not have to be uniquely defined, e.g. because there are several possible mutations that lead to the same fitness increase, or because a neutral plateau must be crossed before new adaptive variants may be generated. If this happens, we retain all sequences with the highest fitness; we then explore all of their 1-mutant neighbors and choose the fittest one(s) of those, etc.

We initiated the walks in all possible sequences not containing any stop codons.

### Weak mutation adaptive walks

The random walks were initiated in a randomly chosen mRNA sequence. In each subsequent step a random single-nucleotide mutation was suggested and accepted with probability

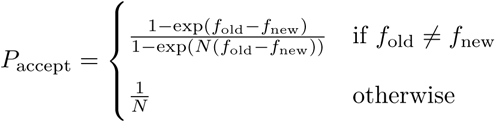

where by *f*_old_ we denote the fitness of the current genotype, *f*_new_ the fitness of the proposed genotype, and *N* is the population size. This corresponds to the exact fixation probability under the Moran process.

We always ran 100,000 adaptive walks of 500 steps for *N ∈ {*10; 100; 10, 000; 1, 000, 000*}*.

### Entropy of the distribution of reached peaks

For the greedy adaptive walks, we compute the entropy of the walks’ targets as

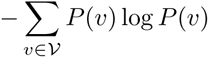

where *V* is the set of all endpoints of the greedy walks (i.e., the set of all adaptive peaks) and *P* (*v*) denotes the proportion of greedy walks that terminate on peak *v*.

### Visualization of the GB1 landscape under rewired genetic codes

We used the visualization method as previously described (McCandlish, 2011). Briefly, we construct a model of molecular evolution where a population evolves via single nucleotide substitutions and the rate at which each possible substitution becomes fixed in the population is related to its relative selective advantage or disadvantage. Specifically, the rate of evolution from sequence *i* to any mutationally adjacent sequence *j* is given by

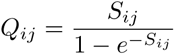

where *S_ij_*is the scaled selection coefficient (population size times the selection coefficient of *j* relative to *i*) and the total leaving rate from each sequence *i* is given by

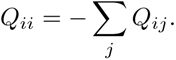

In this context, we assume that the selection coefficient between sequences *i* and *j* is proportional to the difference in log-enrichment scores or fitness *f_j_ − f_i_* and therefore *S_ij_* = *c*(*f_j_ − f_i_*), where *c* controls the strength of selection. For all analyses presented here, we used the simple choice of c=1, which for the standard genetic code gives a mean fitness at stationarity equal to 0.055, similar to the wild-type sequence VDGV (which by definition has fitness 0). Given the rate matrix *Q*, we then construct the visualization by using the subdominant right eigenvectors *r_k_* associated with the smallest magnitude non-zero eigenvalues *λ_k_* of this rate matrix as coordinates for the low dimensional representation of the landscape, where each such coordinate defines one of the “diffusion axes” used in the visualization. This visualization reflects the long-term barriers to diffusion in sequence space and clusters in the representation correspond to sets of initial states from which the evolutionary model approaches its stationary distribution in the same way. Thus, multi-peaked fitness landscapes appear as broadly separated clusters with one peak in each cluster. Moreover, by scaling the axes appropriately, as is done here,

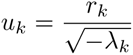

these axes *u_k_* have units of square-root of time, where time is measured in the expected number of neutral substitutions for a completely neutral sequence. In particular, using these coordinates *u_k_*, the squared Euclidean distance between arbitrary sequences *i* and *j* equals the sum of the expected time *H_ij_* to evolve from *i* to *j* and the expected time *H_ji_* to evolve from *j* to *i*, i.e.:

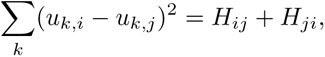

Using the first several *u_k_* (i.e. *u*_1_ and *u*_2_ for a 2-dimensional representation or *u*_1_, *u*_2_ and *u*_3_ for a three dimensional representation) optimally preserves the above relation in a principal components sense (see McCandlish (2011) for details).

## Acknowledgements

This work was funded by Swiss National Science Foundation grants PP00P3_202672 and 310030_192541 (J.L.P.) and NIH grant R35GM133613, an Alfred P. Sloan Research Fellowship, and additional funding from the Simons Center for Quantitative Biology at Cold Spring Harbor Laboratory (D.M.M.). We thank Macarena Toll-Riera for discussions, Václav Rozhoň for help with algorithm design and implementation, and Thuy-Lan Lite for kindly providing the counts data needed for the construction of the ParD-ParE2 and ParD-ParE3 landscapes.

---

**Figure S1:**
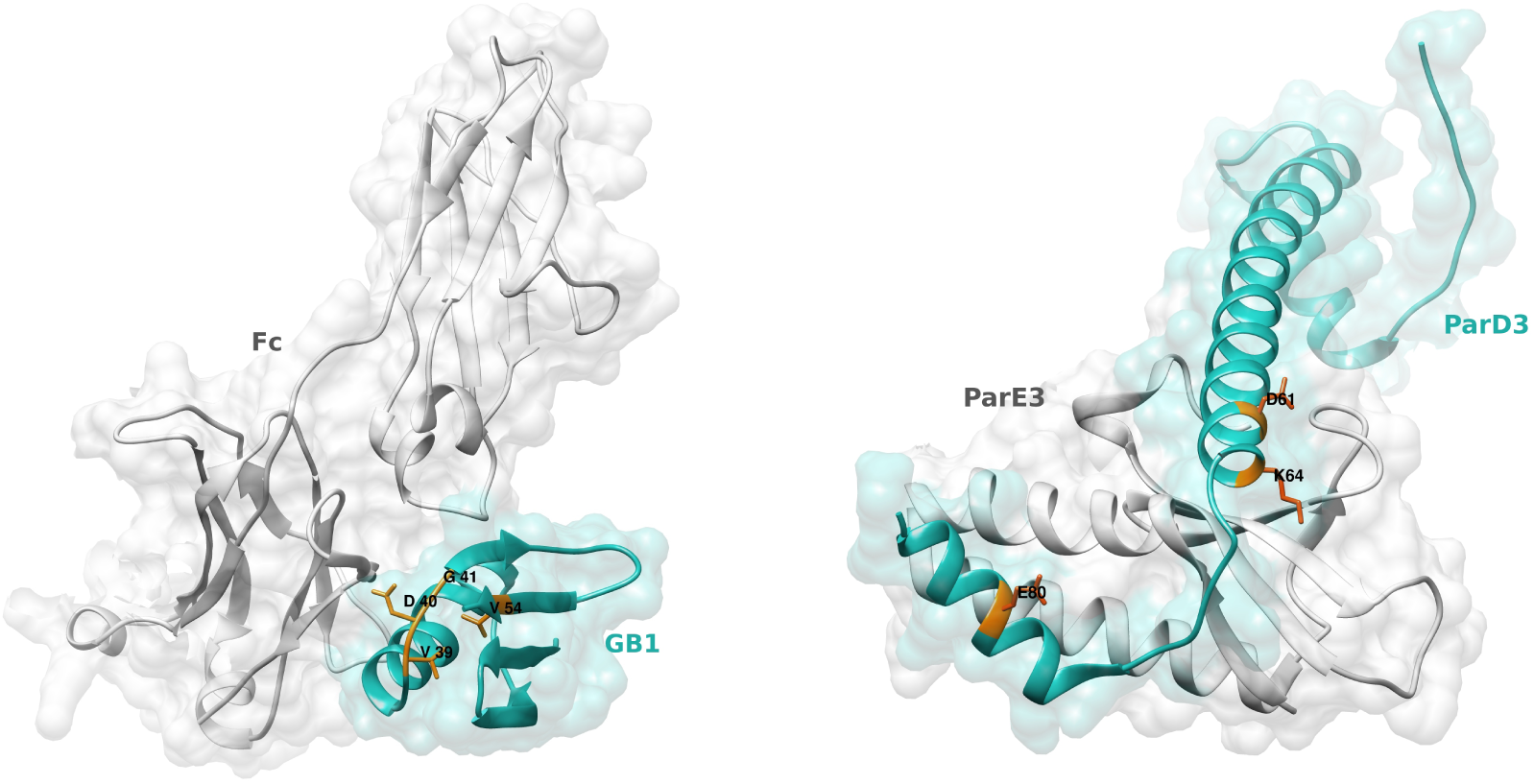
Structures of the (left) GB1 and (right) ParD3 proteins, in complex with their corresponding ligands (Fc domain of IgG for GB1; ParE3 for ParD3). The residues used to build the adaptive landscapes are highlighted in orange. PDB IDs: 1FCC for GB1, 5CEG for ParD3.

**Figure S2:**
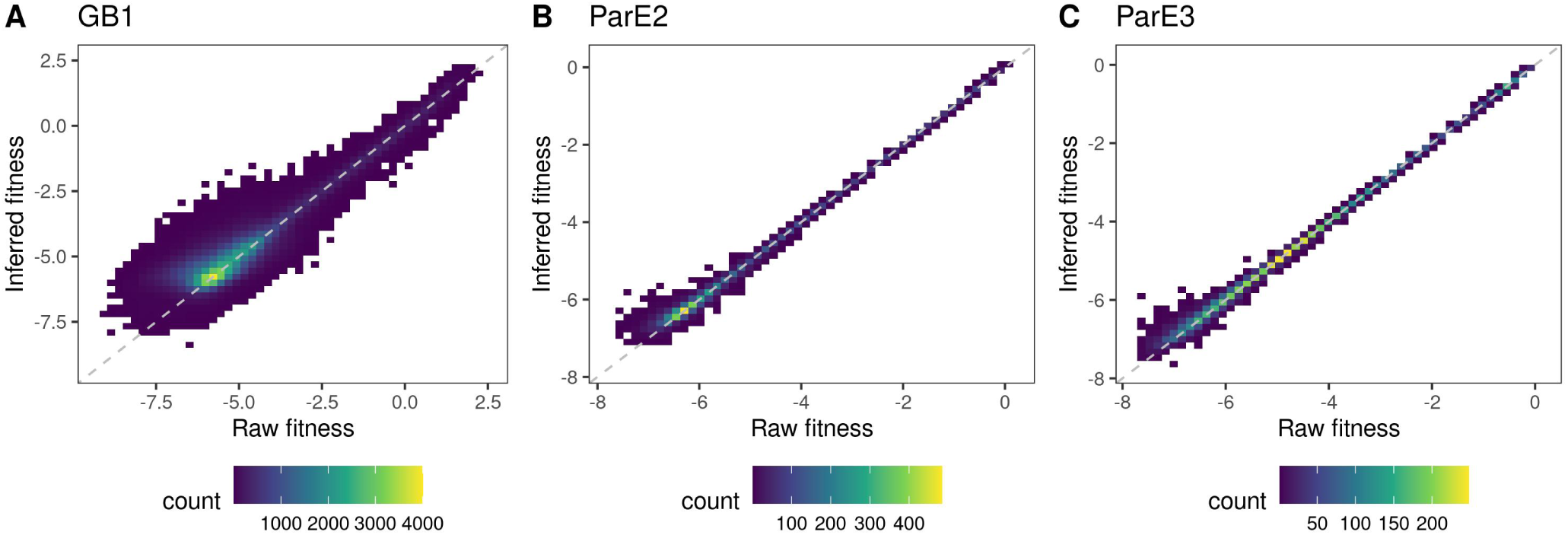
Density plot of the raw fitness values and the fitness values inferred using empirical variance component regression (Methods) (Zhou et al., 2022) for the (A) GB1, (B) ParD-ParE2, and (C) ParD-ParE3 data sets.

**Figure S3:**
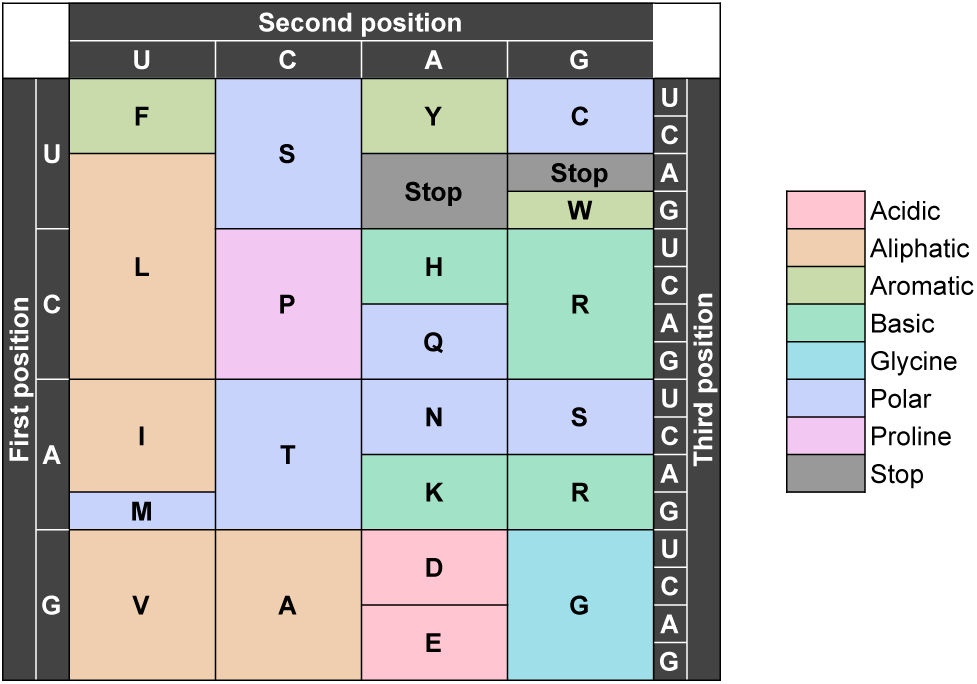
Codon table for the standard genetic code, with codons colored based on the physicochemical property of the encoded amino acid. Classification into physicochemical properties taken from Pines et al. (2017).

**Figure S4:**
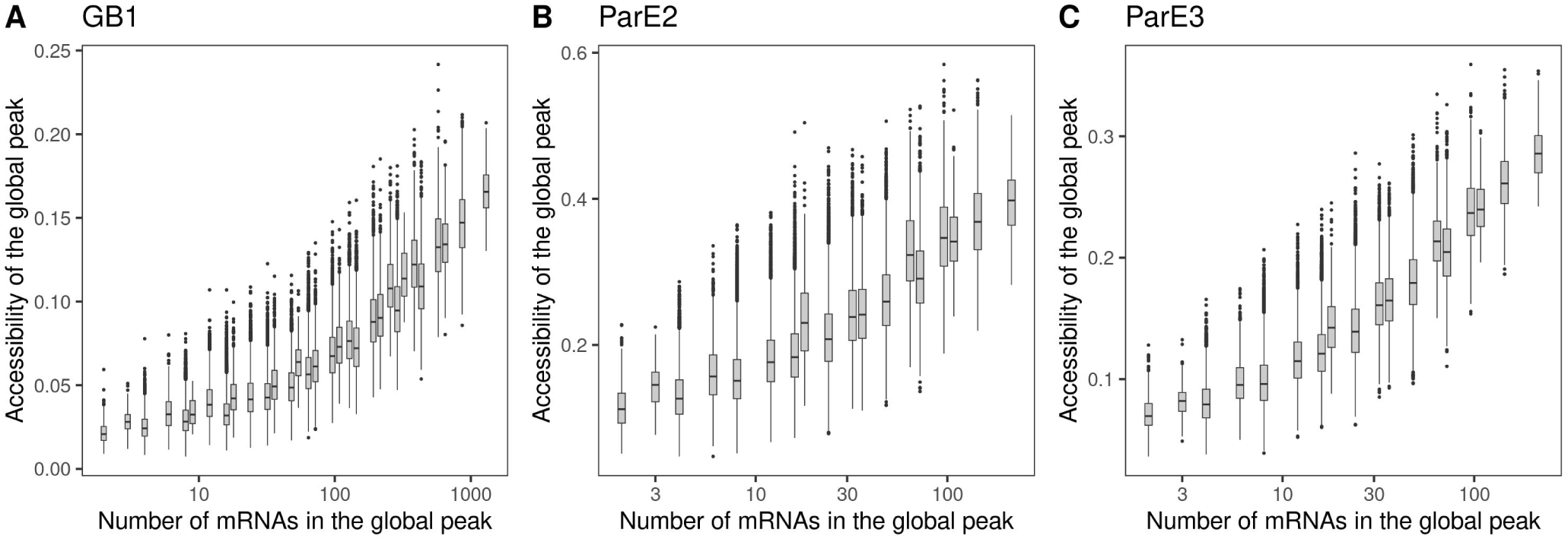
Accessibility of the global peak in relation to its size for the (A) GB1, (B) ParD-ParE2, and (C) ParD-ParE3 landscapes. Mutational accessibility is measured as the proportion of randomly chosen direct paths to the global peak that are accessible, meaning that fitness increases monotonically along the path. Peak size is measured as the number of mRNA sequences encoding the protein with the maximum fitness value. Data pertain to the 100,000 amino acid permutation codes. The box-and-whisker plots show the median, 25th and 75th percentile. The upper whisker extends from the top of the box to the largest value no further than 1.5-times the inter-quartile range, the lower whisker extends from the bottom of the box to the smallest value no further than 1.5-time the inter-quartile range. Data beyond the end of the whiskers are plotted individually.

**Figure S5:**
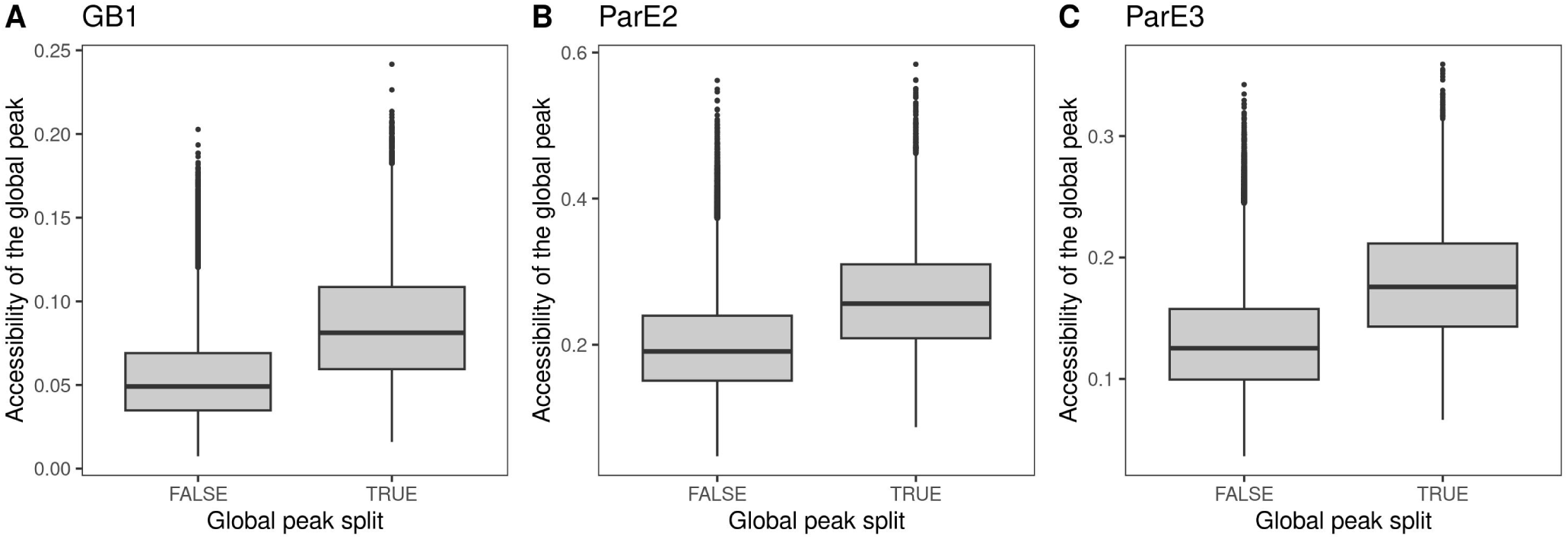
Accessibility of the global peak in relation to whether it forms a single connected region in genotype space for the (A) GB1, (B) ParD-ParE2, and (C) ParD-ParE3 landscapes. The global peak occupies disconnected regions of genotype space when an amino acid in the protein sequence with the highest fitness value is encoded by the split codon block. Data pertain to the 100,000 amino acid permutation codes. Meaning of box-and-whisker plots defined in Fig. S4.

**Figure S6:**
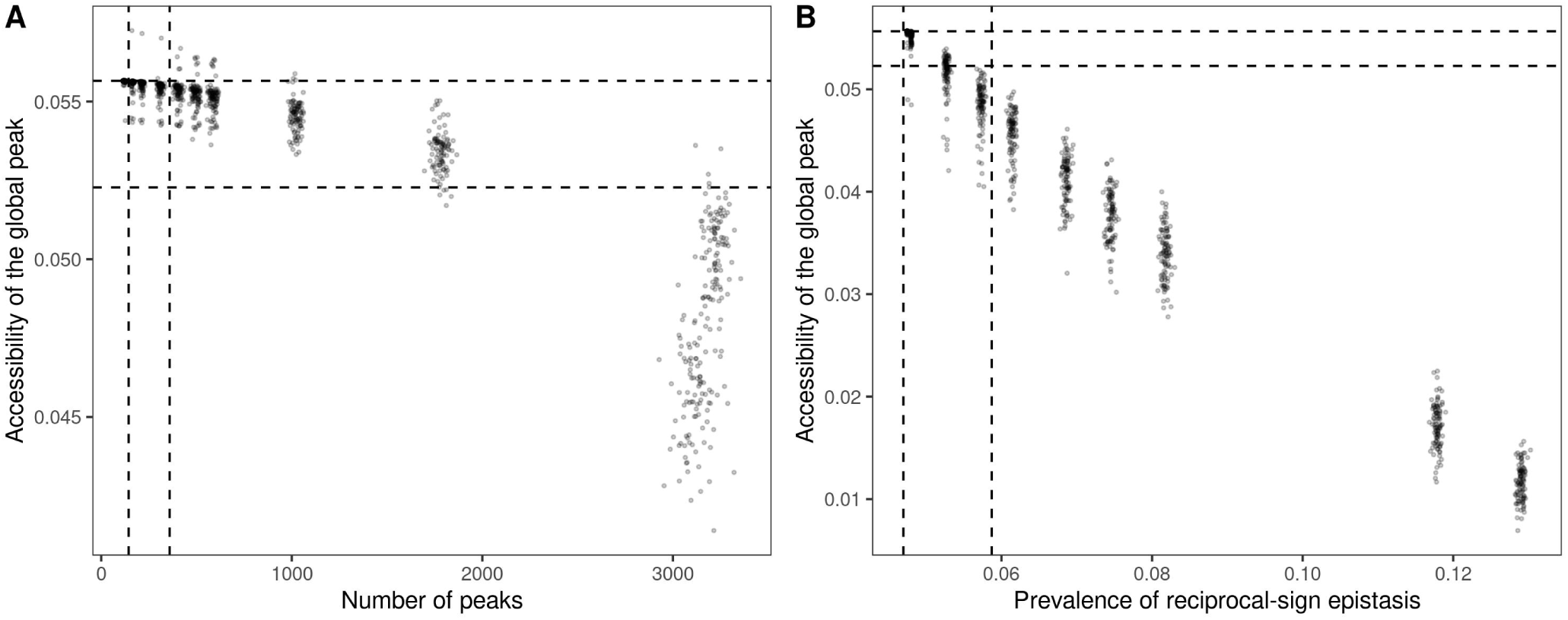
Accessibility of the global peak in relation to two artificially-inflated measures of landscape ruggedness, (A) the number of peaks and (B) the prevalence of reciprocal sign epistasis (Methods). The vertical lines show the 0.01 and 0.99 quantiles in the number of peaks and the prevalence of reciprocal sign epistasis, respectively, for the 100,000 amino acid permutation codes. The horizontal lines show the change in accessibility observed among the 100,000 amino acid permutation codes; the distance between them is the difference between the average accessibility of the global peak in landscapes generated using the 1% most and 1% least robust amino acid permutation codes. For visual clarity, the lines are positioned so that the top line coincides with the mutational accessibility of the global peak in the original landscape. Data pertain to the GB1 landscape under the standard genetic code.

**Figure S7:**
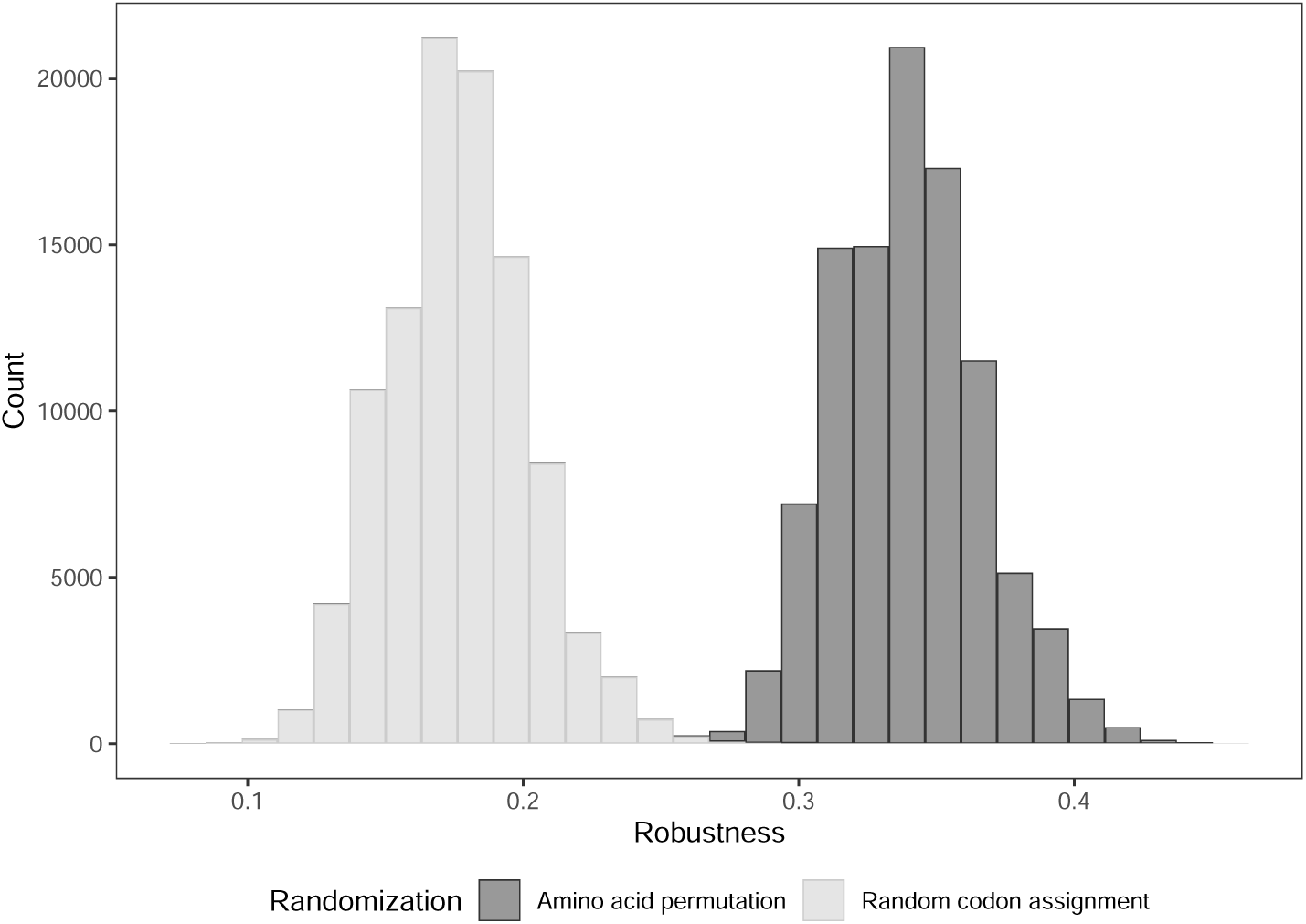
Robustness distributions for genetic codes rewired by amino acid permutation and random codon assignment. Data pertain to 100,000 codes per rewiring scheme.

**Figure S8:**
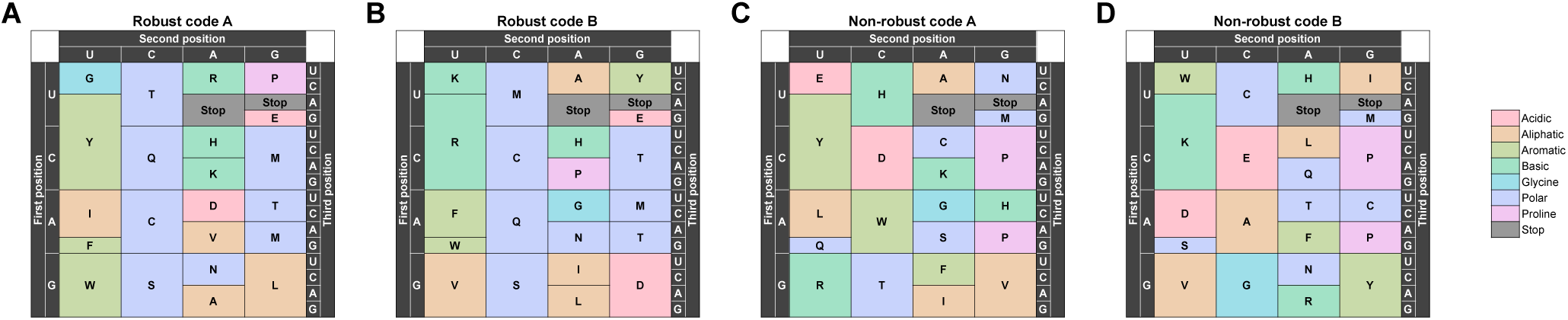
The amino acid permutation codes with the (A, B) highest and (C, D) lowest level of robustness. Codons are colored based on the physicochemical properties of the encoded amino acid, following Pines et al. (2017).

**Figure S9:**
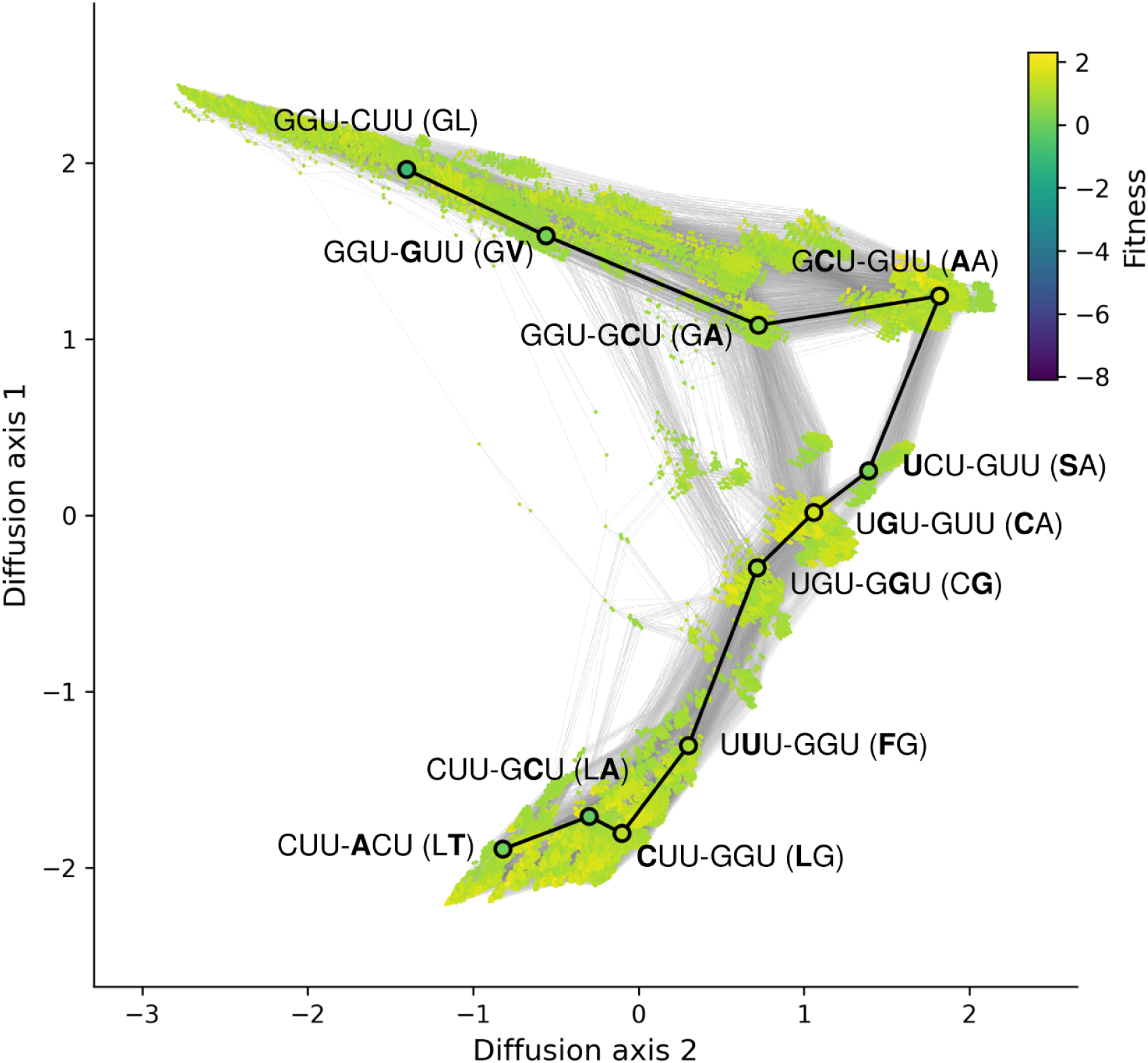
Visualization of the genotype network of high-fitness variants in the GB1 landscape. The highlighted mutational path connects an mRNA sequence for 41G-54L to an mRNA sequence for 41L-54T via a series of intermediate mRNA sequences that are also part of the genotype network (i.e. that are among the 1% most fit sequences). Each highlighted edge corresponds to a non-synonymous point mutation, see Fig. 4 for further information on the layout. The modified nucleotide and amino acid are shown in bold at each step in the path. Color bar indicates protein fitness

**Figure S10:**
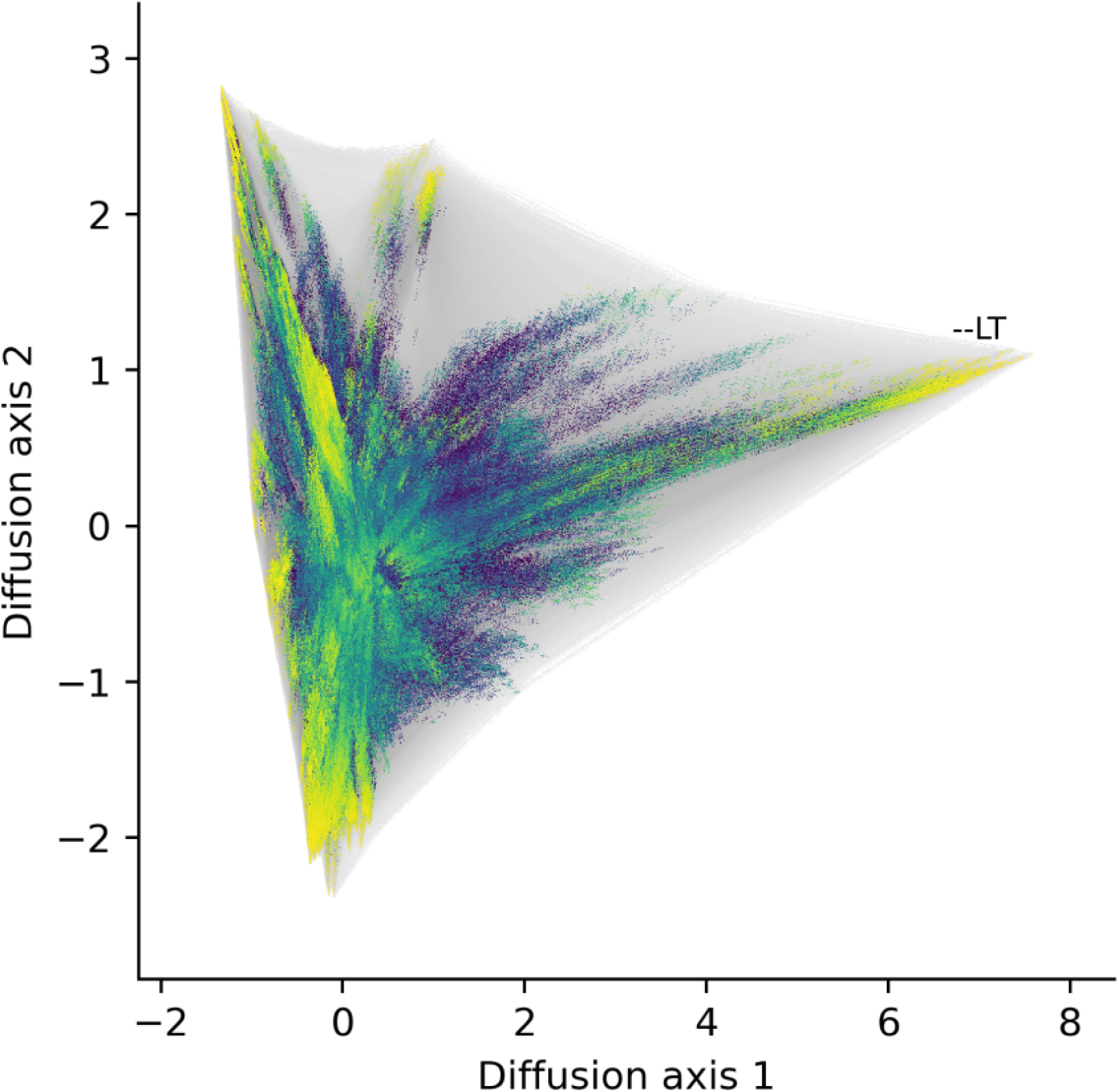
Visualization of the GB1 landscape under Robust Code B. The cluster of 41L-54T variants are indicated, which are separated from the rest of the landscape along Diffusion Axis 1. See Fig. 4 for further information on the layout, as well as the meaning of vertices and edges, and see Fig. 4B for an additional visualization of this landscape along Diffusion Axis 3.

**Figure S11:**
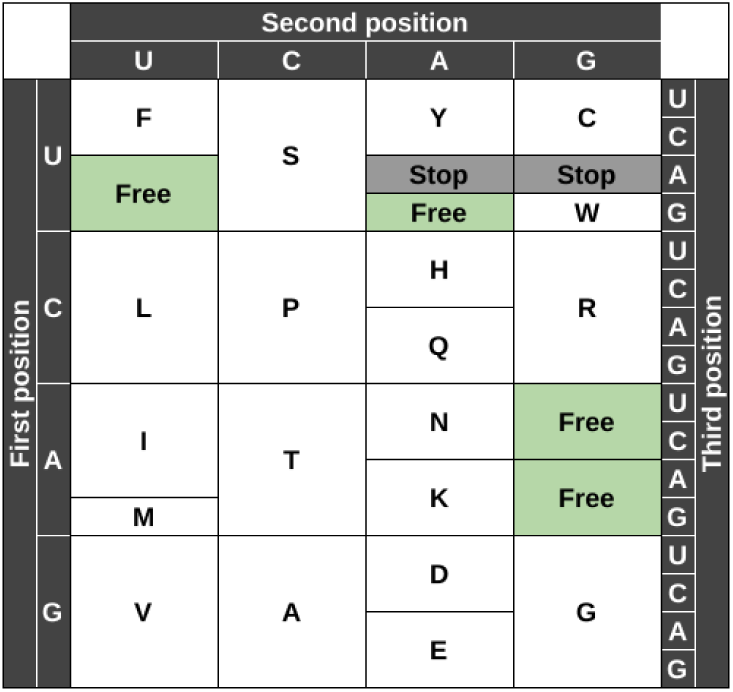
The codon table of the 57-codon *E. coli* genome (Ostrov et al., 2016), highlighting the four codon blocks that have been freed for reassignment.

**Figure S12:**
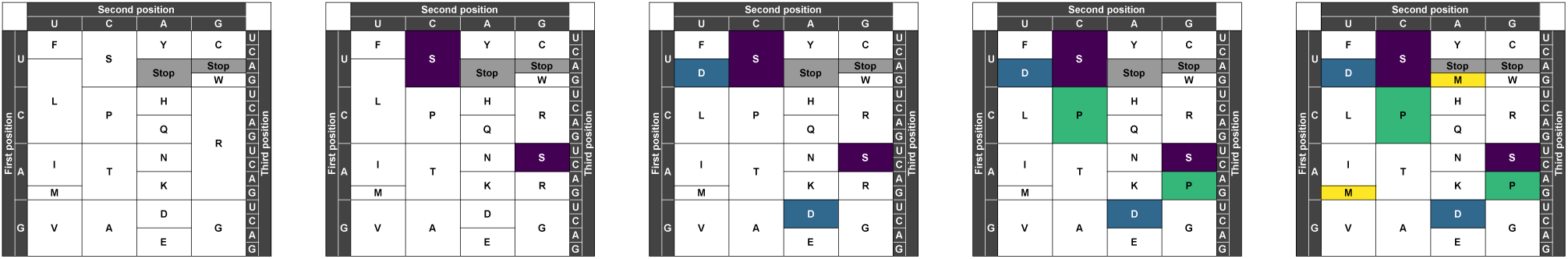
Examples of Ostrov codes with (left) 0 to (right) 4 split codon blocks, which are highlighted in colors.

**Figure S13:**
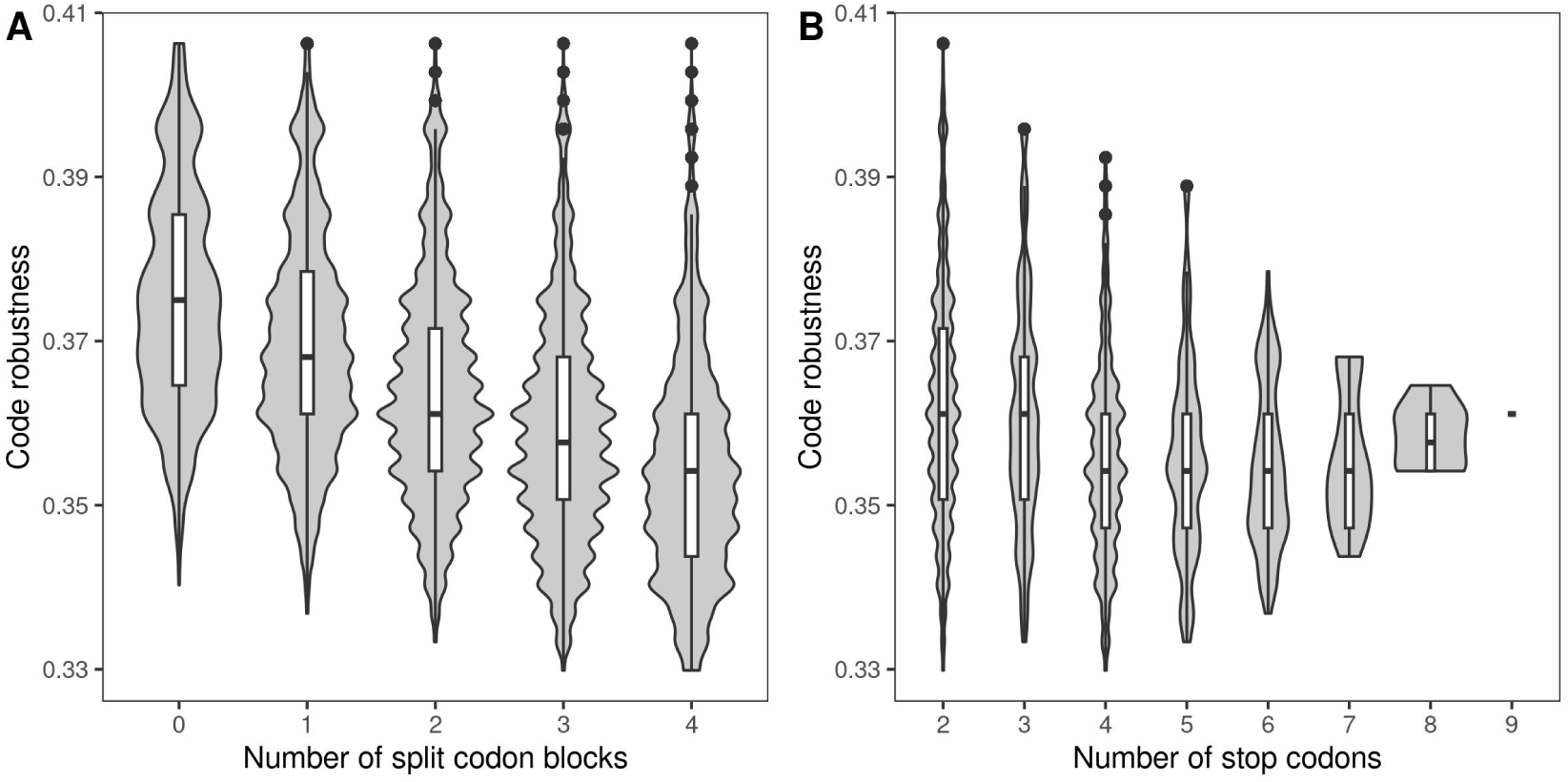
Violin plots of code robustness in relation to (A) the number of split codon blocks and (B) the number of stop codons in the 194,481 Ostrov codes. The violin plots show the distribution and the box-and-whisker plots the median, 25th and 75th percentile of code robustness (see Fig. S4 for details).

**Figure S14:**
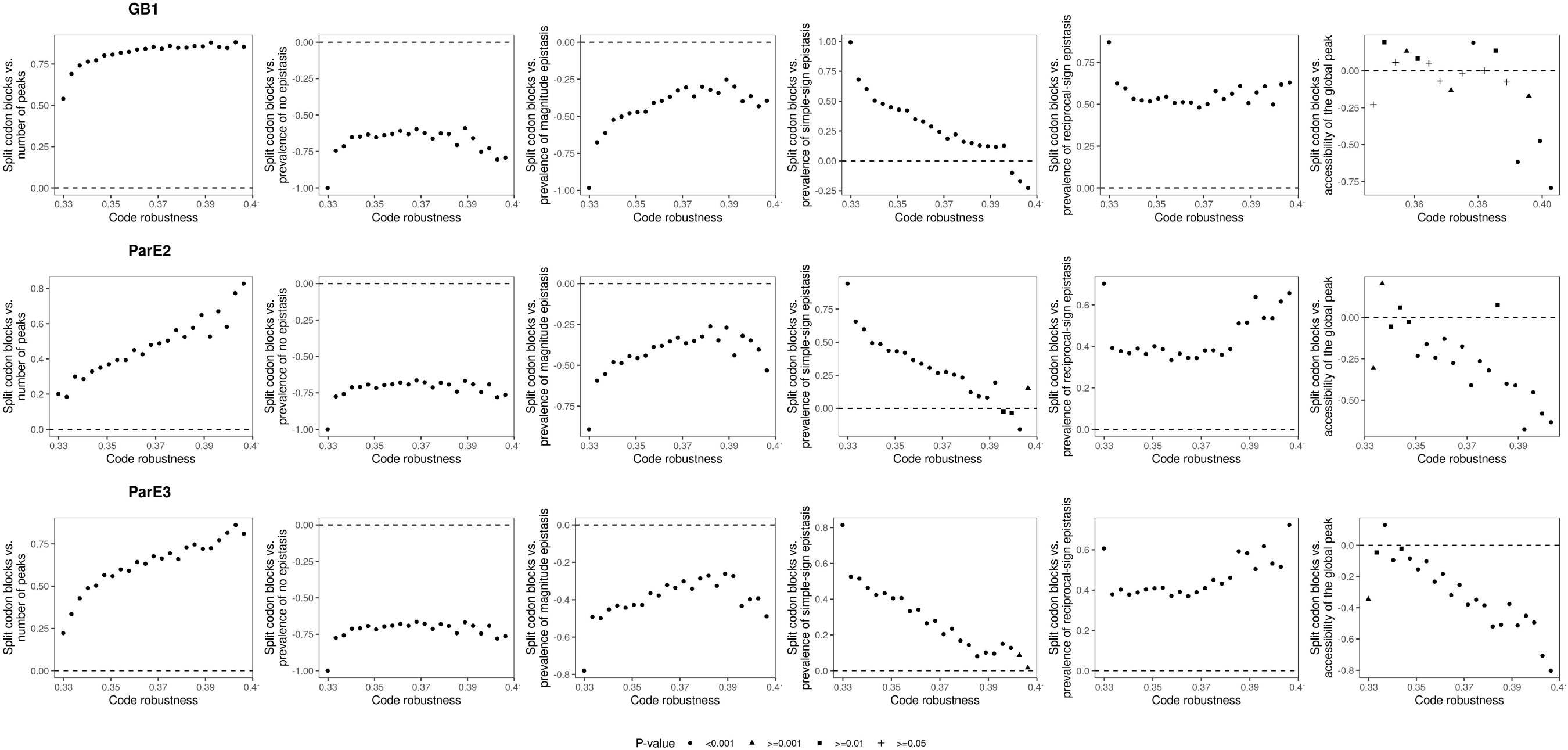
Pearson’s correlation between various measures of landscape ruggedness and the number of split codon blocks, for Ostrov codes with a given value of robustness. The shape of the points denotes the p-value of the correlation coefficient, corrected for testing multiple hypotheses (legend). The magnitude, simple-sign, and reciprocal-sign epistasis results are based on prevalence of a given type of epistasis relative to all epistatic squares. Data for global peak accessibility are based on the subset of Ostrov codes that preserve the size of the global peak and the global peak occupies a single connected region in genotype space. Dashed horizontal lines indicates no correlation.

**Figure S15:**
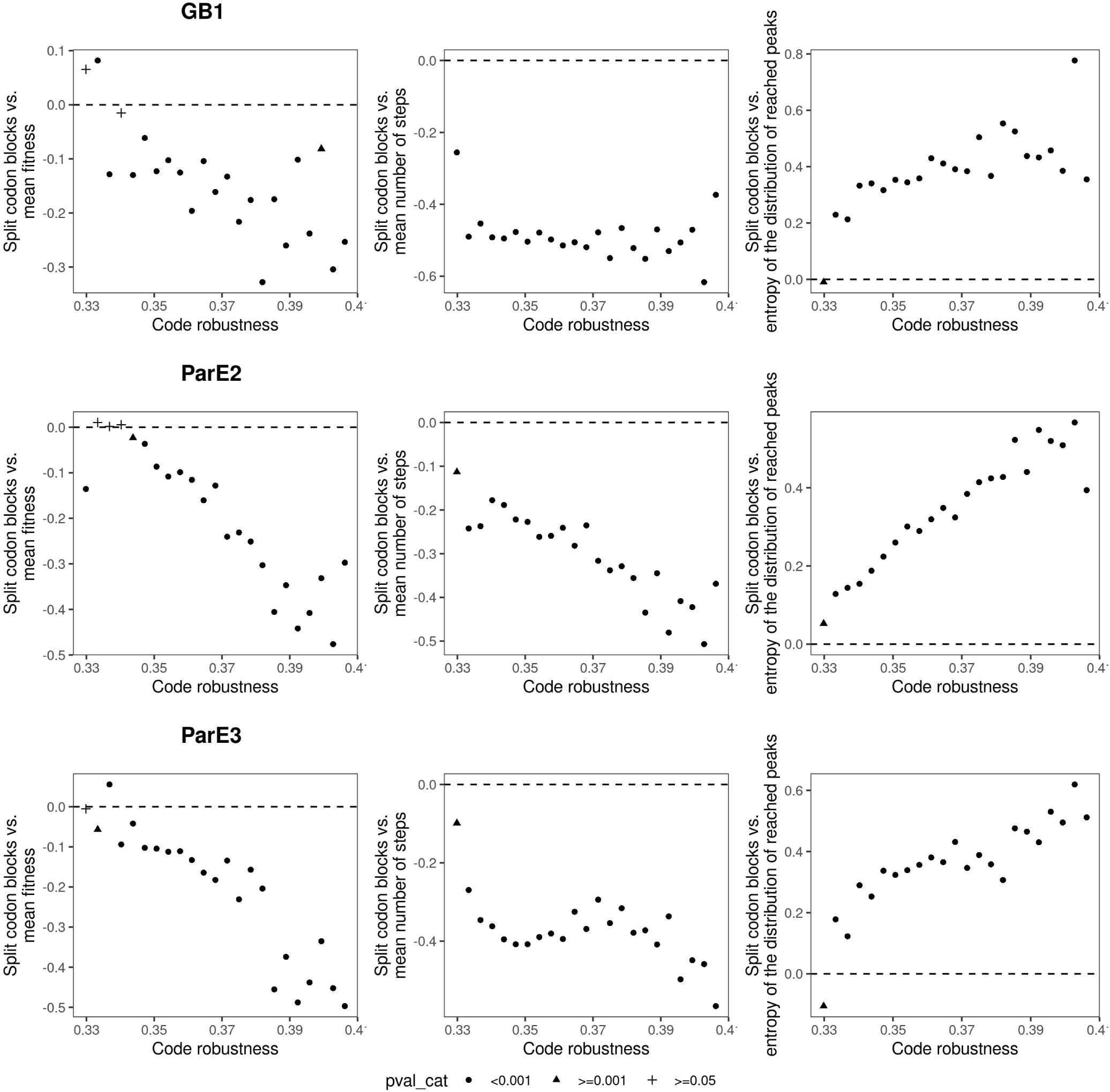
Correlation between various outcomes of the greedy adaptive walks and the number of split codon blocks, for Ostrov codes with a given value of robustness. The shape of the points denotes the p-value of the correlation coefficient, corrected for testing multiple hypotheses (legend). Dashed horizontal lines indicates no correlation.

**Figure S16:**
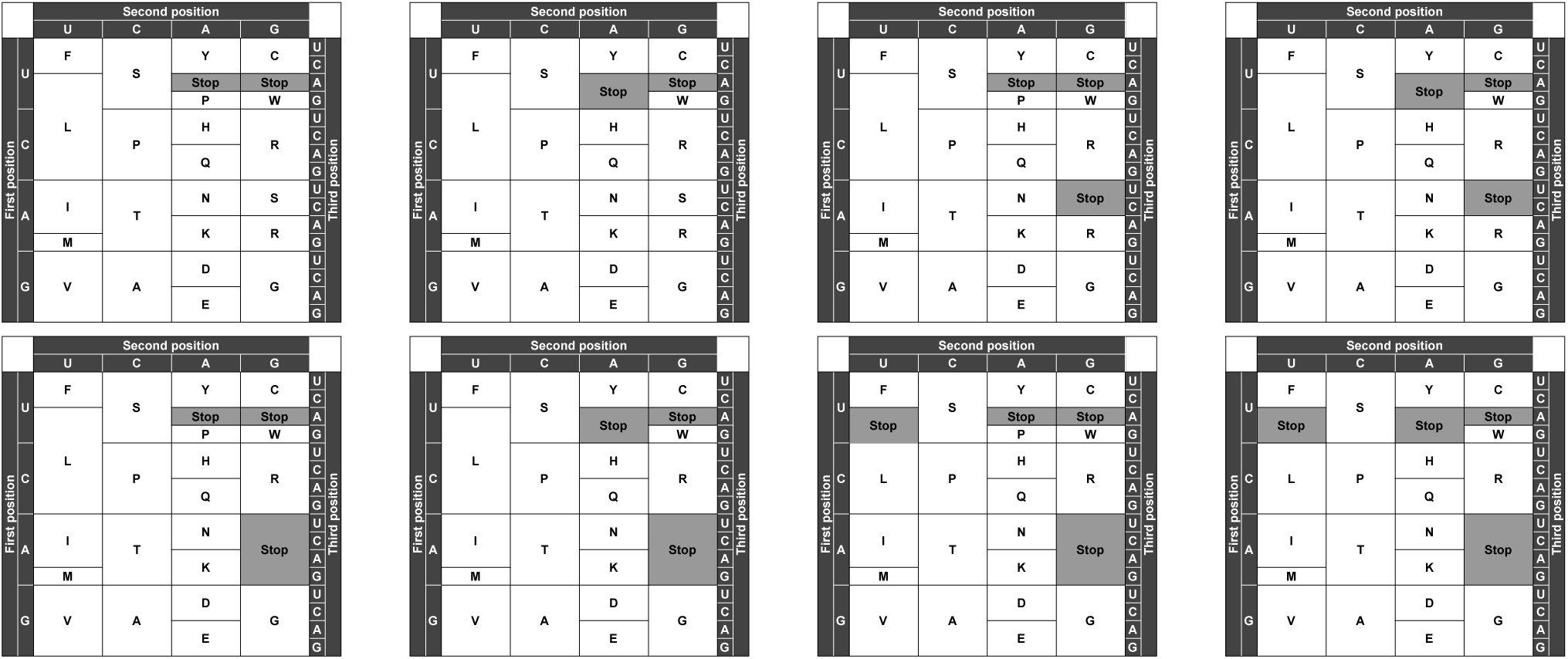
Examples of Ostrov codes with (top left) 2 to (bottom right) 9 stop codons, which are highlighted in grey.

**Figure S17:**
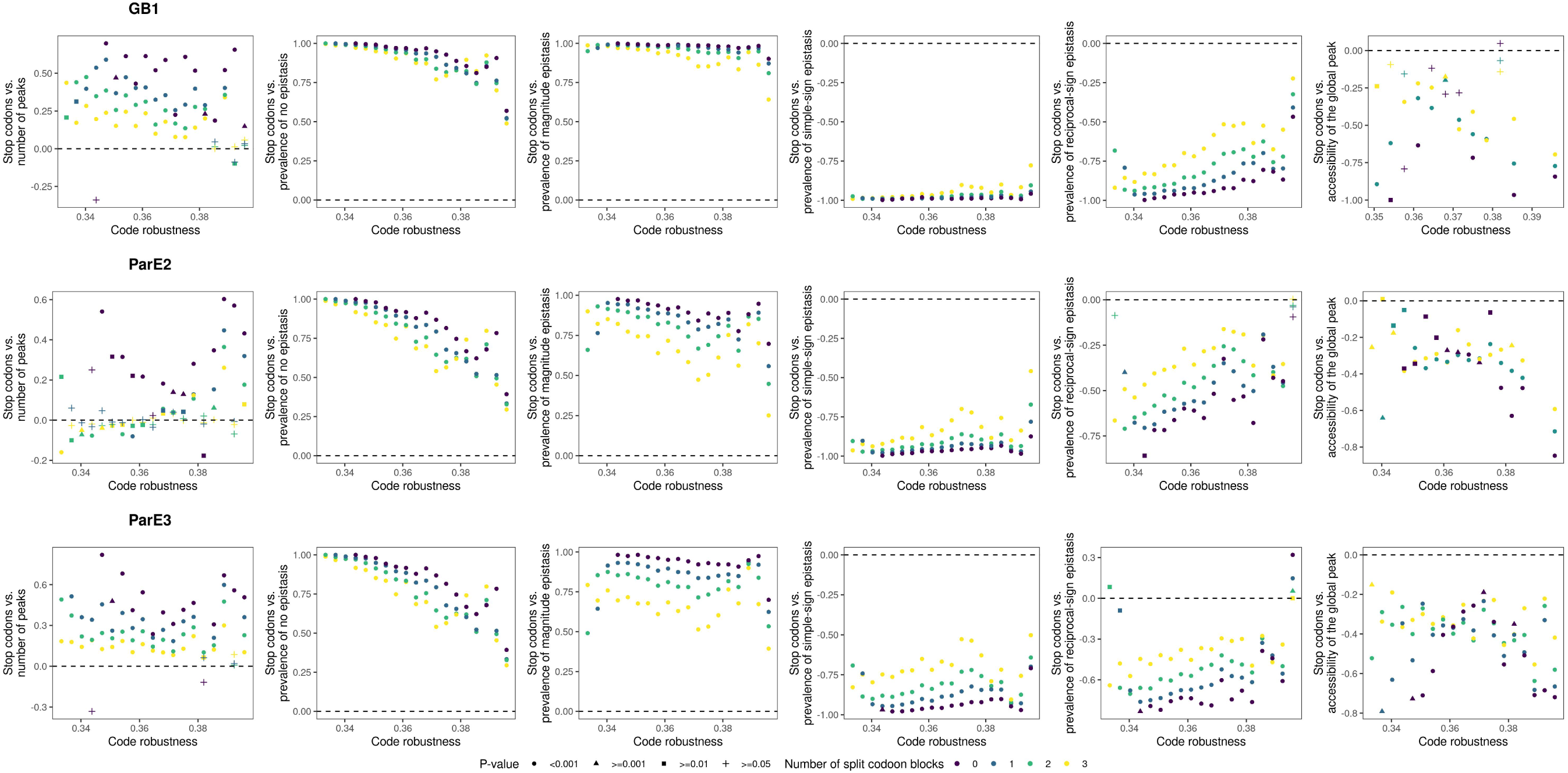
Correlation between various measures of landscape ruggedness and the number of stop codons, for Ostrov codes with a given value of robustness. The shape of the points denotes the p-value of the correlation coefficient, corrected for testing multiple hypotheses, and the color of the points denotes the number of split codon blocks (legend). The magnitude, simple-sign, and reciprocal-sign epistasis results are based on prevalence of a given type of epistasis relative to all epistatic squares. Data for global peak accessibility are based on the subset of Ostrov codes that preserve the size of the global peak and the global peak occupies a single connected region in genotype space. Dashed horizontal lines indicates no correlation. See Supp. Note S1 for further details regarding epistasis.

**Figure S18:**
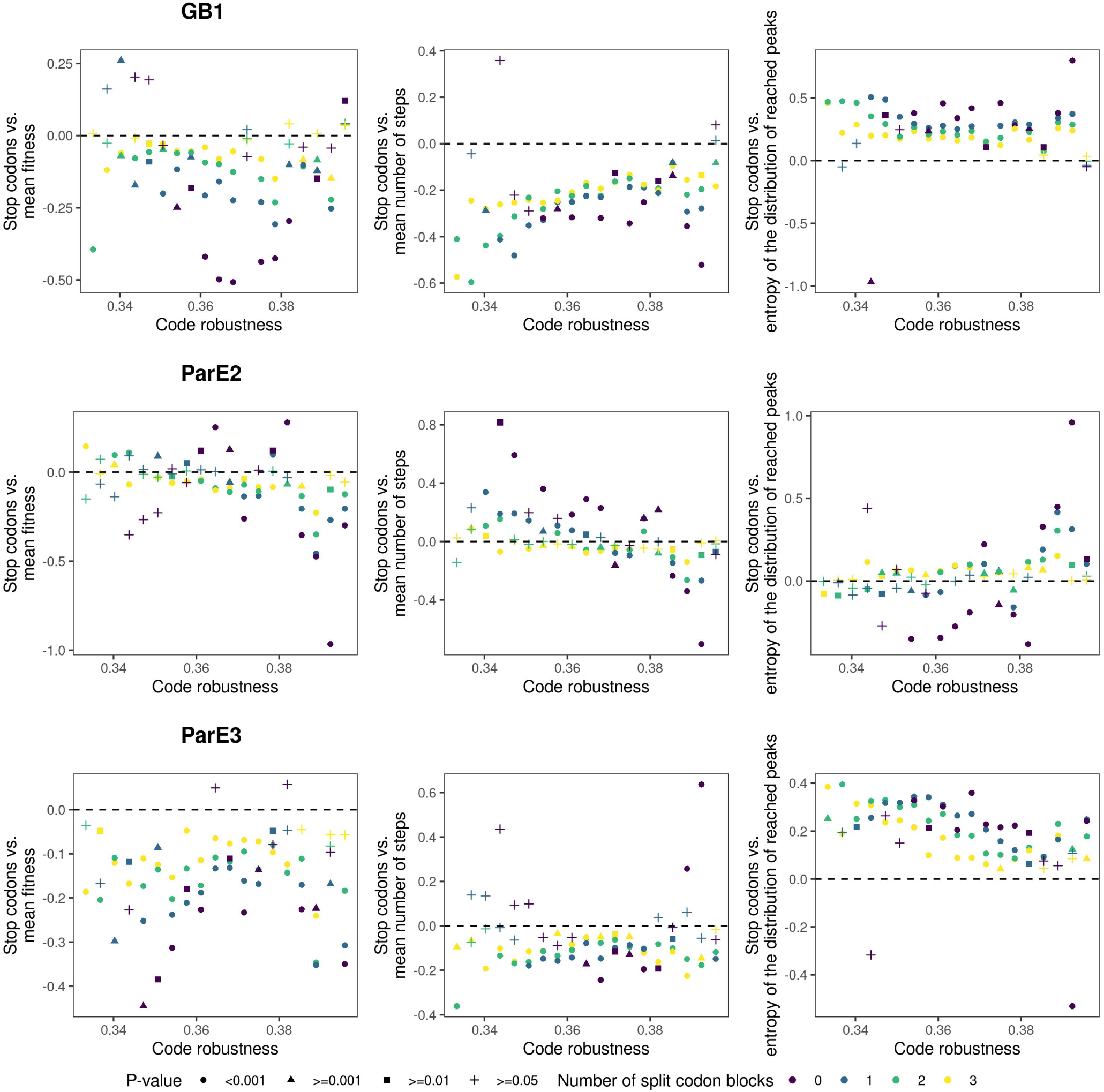
Correlation between various outcomes of the greedy adaptive walks and the number of stop codons, for Ostrov codes with a given value of robustness. The shape of the points denotes the p-value of the correlation coefficient, corrected for testing multiple hypotheses, and the color of the points denotes the number of split codon blocks (legend).

**Figure S19:**
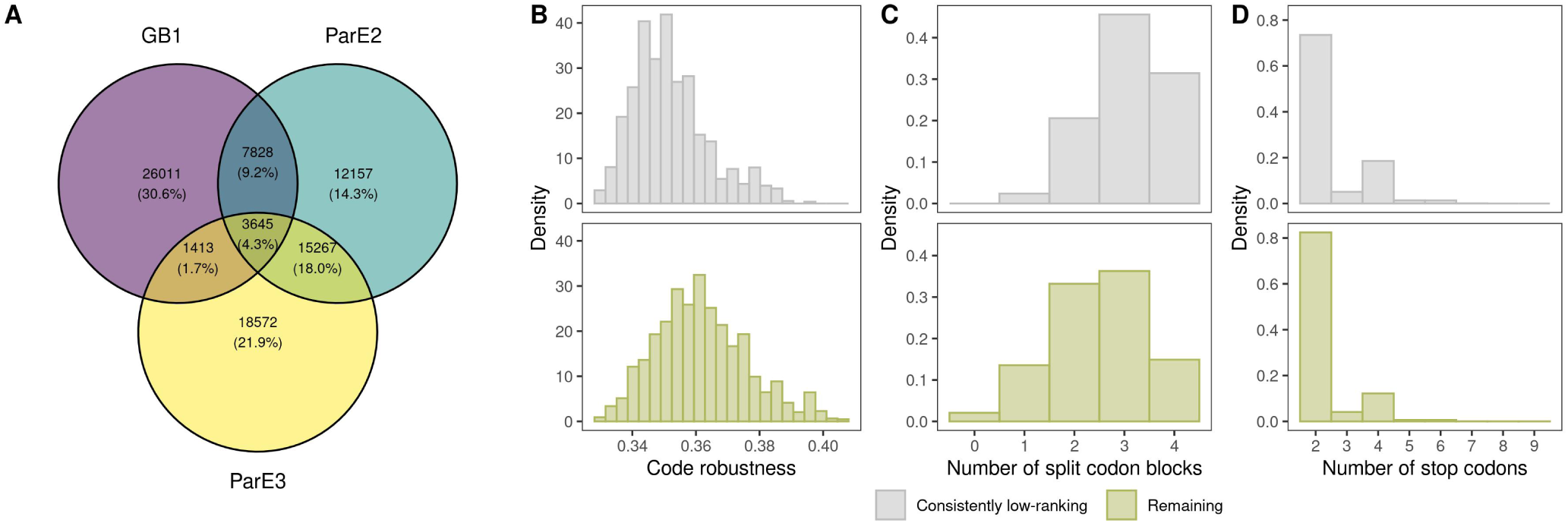
Design principles for diminishing evolvability. (A) Venn diagram of the bottom 20% of Ostrov codes, ranked according to mean fitness reached in the evolutionary simulations, for each of the three data sets. (B)-(D) Comparison of the properties of the 3645 consistently low-ranking codes (three-way intersection in (A)) with the remaining 190,836 codes, in terms of (B) code robustness, (C) number of split codon blocks, and (D) number of stop codons.

**Table S1:**
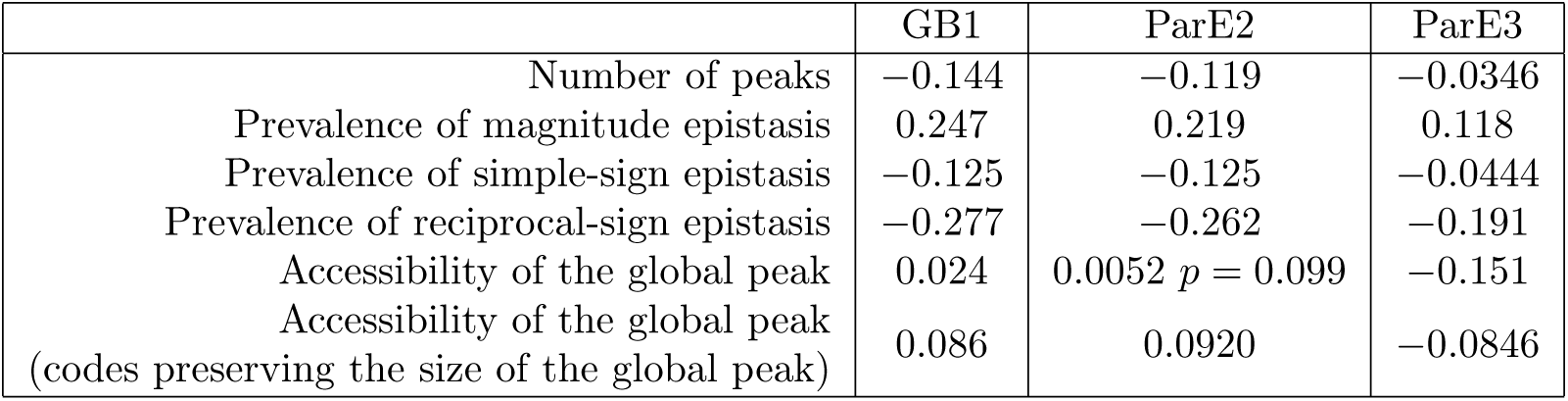
Correlation of various measures of landscape ruggedness with code robustness for amino acid permutation codes. All correlations are statistically significant, unless stated otherwise. Results for the prevalence of no epistasis are not shown, because all amino acid permutation codes have the same proportion of squares with no epistasis: As the fitness values are real numbers, only squares that involve at least two synonymous mutations exhibit no epistasis, and because all 100,000 amino acid permutation codes have the same block structure, the prevalence of such squares is the same for all of them.

**Table S2:** Proportion of the amino acid permutation codes with lower or equal value of a given characteristic, as compared to the standard genetic code. For accessibility of the global peak and the greedy and weak mutation adaptive walks, two proportions are shown: the first one is the proportion of such codes in the whole data set of 100,000 amino acid permutation codes, the second is based on the subset of codes that preserve the size of the global peak and under which the global peak forms a single connected region in the genotype space.

|  |  | GB1 | ParE2 | ParE3 |
| --- | --- | --- | --- | --- |
| Ruggedness | Number of peaks | 0.00037 | 0.326 | 0.570 |
|  | Prevalence of magnitude epistasis | 0.975 | 0.664 | 0.341 |
|  | Prevalence of simple-sign epistasis | 0.123 | 0.548 | 0.604 |
|  | Prevalence of reciprocal-sign epistasis | 0.0169 | 0.189 | 0.676 |
|  | Accessibility of the global peak | 0.538; 0.864 | 0.609; 0.610 | 0.117; 0.643 |
| Greedy walks | Mean fitness | 0.812; 0.857 | 0.961; 0.970 | 0.647; 0.864 |
|  | Mean number of steps | 0.936; 0.903 | 0.993; 0.995 | 0.928; 0.844 |
|  | Entropy of the distribution of reached peaks | 0.00304; 0.0024 | 0.00034; 0.00041 | 0.189; 0.084 |
| Weak mutation walks | Mean fitness after 500 mutations, $N = 10$ | 0.781; 0.839 | 0.756; 0.749 | 0.447; 0.807 |
| | Mean fitness after 500 mutations, $N = 100$ | 0.832; 0.884 | 0.795; 0.802 | 0.385; 0.748 |
| | Mean fitness after 500 mutations, $N = 10,000$ | 0.842; 0.893 | 0.797; 0.806 | 0.404; 0.775 |
| | Mean fitness after 500 mutations, $N = 1,000,000$ | 0.842; 0.894 | 0.796; 0.805 | 0.404; 0.775 |

**Table S3:** Correlation of various measures of landscape ruggedness with code robustness for random codon assignment codes. All correlations are statistically significant, unless stated otherwise. Unlike for the amino acid permutation codes, the random codon assignment codes differ in the proportion of mutations that are synonymous, and hence also differ in the prevalence of squares showing no epistasis. However, the observed correlation with code robustness is exactly the same for all three data sets because prevalence of squares showing no epistasis is determined by the proportion of synonymous mutations in the genetic code.

|  | GB1 | ParE2 | ParE3 |
| --- | --- | --- | --- |
| Number of peaks | −0.387 | −0.392 | −0.329 |
| Prevalence of no epistasis | 0.371 | 0.371 | 0.371 |
| Prevalence of magnitude epistasis | −0.160 | 0.133 | −0.038 |
| Prevalence of simple-sign epistasis | −0.417 | −0.393 | −0.249 |
| Prevalence of reciprocal-sign epistasis | −0.476 | −0.466 | −0.401 |
| Accessibility of the global peak | −0.002, $p = 0.535$ | −0.053 | −0.228 |
| Accessibility of the global peak<br>(codes preserving the size of the global peak) | 0.149 | 0.134 | 0.024, $p = 0.190$ |

**Table S4:** Amino acid properties that are significantly correlated with landscape ruggedness. For each ruggedness measure, the total number of statistically significant properties is shown (number of positively/negatively correlated properties in parentheses). These properties are then broken down into four categories, for which the number of statistically significant properties is shown, along with the p-value of a one-tailed binomial test for over-abundance of the given category among the significantly correlated properties. Statistically significant p-values (*<* 0.05, **in bold**) mean that the category is over-represented among the significantly correlated properties, compared to the null expectation. The proportions of the categories in the database are 26.2% alpha and turn propensity, 8.0% beta propensity, 39.2% hydrophobicity, and 26.6% other.

|  | Total<br>(+/- corr.) | Alpha and turn propensity |  | Beta propensity |  | Hydrophobicity |  | Other |  |
| --- | --- | --- | --- | --- | --- | --- | --- | --- | --- |
|  |  | Observed | Enrichment P-value | Observed | Enrichment P-value | Observed | Enrichment P-value | Observed | Enrichment P-value |
| **GB1** |  |  |  |  |  |  |  |  |  |
| Number of peaks | 171 (169/2) | 14 | 1 | 23 | **0.00937** | 127 | $< 2.2 \cdot 10^{-16}$ | 7 | 1 |
| Magnitude epistasis | 45 (5/40) | 21 | **0.00251** | 9 | **0.00815** | 12 | 0.973 | 3 | 1.00 |
| Simple-sign epistasis | 0 |  |  |  |  |  |  |  |  |
| Reciprocal-sign epistasis | 118 (111/7) | 50 | $1.03 \cdot 10^{-4}$ | 17 | **0.0123** | 43 | 0.762 | 8 | 1.00 |
| Accessibility of the global peak | 6 (5/1) | 0 | 1 | 0 | 1 | 0 | 1 | 6 | $3.53 \cdot 10^{-4}$ |
| **ParE2** |  |  |  |  |  |  |  |  |  |
| Number of peaks | 10 (5/5) | 3 | 0.511 | 0 | 1 | 2 | 0.949 | 5 | 0.0980 |
| Magnitude epistasis | 62 (11/51) | 34 | $1.66 \cdot 10^{-6}$ | 3 | 0.880 | 10 | 1 | 15 | 0.710 |
| Simple-sign epistasis | 3 (0/3) | 0 | 1 | 0 | 1 | 1 | 0.776 | 2 | 0.174 |
| Reciprocal-sign epistasis | 89 (82/7) | 56 | $4.09 \cdot 10^{-13}$ | 6 | 0.720 | 10 | 1 | 17 | 0.961 |
| Accessibility of the global peak | 1 (0/1) | 1 | 0.262 | 0 | 1 | 0 | 1 | 0 | 1 |
| **ParE3** |  |  |  |  |  |  |  |  |  |
| Number of peaks | 7 (2/5) | 0 | 1 | 0 | 1 | 5 | 0.0891 | 2 | 0.594 |
| Magnitude epistasis | 72 (2/70) | 56 | $< 2.2 \cdot 10^{-16}$ | 4 | 0.834 | 4 | 1 | 8 | 1 |
| Simple-sign epistasis | 48 (44/4) | 36 | $2.41 \cdot 10^{-12}$ | 4 | 0.537 | 3 | 1 | 5 | 0.998 |
| Reciprocal-sign epistasis | 95 (95/0) | 74 | $< 2.2 \cdot 10^{-16}$ | 5 | 0.882 | 4 | 1 | 12 | 1 |
| Accessibility of the global peak | 4 (4/0) | 1 | 0.704 | 1 | 0.282 | 2 | 0.512 | 0 | 1 |

**Table S5:** Correlation of code robustness with the outcomes of greedy adaptive walks for amino acid permutation codes. All correlations are statistically significant, unless specified otherwise (p-values in parentheses).

|  |  | GB1 | ParE2 | ParE3 |
| --- | --- | --- | --- | --- |
| Mean fitness | all codes | 0.107 | 0.121 | $-0.00422$ ($p = 0.182$) |
|  | codes preserving the size of the global peak | 0.130 | 0.183 | 0.092 |
| Mean number of steps | all codes | 0.192 | 0.179 | 0.157 |
|  | codes preserving the size of the global peak | 0.239 | 0.170 | 0.124 |
| Entropy of the distribution of reached peaks | all codes | $-0.147$ | $-0.172$ | $-0.0702$ |
| | codes preserving the size of the global peak | $-0.203$ | $-0.213$ | $-0.123$ |

**Table S6:** Correlation of code robustness with the mean fitness reached after 500 steps of the weak mutation adaptive walks, for different values of population size *N*, in the set of amino acid permutation codes. Correlations are statistically significant unless specified otherwise. Data pertain to the subset of amino acid permutation codes that preserve the size of the global peak and under which the global peak forms a single connected region in the genotype space.

|  | GB1 | ParE2 | ParE3 |
| --- | --- | --- | --- |
| $N = 10$ | 0.120 | 0.124 | 0.050 |
| $N = 100$ | 0.096 | 0.155 | 0.029, $p = 0.018$ |
| $N = 10,000$ | 0.091 | 0.148 | 0.018, $p = 0.128$ |
| $N = 1,000,000$ | 0.091 | 0.148 | 0.018, $p = 0.131$ |

**Table S7:** Correlation of code robustness with the outcomes of greedy adaptive walks for the random codon assignment codes. All correlations are statistically significant. For these codes, the subset of codes the preserve the size of the global peak is not required to also fulfill the condition on the global peak forming a single connected region in the genotype space, because due to the nature of the random codon assignment codes this is almost never the case. The size of the subset is *n* = 4, 584 (GB1), *n* = 16, 032 (ParD-ParE2), *n* = 6, 781 (ParD-ParE3).

|  |  | GB1 | ParE2 | ParE3 |
| --- | --- | --- | --- | --- |
| Mean fitness | all codes | 0.140 | 0.127 | -0.0734 |
|  | codes preserving the size of the global peak | 0.141 | 0.183 | 0.092 |
| Mean number of steps | all codes | 0.418 | 0.259 | 0.266 |
|  | codes preserving the size of the global peak | 0.454 | 0.170 | 0.124 |
| Entropy of the distribution of reached peaks | all codes | -0.071 | -0.072 | 0.0635 |
|  | codes preserving the size of the global peak | -0.109 | -0.213 | -0.123 |

**Table S8:** Correlation of code robustness with the mean fitness reached after 500 steps of weak mutation adaptive walks, for different values of population size *N*, in the set of random codon assignment codes. All correlations are statistically significant unless specified otherwise. All results pertain to the subset of codes that preserve the size of the global peak.

|  | GB1 | ParE2 | ParE3 |
| --- | --- | --- | --- |
| $N = 10$ | 0.101 | 0.187 | $-0.010, p = 0.579$ |
| $N = 100$ | 0.058 | 0.197 | $-0.043, p = 0.018$ |
| $N = 10,000$ | 0.053 | 0.185 | $-0.054, p = 0.0032$ |
| $N = 1,000,000$ | 0.053 | 0.185 | $-0.054, p = 0.0032$ |

**Table S9:**
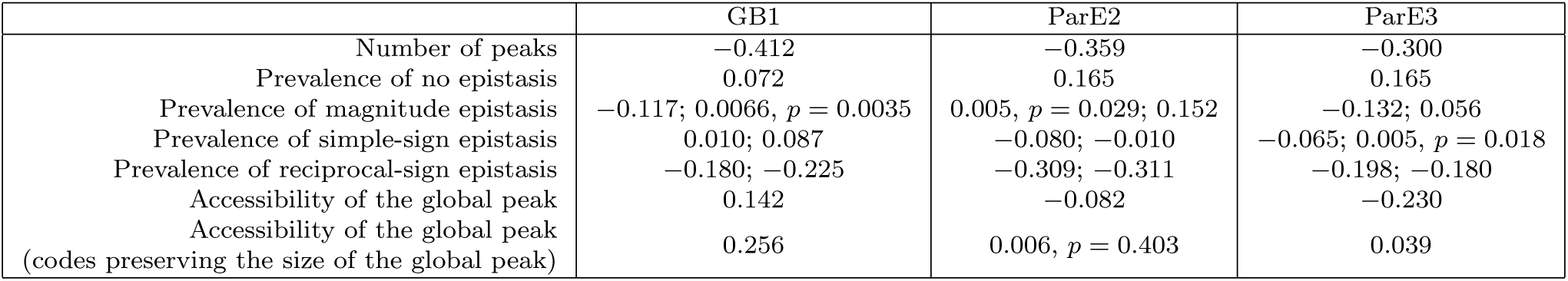
Correlation of various measures of landscape ruggedness with code robustness for the Ostrov codes. All correlations are statistically significant, unless stated otherwise. For magnitude, simple-sign, and reciprocal-sign epistasis, the first number is the correlation between the absolute prevalence of the corresponding type of epistasis and code robustness, while the second one is the correlation between code robustness and the prevalence of a given type of epistasis among epistatic squares only (i.e., in the second case, squares with no epistasis are discarded).

**Table S10:** Correlation of code robustness with the outcomes of greedy adaptive walks for the Ostrov codes. All correlations are statistically significant.

|  |  | GB1 | ParE2 | ParE3 |
| --- | --- | --- | --- | --- |
| Mean fitness | all codes | 0.388 | 0.208 | −0.076 |
|  | codes preserving the size of the global peak | 0.357 | 0.331 | 0.277 |
| Mean number of steps | all codes | 0.359 | 0.266 | 0.261 |
|  | codes preserving the size of the global peak | 0.289 | 0.180 | 0.170 |
| Entropy of the distribution of reached peaks | all codes | −0.264 | −0.151 | 0.040 |
|  | codes preserving the size of the global peak | −0.183 | −0.220 | −0.205 |

**Table S11:** Correlation of code robustness with the mean fitness reached after 500 steps of weak mutation adaptive walks, for different values of population size *N*, in the set of Ostrov codes. All correlations are statistically significant unless stated otherwise. All results pertain to the subset of codes that preserve the size of the global peak and under which the global peak forms a single connected region in the genotype space.

|  | GB1 | ParE2 | ParE3 |
| --- | --- | --- | --- |
| $N = 10$ | 0.300 | 0.295 | -0.141 |
| $N = 100$ | 0.268 | 0.380 | -0.050 |
| $N = 10,000$ | 0.306 | 0.410 | 0.009, $p = 0.0056$ |
| $N = 1,000,000$ | 0.307 | 0.410 | 0.010, $p = 0.0014$ |

**Table S12:** Correlation of various measures of landscape ruggedness with number of split codon blocks for the Ostrov codes. All correlations are statistically significant. The mutational accessibility results pertain to the codes that preserve the size of the global peak and under which the global peak forms a single connected region in the genotype space. For magnitude, simple-sign, and reciprocal-sign epistasis, the first number is the correlation between the absolute prevalence of the corresponding type of epistasis and number of split codon blocks, while the second one is the correlation between number of split codon blocks and the prevalence of a given type of epistasis among epistatic squares only (i.e., in the second case, squares with no epistasis are discarded).

|  | GB1 | ParE2 | ParE3 |
| --- | --- | --- | --- |
| Number of peaks | 0.847 | 0.487 | 0.657 |
| Prevalence of no epistasis | -0.613 | -0.699 | -0.699 |
| Prevalence of magnitude epistasis | 0.681; -0.380 | 0.313; -0.414 | 0.444; -0.364 |
| Prevalence of simple-sign epistasis | 0.452; 0.278 | 0.507; 0.322 | 0.485; 0.297 |
| Prevalence of reciprocal-sign epistasis | 0.614; 0.558 | 0.568; 0.444 | 0.576; 0.434 |
| Accessibility of the global peak | -0.094 | -0.229 | -0.256 |

**Table S13:** Correlation of number of split codon blocks with the outcomes of greedy adaptive walks for the Ostrov codes. All correlations are statistically significant.

|  | GB1 | ParE2 | ParE3 |
| --- | --- | --- | --- |
| Mean fitness | −0.261 | −0.214 | −0.109 |
| Mean number of steps | −0.564 | −0.348 | −0.431 |
| Entropy of the distribution of reached peaks | 0.453 | 0.360 | 0.316 |

**Table S14:** Correlation of number of split codon blocks with the mean fitness reached after 500 steps of random adaptive walks, for different values of population size *N*, for the Ostrov codes. All correlations are statistically significant unless specified otherwise. All results pertain to the subset of codes that preserve the size of the global peak and under which the global peak forms a single connected region in the genotype space.

|  | GB1 | ParE2 | ParE3 |
| --- | --- | --- | --- |
| $N = 10$ | -0.117 | -0.200 | -0.087 |
| $N = 100$ | -0.014, $p = 0.294$ | -0.160 | 0.049 |
| $N = 10,000$ | 0.104 | -0.006, $p = 0.428$ | 0.185 |
| $N = 1,000,000$ | 0.105 | -0.003, $p = 0.703$ | 0.187 |

**Table S15:** Correlation of various measures of landscape ruggedness with number of stop codons, conditioned on the number of split codon blocks, for the Ostrov codes. Results for 4 split codon blocks not shown because all codes with 4 split codon blocks have 2 stop codons. All correlations are statistically significant. The accessibility of the global peak results pertain to the codes that preserve the size of the global peak and under which the global peak forms a single connected region in the genotype space. For magnitude, simple-sign, and reciprocal-sign epistasis, the first number is the correlation between the absolute prevalence of the corresponding type of epistasis and number of stop codons, while the second one is the correlation between number of stop codons and the prevalence of a given type of epistasis among epistatic squares only (i.e., in the second case, squares with no epistasis are discarded). See Supp. Note S1 for the explanation of the epistasis results.

|  | Number of split codon blocks | GB1 | ParE2 | ParE3 |
| --- | --- | --- | --- | --- |
| Number of peaks | 0 | 0.460 | 0.213 | 0.359 |
|  | 1 | 0.390 | 0.103 | 0.304 |
|  | 2 | 0.330 | 0.073 | 0.245 |
|  | 3 | 0.249 | 0.049 | 0.173 |
| Prevalence of no epistasis | 0 | 0.915 | 0.773 | 0.773 |
|  | 1 | 0.922 | 0.789 | 0.789 |
|  | 2 | 0.930 | 0.804 | 0.804 |
|  | 3 | 0.938 | 0.822 | 0.822 |
| Prevalence of magnitude epistasis | 0 | -0.444; 0.986 | 0.102; 0.906 | 0.061; 0.951 |
|  | 1 | -0.429; 0.981 | 0.154; 0.865 | 0.071; 0.893 |
|  | 2 | -0.395; 0.970 | 0.190; 0.806 | 0.088; 0.812 |
|  | 3 | -0.325; 0.939 | 0.189; 0.689 | 0.090; 0.664 |
| Prevalence of simple-sign epistasis | 0 | -0.983; -0.990 | -0.959; -0.969 | -0.953; -0.948 |
|  | 1 | -0.983; -0.987 | -0.952; -0.953 | -0.931; -0.905 |
|  | 2 | -0.982; -0.982 | -0.940; -0.926 | -0.893; -0.842 |
|  | 3 | -0.977; -0.966 | -0.903; -0.860 | -0.805; -0.713 |
| Prevalence of reciprocal-sign epistasis | 0 | -0.928; -0.883 | -0.680; -0.495 | -0.794; -0.688 |
|  | 1 | -0.918; -0.852 | -0.626; -0.446 | -0.753; -0.617 |
|  | 2 | -0.898; -0.802 | -0.555; -0.386 | -0.688; -0.533 |
|  | 3 | -0.843; -0.698 | -0.438; -0.296 | -0.568; -0.409 |
| Accessibility of the global peak | 0 | -0.488 | -0.291 | -0.457 |
|  | 1 | -0.427 | -0.262 | -0.407 |
|  | 2 | -0.330 | -0.205 | -0.341 |
|  | 3 | - | - | -0.250 |

**Table S16:** Correlation of the greedy adaptive walks outcomes with number of stop codons, conditioned on the number of split codon blocks, for the Ostrov codes. Results for 4 split codon blocks not shown because all codes with 4 split codon blocks have 2 stop codons. All correlations are statistically significant.

|  | Number of split codon blocks | GB1 | ParE2 | ParE3 |
| --- | --- | --- | --- | --- |
| Mean fitness | 0 | −0.364 | −0.147 | −0.183 |
|  | 1 | −0.264 | −0.128 | −0.139 |
|  | 2 | −0.196 | −0.107 | −0.104 |
|  | 3 | −0.133 | −0.078 | −0.069 |
| Mean number of steps | 0 | −0.274 | −0.079 | −0.119 |
|  | 1 | −0.281 | −0.074 | −0.131 |
|  | 2 | −0.277 | −0.075 | −0.135 |
|  | 3 | −0.259 | −0.074 | −0.127 |
| Entropy of the distribution of reached peaks | 0 | 0.310 | 0.064 | 0.182 |
|  | 1 | 0.295 | 0.068 | 0.181 |
|  | 2 | 0.269 | 0.071 | 0.165 |
|  | 3 | 0.222 | 0.064 | 0.132 |

**Table S17:** Correlation of the mean fitness reached after 500 steps of a random walk with number of stop codons, conditioned on the number of split codon blocks, for the Ostrov codes. Results for 4 split codon blocks not shown because all codes with 4 split codon blocks have 2 stop codons. All correlations are statistically significant. All results pertain to the subset of codes that preserve the size of the global peak and under which the global peak forms a single connected region in the genotype space.

|  | Number of split codon blocks | GB1 | ParE2 | ParE3 |
| --- | --- | --- | --- | --- |
| $N = 10$ | 0 | -0.255 | -0.088, $p = 0.00268$ | -0.044, $p = 0.037$ |
| | 1 | -0.145 | -0.072 | -0.008, $p = 0.337$ |
| | 2 | -0.084 | -0.050 | 0.011, $p = 0.040$ |
| | 3 | - | - | 0.016, $p = 0.0017$ |
| $N = 100$ | 0 | -0.376 | -0.229 | -0.077, $p = 2.42 \cdot 10^{-4}$ |
|  | 1 | -0.246 | -0.145 | -0.033 |
| | 2 | -0.149 | -0.077 | 0.0011, $p = 0.828$ |
|  | 3 | - | - | 0.021 |
| $N = 10,000$ | 0 | -0.769 | -0.765 | -0.565 |
|  | 1 | -0.631 | -0.626 | -0.495 |
|  | 2 | -0.458 | -0.432 | -0.402 |
|  | 3 | - | - | -0.275 |
| $N = 1,000,000$ | 0 | -0.773 | -0.634 | -0.573 |
|  | 1 | -0.635 | -0.771 | -0.504 |
|  | 2 | -0.463 | -0.440 | -0.411 |
|  | 3 | - | - | -0.282 |

**Table S18:**
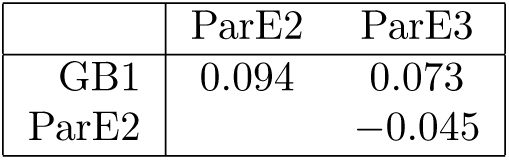
Correlation of the mean fitness reached in the greedy walks by individual Ostrov codes across the three data sets.

**Table S19:**
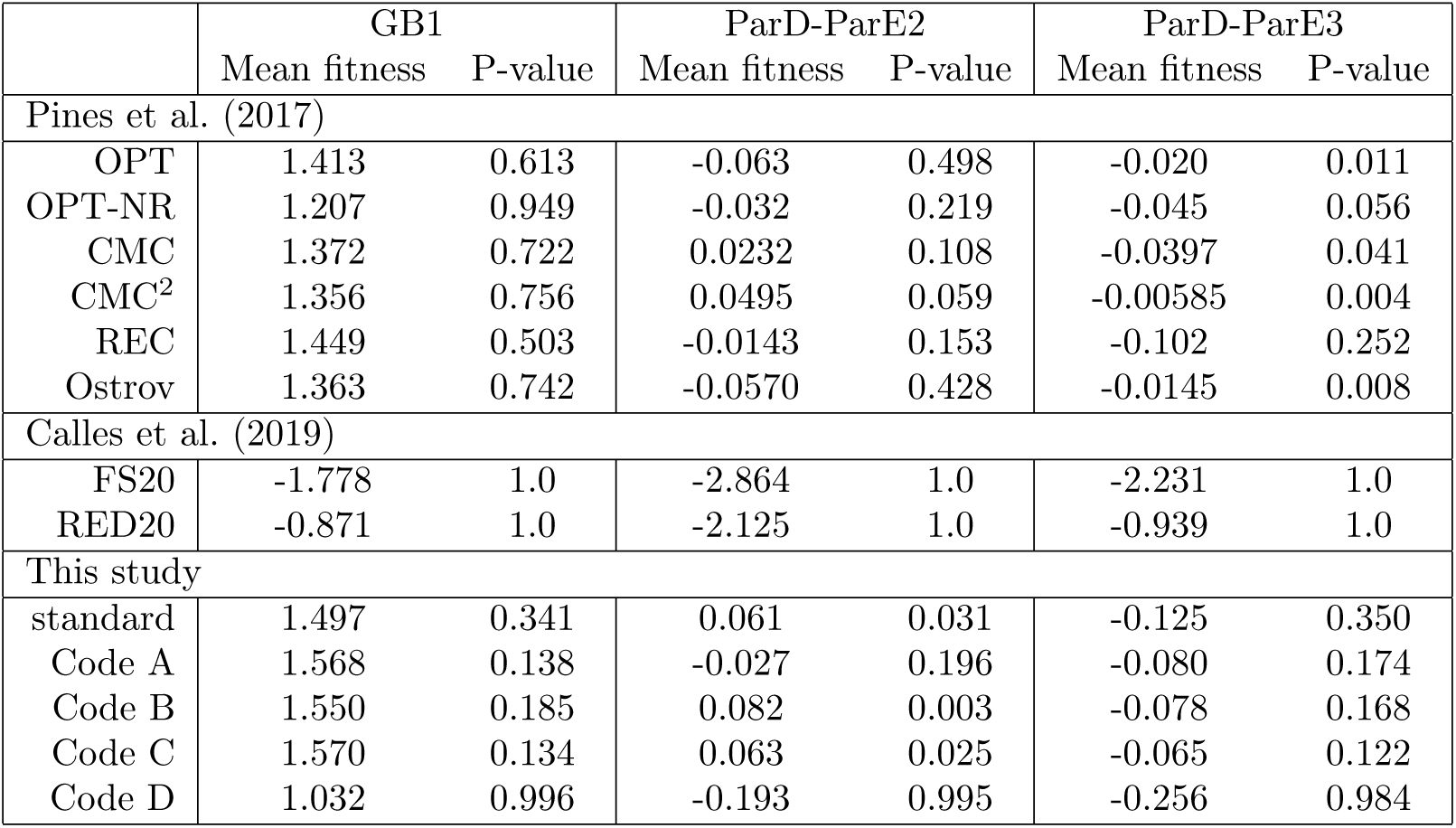
Mean fitness reached in the greedy adaptive walks using the evolvability-promoting codes of Pines et al. (2017), evolvability-diminishing codes of Calles et al. (2019), and the codes identified in this study (codes A-C promote evolvability, code D diminishes). The p-value equals the proportion of the Ostrov codes that have reached the same or higher fitness in the greedy walks.

|  | GB1 |  | ParD-ParE2 |  | ParD-ParE3 |  |
| --- | --- | --- | --- | --- | --- | --- |
|  | Mean fitness | P-value | Mean fitness | P-value | Mean fitness | P-value |
| Pines et al. (2017) |  |  |  |  |  |  |
| OPT | 1.413 | 0.613 | -0.063 | 0.498 | -0.020 | 0.011 |
| OPT-NR | 1.207 | 0.949 | -0.032 | 0.219 | -0.045 | 0.056 |
| CMC | 1.372 | 0.722 | 0.0232 | 0.108 | -0.0397 | 0.041 |
| CMC<sup>2</sup> | 1.356 | 0.756 | 0.0495 | 0.059 | -0.00585 | 0.004 |
| REC | 1.449 | 0.503 | -0.0143 | 0.153 | -0.102 | 0.252 |
| Ostrov | 1.363 | 0.742 | -0.0570 | 0.428 | -0.0145 | 0.008 |
| Calles et al. (2019) |  |  |  |  |  |  |
| FS20 | -1.778 | 1.0 | -2.864 | 1.0 | -2.231 | 1.0 |
| RED20 | -0.871 | 1.0 | -2.125 | 1.0 | -0.939 | 1.0 |
| This study |  |  |  |  |  |  |
| standard | 1.497 | 0.341 | 0.061 | 0.031 | -0.125 | 0.350 |
| Code A | 1.568 | 0.138 | -0.027 | 0.196 | -0.080 | 0.174 |
| Code B | 1.550 | 0.185 | 0.082 | 0.003 | -0.078 | 0.168 |
| Code C | 1.570 | 0.134 | 0.063 | 0.025 | -0.065 | 0.122 |
| Code D | 1.032 | 0.996 | -0.193 | 0.995 | -0.256 | 0.984 |

## S1 Epistasis under the Ostrov codes

The Ostrov codes can vary in the number of split codon blocks and the number of stop codons they contain. The effect of increasing the number of split codon blocks on the prevalence of the different forms of epistasis follows intuition: As the number of split codon blocks increases, the number of synonymous mutations decreases, thus decreasing the prevalence of squares with no epistasis. And among the epistatic squares, increasing the number of split codon blocks increases the prevalence of simple and reciprocal sign epistasis (Supp. Tab. S12 and Supp. Fig. S14). However, the effect of increasing the number of stop codons on the different forms of epistasis is more complicated (Supp. Tab. S15 and Supp. Fig. S17). First, we observe a strong positive correlation between the number of stop codons and the prevalence of no epistasis. This is because the more stop codons a genetic code has, the more mRNAs contain at least one stop codon, and hence the more squares that consist entirely of sequences containing stop codons. As we have assigned the same fitness value to all sequences containing stop codons (Methods), these squares will be classified as exhibiting no epistasis. Second, contrary to expectation, we observe a strong positive correlation between the number of stop codons and the prevalence of magnitude epistasis, and a strong negative correlation between the number of stop codons and simple, as well as reciprocal, sign epistasis (Supp. Tab. S15).

To understand these results, we must think of the types of squares that mRNA sequences containing stop codons can be part of. Not considering the trivial squares consisting only of sequences containing at least one stop codon, there are 6 possible configurations, which we depict in Fig. S20. In the following, we will mostly focus on the configurations in Fig. S20A and S20B, and discuss the remaining 4 configurations at the end. Configuration A involves squares where the “wild type” sequence contains one stop codon and one of the mutations changes the stop codon to a sense codon, while the second mutation happens in any of the remaining codons (Fig. S20A). Configuration B involves squares where the wild type sequence contains one stop codon and both mutations change the stop codon to a sense codon (Fig. S20B). In both cases we assume that the double mutant does not contain any stop codons. To understand the influence these squares have on the prevalence of epistasis, we need to know what types of epistasis squares A and B can exhibit and how many of these squares there are.

All the type A squares show magnitude epistasis, regardless of the fitness values of the two sequences without stop codons. How many squares of this type are there? While the exact number will depend on the particular location of the stop codons in the genetic code, we can estimate the number to be roughly

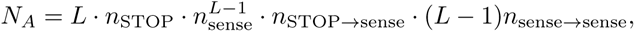

where *L* is the total number of codons in the sequence, *n*_STOP_ is the number of stop codons in the code, *n*_sense_ is the number of sense codons in the code, *n*_STOP_*_→_*_sense_ is the expected number of mutations from a stop codon to a sense codon, and *n*_sense_*_→_*_sense_ is the expected number of mutations from a sense codon to another sense codon. The first three terms quantify the number of sequences containing exactly one stop codon, while the fourth and fifth terms quantify the number of possible mutations that would give rise a type A square.

For the type B squares, the type of epistasis depends on the exact fitness values of the three sequences that do not contain stop codons, which can cause the square to exhibit magnitude, simple sign, or reciprocal sign epistasis. Using reasoning similar to that above, we estimate the number of these squares as roughly

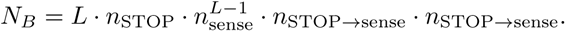

Even without taking into account the fact that the two mutations must happen in two different nucleotide positions, and hence the last two terms are at most 9 *·* 6, *N_A_* is clearly at least (*L −* 1)-times bigger than *N_B_*. In other words, it is much more likely that the second mutation happens in a different codon, than that both mutations happen in the stop codon. As all type A squares exhibit magnitude epistasis, we would thus expect that the prevalence of magnitude epistasis, relative to simple sign and reciprocal sign epistasis, increases as the number of stop codons increases.

Do the remaining possible configurations (Fig. S20C and D) change the result? All squares in Fig. S20C exhibit magnitude epistasis and will thus further increase the prevalence of magnitude epistasis. On the other hand, squares in Fig. S20D, consisting of a wild type and a double mutant that do contain a stop codon and two single mutants that do not, always exhibit reciprocal-sign epistasis. However, in the Ostrov codes the number of such squares is extremely low, due to the fact that the stop codons can be placed in only a small handful of positions; in fact, the maximum number of type D squares in the whole landscape is 2 (see Fig. S20D), so their effect on reciprocal sign epistasis is negligible.

To conclude, contrary to expectation, increasing the number of stop codons increases the prevalence of magnitude epistasis, relative to simple sign and reciprocal sign epistasis, because the number of squares with two neighboring sequences containing stop codons (as in Fig. S20A and C) is much larger than the number of squares where a low-fitness sequence containing a stop codon separates two higher-fitness variants without stop codons (Fig. S11B and D).

**Figure S20:**
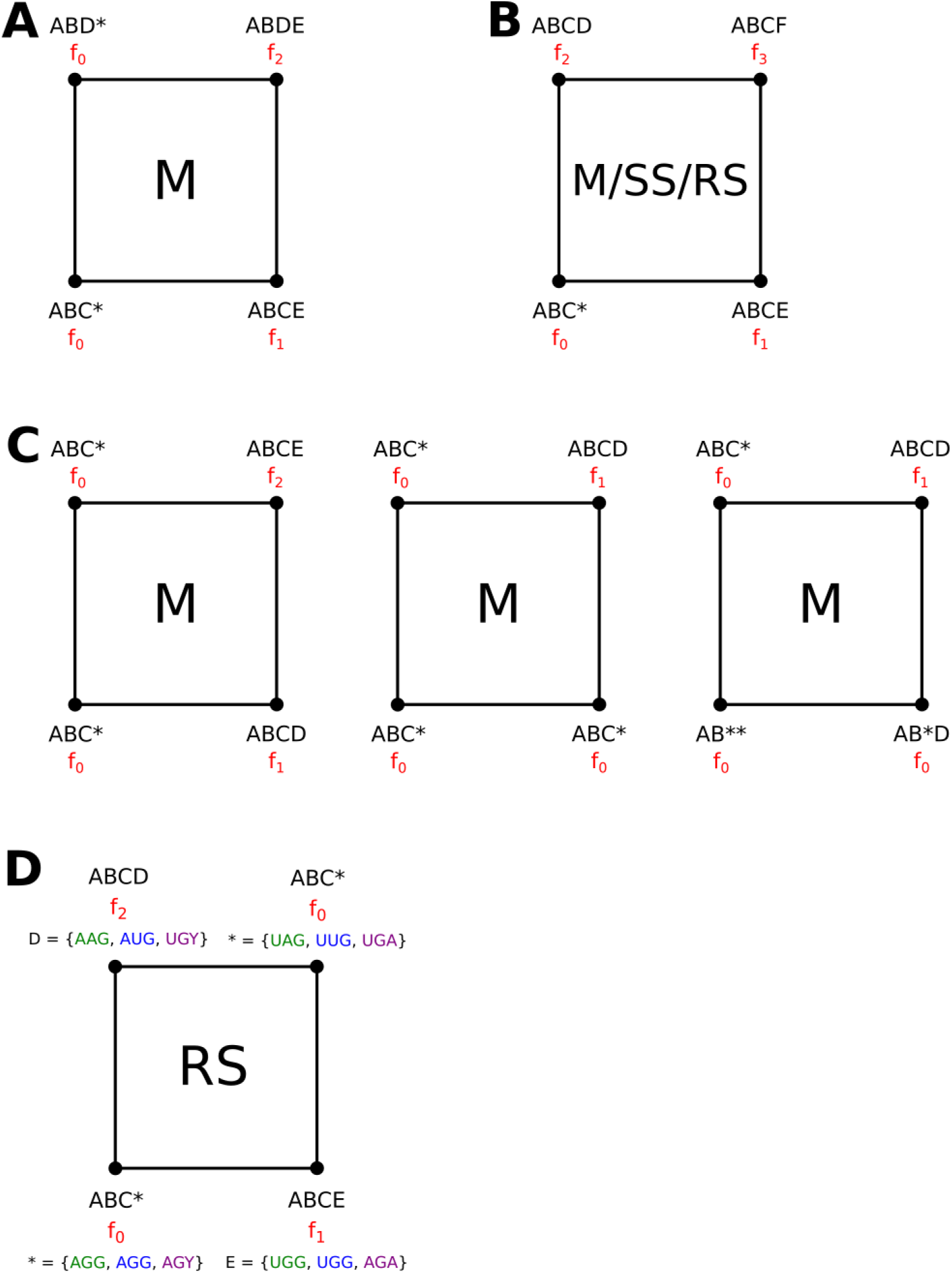
Possible configurations of squares involving at least 1 and at most 3 sequences containing stop codons, for sequences of length *L* = 4, with the stop codon occupying the last position. A, B, C, D, and E denote arbitrary amino acids, not necessarily different from each other. Fitness values are denoted by *f*_0_*, . . ., f*_3_, with *f*_0_ *< f_i_* for *i* = 1, 2, 3, while we assume no particular ordering of *f*_1_, *f*_2_, and *f*_3_. The letters in the middle of the squares denote the possible types of epistasis a given configuration can exhibit; M = magnitude epistasis, SS = simple-sign epistasis, RS = reciprocal-sign epistasis. In D, possible assignments of the last codon, based on the Ostrov codes, are listed in green, blue, and violet; notice that the green and blue assignments are not compatible with the violet one, as AGG and AGA codons need to be assigned the same amino acid (or stop signal), and cannot thus at the same time encode a stop signal (green, blue) and an amino acid (violet). Thus, the maximum number of squares of type D in a landscape caused by an Ostrov code is 2.

